# Glyco-STORM: Nanoscale Mapping of the Cellular Glycosylation Landscape

**DOI:** 10.1101/2025.02.19.639131

**Authors:** Helene Gregoria Schroeter, Steffen Sass, Mike Heilemann, Thomas Kuner, Maja Klevanski

## Abstract

Glycosylation is a crucial biochemical modification of proteins and other biomolecules in cells that generates an exceptional structural and functional diversity. Aberrant glycosylation is implicated in numerous diseases, including neurodegenerative disorders and cancer. Despite its significance, methodological constraints to date have limited the exploration of the nanometer scale spatial arrangement of glycans across entire cells. We developed Glyco-STORM, a super-resolution imaging approach that generates nano-structural maps of cellular glycosylation. Glyco-STORM employs fluorophore-labeled lectins and multiplexed single-molecule super-resolution microscopy, in combination with nanoscale spatial pattern analysis. For example, Glyco-STORM unraveled nanodomains within the endoplasmic reticulum, subdomains along the Golgi axes, and a polarized lysosomal clathrin coat. At synaptic contact sites, mature glycans delineate the synaptic cleft and subsynaptic tubules adjacent to the postsynaptic density. In summary, Glyco-STORM elucidates the spatial arrangement of glycosylation sites from subcellular to molecular levels, revealing the previously obscured glycosylation landscape at nanoscale and establishing a ’spatial glycosylation code’ that provides a unique perspective on cellular organization distinct from traditional protein-centric views.

## Introduction

Glycosylation is one of the major post-translational mechanisms that fine-tunes the function of proteins and lipids in cells. In terms of complexity, the repertoire of mammalian glycans surpasses the proteome^1,2^. Glycans are key-players in many biological processes such as cell signaling, cell-cell adhesion, and cell migration. Their significance extends to various synaptic functions such as neurite outgrowth, formation and maturation of the synapse, and modulation of synaptic transmission and plasticity^3–5^. Carbohydrate modifications are present on major synaptic components, including synaptic vesicle proteins SV2 and Synaptophysin, ionotropic glutamate receptors, calcium channels, and extracellular matrix components in the synaptic cleft. Recent findings highlight significant depolymerization-dependent changes in sialylation of over a hundred synaptic proteins^6^. Neurological abnormalities were found in 80% of all known inherited glycosylation disorders^7^. Furthermore, dysregulation of glycosylation is a well-known phenomenon in cancer cells and tissues^8,9^, which can drive tumor invasion and progression^10^, and glycan-lectin interactions are involved in immunosuppressive mechanisms of tumor cells’ immune escape^11^.

Despite the great biological and medical importance, tools for super-resolution microscopy that map carbohydrates *in situ* and allow their spatial correlation with organelles and proteins remain limited. For super-resolution microscopy, existing glycan labeling primarily employed chemically labeled monosaccharides^12–14^. While metabolic labeling allows very precise analysis of individual sugar moieties, it is impractical for tissue applications, offers limited differentiation of complex carbohydrates, and does not support multiplexing. An alternative method involves carbohydrate-binding proteins, called lectins, which can be used to directly label and map glycans in their cellular context (Supplementary Discussion, section 2). Their low size ranging from approximately 3 to 8 nm is comparable to small antibodies like nanobodies or F(ab’)_2_ fragments. This is advantageous in terms of sample penetration during staining and ensures a shorter linkage error compared to conventional immunohistochemistry employing primary and secondary antibodies (Fig. 1a). So far, direct super-resolution microscopy of glycans *in situ* was limited to a few fluorophore-labeled lectins^15,16^, with WGA being the best-characterized lectin of them. It was employed in *d*STORM to study nuclear pores^15^ and resPAINT^17^ where it stained the glycocalyx.

**Fig. 1.**
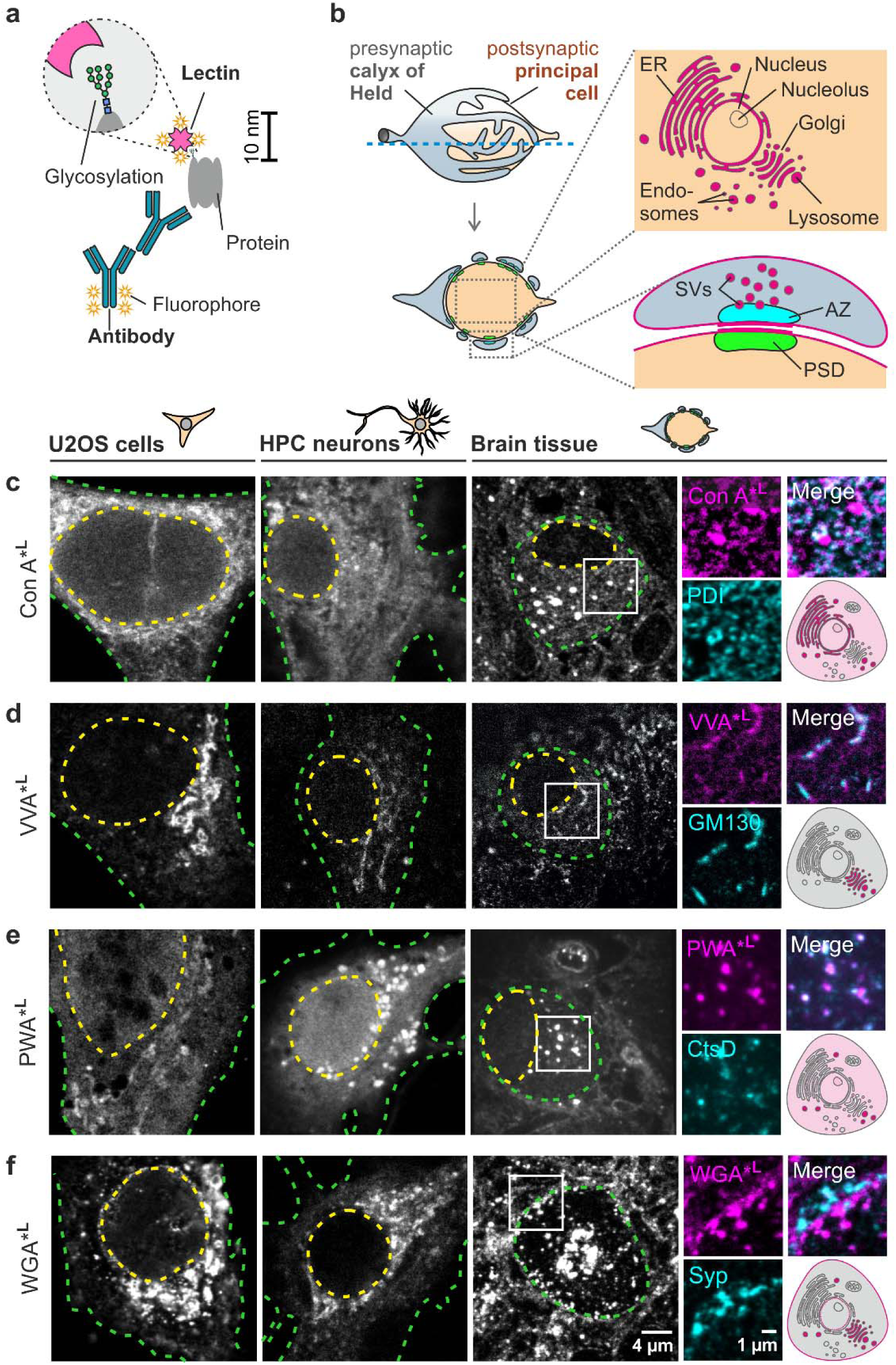
Overview of lectin labeling and confocal microscopy-based screening of selected lectins with diverse labeling patterns. **a**, Fluorophore-conjugated lectins (drawn to scale) have a small linkage error (≤7 nm). b-f, Lectin staining is tested across three biological models: osteosarcoma epithelial cells (U2OS), hippocampal (HPC) neurons, and brain tissue containing giant glutamatergic calyx of Held synapses. **b**, Giant synapse between the calyx of Held and principal cell in semi-thin brain sections. The large soma of the principal cell contains organelles typical for mammalian cells and highlights organelle-specific glycosylation (magenta). Postsynaptic densities (PSDs) align with presynaptic active zones (AZs), enabling the study of glycosylation in synaptic membranes and vesicles (SVs). **c-f**, Confocal microscopy-based lectin screening in three biological models. The nucleus and cell border are delineated by a yellow and green dashed line, respectively. For brain sections, different lectins (magenta) were co-stained with antibodies (cyan) for validation.

The limited application of lectins as labels in super-resolution imaging is due to (1) the lack of a systematic characterization of lectins with diverse sugar-specificities for distinct cellular compartments and individual glycosylated molecules, (2) the limited multiplex capability of most super-resolution methods that would allow validation of lectin binding by established markers, and (3) the absence of a universal easy-to-use protocol for super-resolution imaging of lectins. Glyco-STORM overcomes these obstacles. In our approach, we integrate fluorophore-labeled lectins with advanced automated 3D direct stochastic optical reconstruction microscopy (*d*STORM) designed for multiplex experiments^18^. Presenting a universal protocol, we offer a streamlined process for *d*STORM imaging of lectin-labeled glycans combined with conventional immunohistochemistry. Through this approach, we were able to characterize an array of lectins with diverse sugar specificities and showed that lectins specifically bind to distinct organelles and specializations in neuronal and non-neuronal cells, yielding superior resolution comparable to directly labeled primary antibodies and nanobodies. Our comprehensive screening generated a toolbox of 17 lectins that contributed to and will further complement our knowledge on the role of glycosylation in different subcellular areas. In particular, we show that Con A and POL can be specifically used to study how mannosylation is compartmentalized within the endoplasmic reticulum (ER). Numerous lectins, including VVA, VEA, PSA, MAA II, and WGA enable to study the nanostructure of the Golgi apparatus from a ‘glyco-centric’ perspective. Amongst others, PWA and PSA label carbohydrates of pre-lysosomal compartments (PLCs) and lysosomes. We show that PSA and WGA bind synaptic membranes, are enriched in synaptic clefts, and localize to synaptic vesicle clusters, paving new ways of studying synaptic architecture and its implications for neurotransmission. Finally, we provide researchers with a list of recommended lectins and an informative guide for their strategic application in addressing various scientific questions.

Thus, Glyco-STORM brings a novel, ‘glyco-centric’ perspective to cell biology by incorporating spatial glycosylation as a crucial layer of organization, overarching the proteome level.

## Results

### Screening for lectins suitable for spatial glycosylation mapping

To determine carbohydrate-binding candidates suitable for super-resolution analysis and to demonstrate their binding properties across different biological models, we first screened 17 lectins with various predominant binding motifs (Table S1) in three different models (Fig. 1b-f): (1) the osteosarcoma cell line U2OS, representing a non-neuronal model, (2) primary hippocampal (HPC) neurons, and (3) brain tissue sections containing giant glutamatergic synapses (calyces of Held). The defined geometry of this synapse in 400 nm thin cross-sections allows for simultaneous analysis of subcellular lectin signals within the large postsynaptic principal cell and glycosylation of synaptic specializations at the presynaptic calyx and at the periphery of the principal cell (Fig. 1b).

Chemically fixed cells and tissue were incubated with fluorophore-conjugated lectins in staining buffer supplemented with specific ions for optimal lectin binding (Table S2). To pinpoint the lectin binding areas more precisely during the initial screening, cells and tissue were co-stained with antibody markers for various organelles (Table S3-4). Confocal imaging revealed promising lectin candidates (denoted by ‘*^L^’ throughout the figures, e.g., Con A*^L^) that showed enhanced signal in different intra- and peri-cellular regions (Fig. 1c-f and Fig. S1). The reticular signals of Con A and POL partially overlapped with the ER marker PDI (Fig. 1c and Fig. S1a), whereas their vesicular staining patterns showed no such overlap. Multiple lectins, including VVA, WGA, VEA, PSA, MAA II, and WGA, localized at or next to the Golgi apparatus as confirmed by Golgi marker GM130 (Fig. 1d,f and Fig. S1b-f). Con A, PWA, WGA, VEA, PSA, and LCA, stained large vesicular structures (Fig. 1c,e,f and Fig. S1c,d,g), with PWA colocalizing with the pre-lysosomal and lysosomal marker cathepsin D (Fig. 1e). While Golgi-associated staining was consistent among the different cell systems, the abundance and size of vesicular structures varied. Brain tissue sections allowed to identify WGA and PSA as lectin candidates exhibiting the most pronounced staining around synaptic compartments, presumably at plasma membranes (Fig.1f and Fig. S1d). Additionally, multiple lectins (DBA, PWA, VVA, WGA, LCA, PSA, MAA II, UEA I, UDA, LTL) showed abundant homogeneous signal throughout the cytoplasm that could not be mapped to specific organelles –– with a subset of these (DBA, PWA, MAA II, UEA I, UDA, LTL) also showing nuclear staining (Fig.1 and S1). Interestingly, in neuronal tissue, DBA showed widespread staining but was notably absent in the Golgi/TGN area occupied by WGA (Fig. S1i). Some candidates, including HHA, CAL, PNA, and ABA, showed variable performance (Fig. S2 and Supplementary Discussion, section 1.2), and were not included in most super-resolution experiments. Lectins identified as robust binders were further validated using competing carbohydrates (Fig. S3; Table S5, 6; Supplementary Discussion, section 1.1) and subjected to super-resolution microscopy.

### Glyco-STORM: A lectin-based super-resolution imaging approach

To map subcellular and suborganellar distribution of carbohydrates, lectin-stained samples were imaged using maS³TORM^18^, an approach that combines automated 3D dual-channel imaging of two spectrally close fluorophores with automated robot-mediated solution exchange. For universal and concise staining and *d*STORM imaging of fluorophore-labeled lectins and antibodies, we streamlined our staining and imaging workflow (Fig. S4, Methods section ‘Lectin and antibody staining’) by applying lectins alongside with directly labeled primary or secondary antibodies. This involved a transition from PBS- to Tris-based staining and imaging buffers, accommodating the lectins’ need for ions incompatible with PBS. For *d*STORM imaging, we adjusted Tris, MEA, and KOH concentrations to match the performance of AF647 and CF680 fluorophores that we previously achieved with the PBS-based buffers (Fig. S5a,b; for final protocol, see Methods section ‘Multiplexed *d*STORM imaging’). In our multiplexing approach, lectin and antibody signals from prior imaging rounds were effectively removed through photobleaching, which not only bleaches fluorophores but also facilitates photo-unbinding of the fluorophore-conjugated labels^18,19^. In cases when secondary antibodies against host species that were employed in previous rounds had to be re-applied in the upcoming round, chemical elution was additionally employed to achieve effective clearance of antibodies. Lectins were tested on three different sample types: nervous tissue, U2OS cells, and hippocampal neurons (Tables S8-10). Among these, brain tissue was chosen as the primary focus for Glyco-STORM imaging due to its superior accessibility and geometry.

Using nearest neighbor and decorrelation analyses^20,21^, we determined a localization precision of 11.9 ± 2.5 nm and a resolution of 26.3 ± 0.6 nm in lectin-based imaging, *en par* with direct labels (Fig. S5c,d). While these analyses can sense major alterations in quantum yield and blinking dynamics of the fluorophore and the label affinity, a different approach is required to determine label-epitope distances. For example, previously we demonstrated that the glycosylated spike protein of SARS-CoV-2 virions can be stained with Con A (∼7 nm) and compared this to an immunolabeling approach using a sandwich of a primary minibody and a secondary F(ab’)_2_ fragment antibody (12–14 nm in total). Measuring the radius of the labeled virions revealed a reduction by ∼5 nm when using Con A in comparison to the sandwich, demonstrating a smaller linkage error of lectins ^22^.

### Lectins with nuclear and perinuclear binding

To leverage the unique capabilities of lectins for probing glycans at nanoscale resolution using *d*STORM, we selected the nucleus as a first target organelle for two main reasons. Technically, its large size and distinct structure facilitates segmentation and quantification. Biologically, the nucleus presents a rich field of study: while it is known that a multitude of glycosylated molecules are present in and around the nucleus, the identity, distribution, and role of many glycans remain unclear. Indeed, Glyco-STORM imaging revealed lectins that were abundant in the nuclear, nucleolar, and perinuclear areas. MAA II displayed a substantial signal in the Hoechst-JF646-stained nucleus and a more pronounced signal in the nucleolus area, marked by the fibrillarin antibody (Fig. 2a). WGA stained nuclear pores, aligning with previous studies^15,23^, while Con A showed signals in the perinuclear region, corresponding to the perinuclear ER. In addition, WGA and Con A displayed a moderate signal within the nucleus. To evaluate the relative abundance of lectins in the nucleoplasm, we quantified the absolute signal density (Fig. S6) and the density of signal clusters (Fig. 2b,c). Nuclear signals of DBA, UDA, MAA II, WGA, Con A, and VVA significantly exceeded background signal measured at the surface of the glass-bottom dish, indicating that their staining is nucleus-related (Fig. S6; see Supplementary Discussion for details). While DBA, UDA, and MAA II displayed a broad and abundant distribution across the nucleoplasm, LTL, WGA, VVA, and Con A were confined to sparse non-homogeneously distributed clusters (Fig. 2b,c). Thus, Glyco-STORM identified multiple lectins with diverse sugar specificities (Fig. 2c) that bind various nuclear and peri-nuclear regions, yielding both abundant and confined signals. These lectins provide a starting point to investigating glycosylation in the nucleoplasm, the import of glycosylated proteins, and function of glycans in the nucleus.

**Fig. 2.**
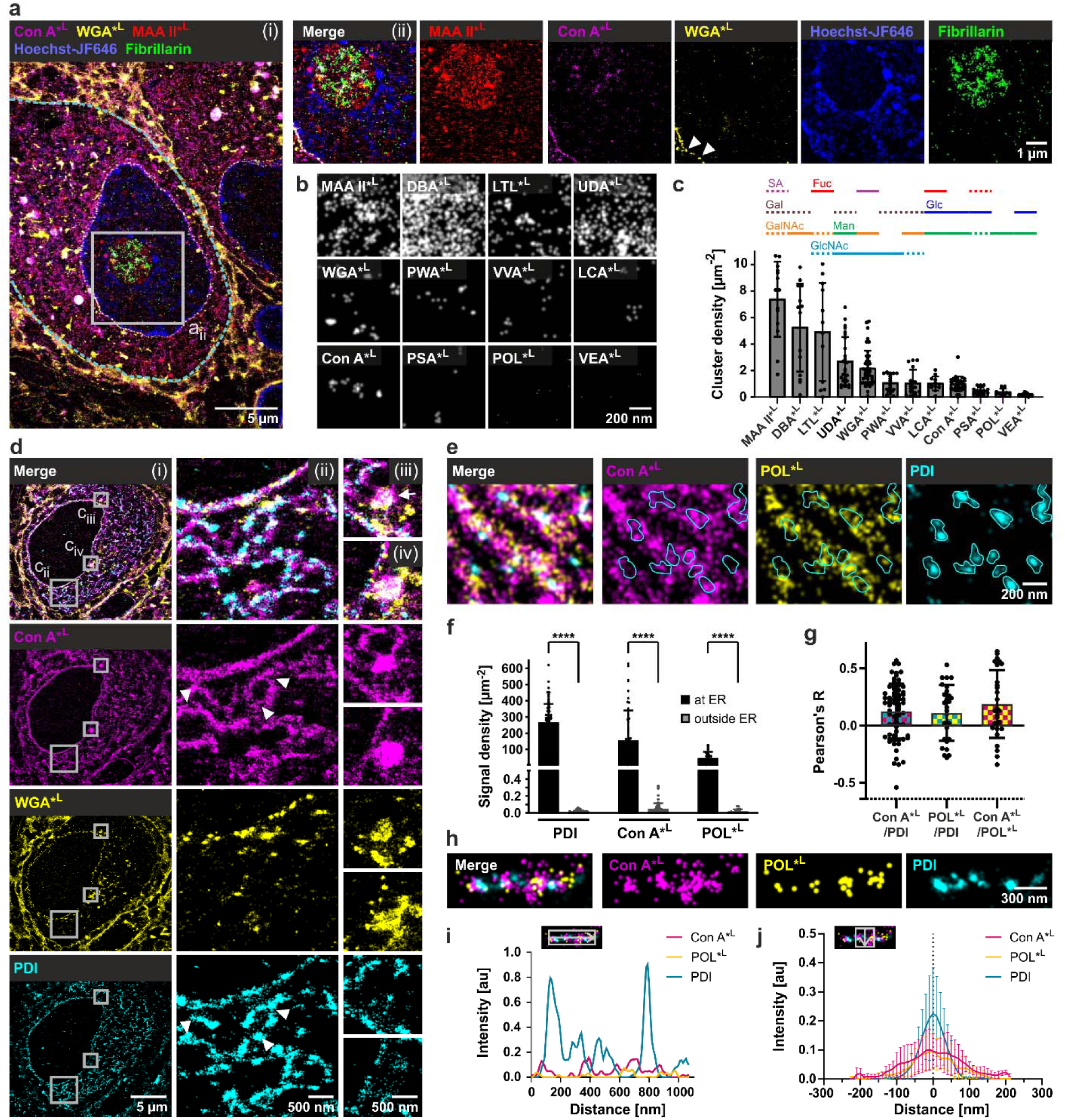
Visualizing nuclear glycosylation patterns and the nanostructure of the endoplasmic reticulum using multiplexed Glyco-STORM. **a**, Super-resolution image of a postsynaptic principal cell soma (delineated by the dashed cyan line) in brain tissue showing an overlay of five markers (i): lectin targets Con A, WGA, and MAA II, a nucleolus antibody marker against Fibrillarin (*d*STORM imaging), and the nucleus marker Hoechst-JF646 (PAINT imaging). Magnified views (ii) depict perinuclear binding of Con A, nuclear pore staining by WGA, and enriched nucleolar as well as moderate nuclear signal of MAA II. **b,** Exemplary images showing signals of 12 lectins within the nucleus. **c,** Density of nuclear clusters. Bar plots represent means ±SD from 3 to 8 independent experiments per lectin (*3* ≤ *N* ≤8) with 3 to 8 measured areas per experiment (3 ≤ *n* ≤ 8). Solid lines highlight predominant sugar specificity; dashed lines denote sugar moieties that are primarily bound as part of a glycan motif. Abbreviations: Fuc, fucose; Gal, galactose; GalNAc, N-acetylgalactosamine; Glc, glucose; GlcNAc, N-acetylglucosamine; Man, mannose; SA, sialic acid. d, *d*STORM images in brain tissue showing that Con A signal colocalizes with the endoplasmic reticulum (ER) antibody marker PDI (i,ii). The perinuclear portion of the Con A signal surrounds the nuclear pore staining of WGA. Con A also displays signal at round structures that do not contain the ER marker PDI but are partially superimposed by WGA signal (iii,iv). e, *d*STORM image showing enriched binding of lectin markers POL and Con A at the ER. **f,** Localization density of antibody and lectin markers within ER segments (total of *n* = 25 to 73) from 6 to 15 principal cells (*6* ≤ *N* ≤15) shown as mean ±SD. Asterisks denote statistical significance (****p ≤ 0.0001; Mann-Whiteney U test). **g,** Colocalization analysis using Pearson’s R test. Bars represent mean ±SD. **h,i,** Close-up view of an ER segment (h) with an exemplified longitudinal signal intensity profile (i) showing an anti-correlation between the PDI signal and the lectin markers. **j,** Averaged intensity distribution from multiple ER segments (*n* = 9) from 3 experiments (*N* = 3) with the line profiles oriented perpendicularly to the segments. Error bars represent SD.

### Suborganellar domains in the endoplasmic reticulum

Next, we aimed to verify the binding of Con A and POL at the ER (Fig. 1c). Indeed, Glyco-STORM imaging of these two mannose-binding lectins with the antibody marker against the ER-specific protein disulfide isomerase (PDI) revealed their common distribution along ER tubules (Fig. 2d,e), while WGA did not show ER-specific staining (Fig. 2d). To evaluate the specificity of Con A and POL for ER structures, we quantified the number of localizations in individually selected ER segments containing PDI signal and adjacent cytoplasmic areas devoid of PDI. Quantification in ER segments revealed the highest signal density for the anti-PDI antibody, followed by the lectins Con A and POL (Fig. 2f). When exploring adjacent PDI-free regions that were considered to lie outside of the ER, Con A and POL showed localization densities comparable to background levels. This indicates that Con A and POL predominantly target the ER, confirming the long-standing recognition of Con A as an ER-specific label^24,25^ and introducing POL as a novel ER marker. However, Glyco-STORM revealed that aside from binding to the reticular ER structures, Con A (Fig. 2d_iii__,iv_) and POL also bound ER-embedded spherical or ellipsoidal structures negative for PDI that were also positive for WGA signal.

Closer examination of the distribution of ER-binding lectins revealed that, although PDI showed higher signal densities compared to Con A and POL (Fig. 2f), it clustered in a confined area, while in most experiments Con A and POL displayed a rather abundant distribution along ER segments (Fig. 2d_ii_,e,h). Interestingly, local regions positive for the PDI signal often showed reduced Con A and POL signals (arrowheads in Fig. 2d_ii_, PDI footprints in 2e). This was also confirmed by very low colocalization between Con A or POL with PDI, despite blurring with a sigma of 2 (Fig. 2g). Furthermore, line profiles along ER tubules often revealed anti-correlation between the lectins and PDI (Fig. 2h,i). This anti-colocalization of PDI and high-mannose-binding lectins could be indicative of PDI’s binding preference for non-glycosylated proteins (for alternative explanations, see Supplementary Discussion, section 4.2). Averaged line profiles laid perpendicularly to the ER tubules unraveled a broader distribution of lectins compared to PDI (Fig. 2j) with a full width at half maximum (FWHM) of 184.5 nm and 143.9 nm for Con A and POL, respectively, and 77.5 nm for PDI. Given that mammalian ER tubules have a diameter of approximately 50 to 100 nm^26,27^, a FWHM considerably above 100 nm suggests that Con A and POL not only bind intra-tubular glycans, but also glycans at the outer ER surface (prior to flippase activity) or the surrounding area.

Thus, ConA and POL in conjunction with PDI labeling revealed distinct sub-compartments of the ER that may infer different functional zones.

### Glyco-STORM imaging elucidates the complex architecture and glycan distribution of the Golgi apparatus

To gain further insights into the architecture of the glycan assembly process, we used lectins to visualize the sequence of carbohydrate attachment at the next station of the ‘glycan assembly line’ – the Golgi apparatus. This complex, convoluted three-dimensional structure consists of axially layered stacks of *cis-*, medial, and *trans-*cisternae, which can be laterally interconnected through non-compact regions to form Golgi ribbons. Glyco-STORM imaging delineated glycan distributions (Fig. 3a, bottom panels) in relation to Golgi antibody markers (Fig. 3a, top panels). The lectins VVA, VEA, MAA II, WGA, and PSA (Fig. 3a,b) stained distinct Golgi and post-Golgi domains, with some, like VVA, matching antibody patterns and others, such as WGA and PSA, revealing a broader, more comprehensive view of the Golgi and downstream structures. First, we analyzed the precise spatial organization of lectins and antibody markers in the *cis*-to *trans*-Golgi direction by averaging line profiles from multiple selections drawn across Golgi cisternae imaged in side view in brain tissue sections (top panel in Fig. 3c). The analysis confirmed the anticipated arrangement of key antibody markers: *cis*-Golgi marker GM130^28^, peaked at 0 nm, which served as a reference point, and TGN38, an integral membrane protein of the Golgi-network (TGN)^29^, displayed a primary peak at 230 nm. For Giantin we observed three peaks of similar height. The first maximum, preceding GM130 at -60 nm, could reflect COPI vesicles known to carry Giantin, which mediates their tethering to the *cis*-Golgi^30^. This peak was followed by a plateau indicating Giantin’s location in the medial Golgi consistent with the conventional notion^31^. The subsequent maxima at 210 nm and 390 nm could reflect Golgi invaginations in non-compact areas. Finally, clathrin marker CHC17 was found downstream of the TGN38 peaks with two maxima: at 540 nm and 950 nm, respectively (Fig. 3b,c). While the first peak corresponds to the budding *trans*-most secretory TGN region, the second peak could indicate clathrin at endosomes and clathrin-coated vesicles further away from the core TGN area.

**Fig. 3.**
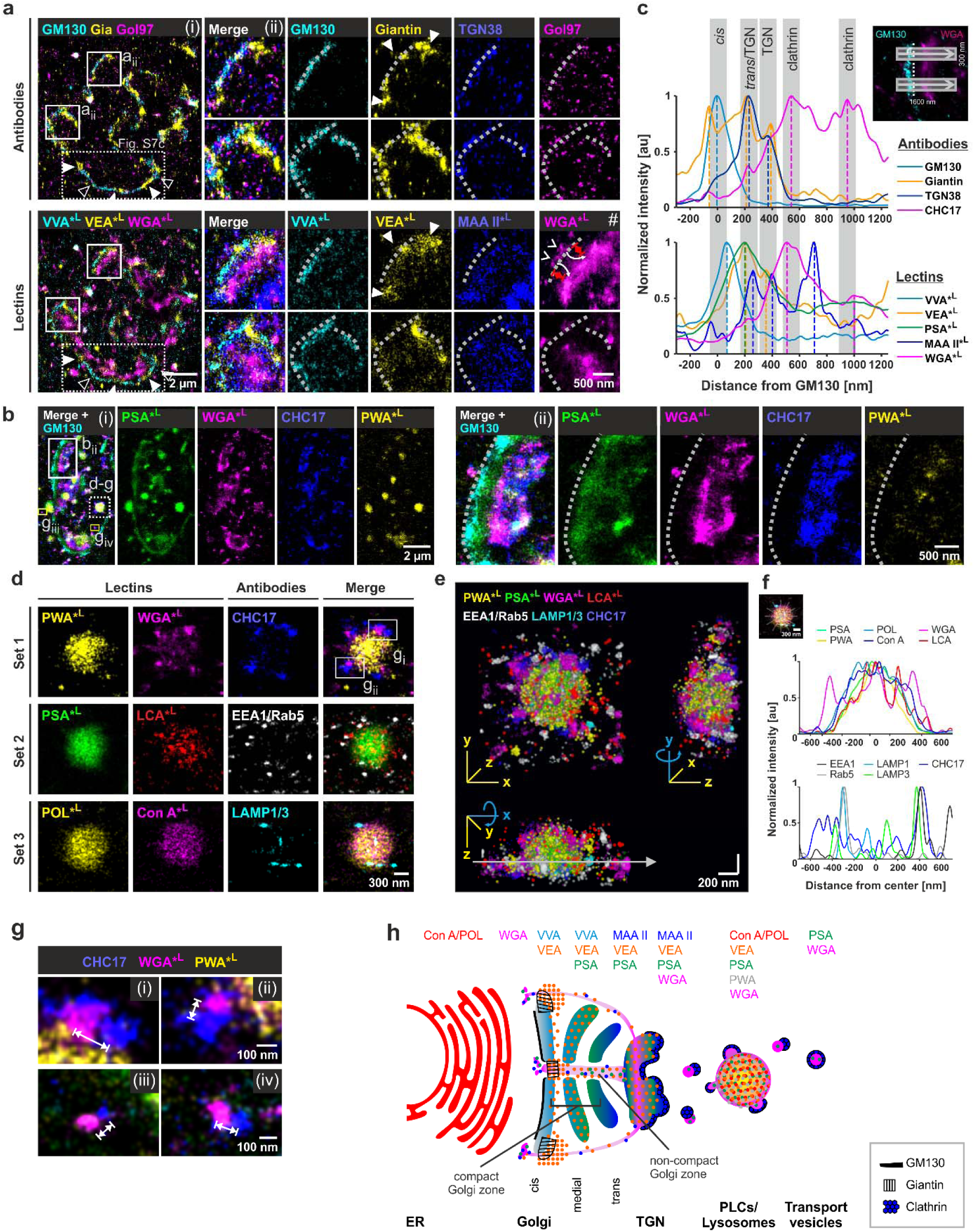
Visualizing local glycosylation at the Golgi apparatus and lysosomes using fluorescently labeled lectins. **a**, Golgi apparatus nanostructure visualized by conventional antibody markers and glycosylation map resolved by Glyco-STORM imaging of Golgi-associated lectins in brain tissue. VVA, VEA, MAA II, and WGA reveal the *in situ* glycosylation status of glycans as they traverse the Golgi apparatus, progressing from the *cis*-Golgi to the TGN and beyond. Giantin signal (filled arrowheads) flanks Golgi stacks with increased GM130 signal (open arrowheads). VEA signal enrichment correlates with Giantin. **b,** Additional example of a Golgi ribbon (i), boxed area magnified in (ii). clathrin-coated vesicles visualized by clathrin marker CHC17. Dashed lines in **a** and **b** delineate the outer rim of the *cis*-Golgi determined based on the antibody marker GM130. **c,** Multiple line profiles drawn across Golgi cisternae were averaged to map the positions of the individual lectins relative to conventional antibody markers. Dashed lines represent signal maxima. The global maximum of the GM130 peak served as a reference point and was set to 0. For better perception, line profiles were normalized, with the global maxima of each marker scaled to 1. VVA, VEA, PSA, Giantin, TGN38: *n* = 55 line profiles from *N* = 4 individual experiments. WGA and GM130: *n* = 79, *N* = 5. CHC17: n = 35, *N* = 3; MAA II: *n* = 21, *N* = 2. **d-g,** Localization of antibody markers and lectins in pre-lysosomal compartments (PLCs) or lysosomes and in their proximity. **d,** PLCs/lysosomes bound by early endosome, lysosome, and clathrin markers at the periphery while the majority of lectins bind within the interior. **e,** 3D rendering of the PLC/lysosome shown in **d**. **f,** Averaged diagram of line profiles rotated by 30° on the exemplary PLC/lysosome demonstrating the localization of the WGA and LCA signal at the periphery of the lysosome along with clathrin, early endosome, and peripheral lysosome markers. **g,** Magnified views of WGA signal paired with juxtaposed clathrin signal at lysosome periphery (i, ii magnified from **d**) and beyond (iii, iv magnified from **b**). **h,** Model of the Golgi-ribbon organization proposing that compact stack regions are laterally organized by Giantin, which delineates non-compact areas allowing a *cis*-to-*trans* passage for vesicles that could by positive for certain lectins, such as WGA.

We delineated a distinct stratification of the ‘glycan assembly line’ by mapping the lectin signals (Fig. 3c, bottom panel). VVA with a specificity for terminal GalNAc and GalNAcβ1-4GlcNAc peaked at 70 nm and presumably visualizes the initial steps of O-linked glycosylation in the *cis*- and medial Golgi cisternae. Next, PSA peaked at 200 nm, attributable to recognizing core-fucosylated N-glycans in the medial and *trans*-Golgi and the TGN. Furthermore, PSA featured a broad secondary peak at 1000 nm, coinciding with the outermost clathrin peak. VEA, whose binding was initially described as α-mannose- and α-glucose-specific^32^, although data on its specificity is limited, displayed a medial Golgi-to-TGN distribution similar to that of PSA, likely reflecting the distribution of high-mannose N-glycans that have not (yet) undergone further terminal modifications. MAA II showed three peaks, with 260 nm corresponding to the proximal TGN regions adjacent to the *trans*-Golgi stack and a 400 nm maximum in the distal TGN regions near the clathrin-coated TGN exit sites. This is in line with MAA II’s specificity for sialylated glycans and sulfated β-galactose that are mainly generated in the TGN. A third peak was located at 710 nm, beyond the first prominent clathrin peak, and could reflect the cytoplasmic signal of MAA II. WGA showed a plateau at 210 nm, indicating its minor, though detectable, presence in the *trans*-Golgi, where it can bind more complex glycans with terminal β-linked GlcNAc or GalNAc. This was followed by the primary peak at 510 nm, corresponding to the distal sub-domain of TGN, attributable to its SA-specificity, and a secondary peak at 1000 nm beyond the region typically recognized as the TGN. Interestingly, although no full colocalization between CHC17 and WGA could be observed (Fig. 3b), their two main post-Golgi peaks showed high correlation. Together, Glyco-STORM imaging provides the first super-resolution visualization of glycan assembly progression within the Golgi through direct staining against multiple carbohydrate motifs, revealing a distinct *cis*–*trans* layering that aligns with previous findings primarily inferred from biochemical and immunofluorescence studies (Fig. 3h).

Next, we investigated the lateral distribution of antibody and lectin markers along the cisternal axis. Earlier electron tomography reported the inter-stack areas as being non-compact regions characterized by fenestrated cisternae filled with endosomes, vesicles, and tubules^33,34^. Recent studies showed that Giantin plays an important role in the lateral organization of Golgi stacks^35–37^. Knockdown of Giantin was previously found to result in reduced fenestration of cisternae in the non-compact region^36^. Recent super-resolution microscopy studies using nocodazole-treated cells demonstrated that Giantin is located at the rim of Golgi mini-stacks imaged *en face*^38,39^. However, the exact position of Giantin in the larger, complex Golgi ribbons of intact cells has been less clearly defined, with only one report, to our knowledge, indicating its inter-stack location^40^. Here, we directly visualized Giantin adjacent to the GM130 signal (filled arrowheads in Fig. 3a, Fig. S7b,c), suggesting its presence at regions flanking or intermitting individual Golgi stacks. In some experiments (Fig. 3a; Fig. S7a_i_) but not all (Fig. S7a_ii-iv_), these regions exhibited elevated levels of α-mannose- and α-glucose-binding lectin VEA (filled arrowheads in Fig. 3a_ii_), suggesting glycosylation heterogeneity along the Golgi’s lateral axis alongside the *cis*-to-*trans* glycosylation progression. In addition, especially in Giantin-rich regions, we observed a pronounced WGA signal preceding the *cis*-most cisterns delineated by GM130 (chevron arrowheads in Fig. 3a_ii_(buttom, #); dashed arrows in Fig. S7b,c). As this signal progresses along the *cis*-to-*trans* axis, it becomes subtler in the region between the *cis* and *trans* faces and intensifies again as it unites with the WGA-rich TGN parts (curved arrows). In contrast, GM130-rich segments devoid of Giantin showed only trace amounts of WGA signal, which predominantly emerged in the rear TGN and post-TGN areas (the distance between the GM130 core and the predominant onset of WGA signal is denoted by red double-headed arrows). Collectively, these observations suggest that lectins, such as VEA and WGA may be enriched in the non-compact regions of the Golgi, which could be enclosed by Giantin (Fig. 3h). These regions are proposed to be transport highways that enable dynamic anterograde and retrograde transport bypassing Golgi stacks^34^.

We extended our investigations to hippocampal and U2OS cells. While imaging of non-sectioned voluminous cells and the anatomical peculiarities of their Golgi limited the detection of side view orientation and precluded quantitative analysis, specific lectin binding to the Golgi was confirmed in these models as well (Fig. S8).

Taken together we demonstrate that lectin-aided imaging can resolve individual layers and can be used to subdivide the Golgi into local domains both in the *cis*-to-*trans* and the lateral direction. Mapping glycan products, rather than the biosynthetic machinery, along the ‘glycan assembly line’ opens a new experimental readout for mechanistic glycosylation studies.

### Lectin tagging of pre-lysosomal compartments, lysosomes, and clathrin-coated transport vesicles

Confocal microscopy screening revealed specific binding of PWA to organelles positive for cathepsin D, a marker for pre-lysosomal compartments (PLCs) and lysosomes (Fig. 1e, Fig. S9a). On the nanoscopic level, PWA staining was abundant (Fig. 3d) and predominantly anti-colocalized with cathepsin D signal that appeared in distinct nano-clusters (Fig. S8b). Several other lectins displayed noticeable signals inside and/or around these organelles, with PSA showing highest signal density followed by high performance of WGA, PWA, Con A, POL, LCA (Fig. 3d; Fig. S9c,d), and VEA (Fig. S9c). Interestingly, among the tested lectins PWA was the only lectin whose binding was almost exclusively restricted to PLCs/lysosomes (apart from the dispersed nucleocytoplasmic staining), possibly pointing to its specificity for certain glycans produced during degradation. To characterize the lectin specificity more precisely, we included further organelle markers: CHC17 for clathrin-coated vesicles, EEA1 and Rab5 for early endosomes, and lysosome markers LAMP1 and LAMP3. We found abundant clusters positive for early endosome markers and very sparse lysosome marker signal around the organelle (Fig. 3d,e). A rotated averaged line profile revealed that while PWA, PSA, POL, and Con A were mainly enriched in the center of the spherical organelle, WGA and LCA showed additional signal at the periphery of PLCs/lysosomes (Fig. 3f). Possible explanations for the peripheral staining by these lectins could be their binding to glycoproteins on the surface of PLCs/lysosomes or on/in transport vesicles. Inspection of an optical 2D section through the lysosome revealed a high overlap between PSA, POL, and PWA at nanoscale while Con A’s and WGA’s distribution appeared less correlated with these lectins (exemplified in Fig. S9d). It is of note that in some PWA-positive organelles, Con A and POL did not show pronounced luminal signal, but instead displayed large domains in the periphery (Fig. S9e). Furthermore, other tested models showed similar enriched lectin signals in large round organelles (Fig. S10a,b). For example, U2OS cells (Fig. S10a) displayed PWA, WGA, PSA, VEA, and POL signals. Interestingly, in these cells, the organelles were clearly surrounded by the early endosome marker EEA1, with Con A coinciding at these sites, suggesting that these regions may participate in processes such as the recycling of high-mannose glycans to the ER.

Many PWA-positive organelles were surrounded by individual clathrin patches that were paired with WGA clusters. WGA and CHC17 showed no direct colocalization (double-headed arrows in Fig. 3g_i__,ii_). These structures could represent clathrin-coated transport vesicles that bud from the PLCs/lysosomes or newly arriving vesicles that deliver lysosomal enzymes from the TGN, with WGA binding to the vesicle contents. Interestingly, in most cases the WGA signal was not fully covered by the clathrin coat. This polarized localization of clathrin relative to the WGA signal extended beyond the immediate vicinity of PLCs/lysosomes and could be found throughout the cytosol (Fig. 3g_iii__,iv_; magnified from boxed regions in Fig. 3b). This could point towards the asymmetric architecture of clathrin-coated transport vesicles with the clathrin coat remaining at one side of TGN-, PLC-, or lysosome-derived vesicles that underwent fission^41,42^ (Fig 3h).

In summary, we show that lectins can be used to visualize nanoscale distribution of glycans inside PLCs and lysosomes and to unravel the relationship between glycans and clathrin vesicles or pits.

### Nanoscale glycan landscape of synapses

To assess the applicability of lectins to study synaptic architecture, we screened lectins for their presence in presynaptic compartments of the calyx of Held, synaptic contacts, and plasma membranes of pre- and postsynaptic cells in brain sections (Fig. 4a,b). Glyco-STORM imaging accompanied by synaptic markers revealed two lectins – PSA and WGA – delineating the synaptic membrane. In particular, WGA and PSA delineated presynaptic compartments occupied by synaptic marker Rab3a (Fig. 4c, blue in 1^st^ column). Notably, both lectins showed enriched signals at synaptic contacts positive for active zone (AZ) markers Bassoon and Piccolo or postsynaptic density (PSD) marker Homer1b/c (Fig. 4c; PSA enrichment is indicated by arrowheads in the 2^nd^ column). This suggests that synaptic contacts harbor increased numbers of glycans presumably rich in core fucosylation recognized by PSA, as well as other types of glycosylation broadly detected by WGA, including terminal N-acetylglucosamine and sialic acid – features predominantly characteristic of mature N-glycans ^43^. Together, the concurrent binding of WGA and PSA further supports the enrichment of mature glycans within the synaptic contact area.

**Fig. 4.**
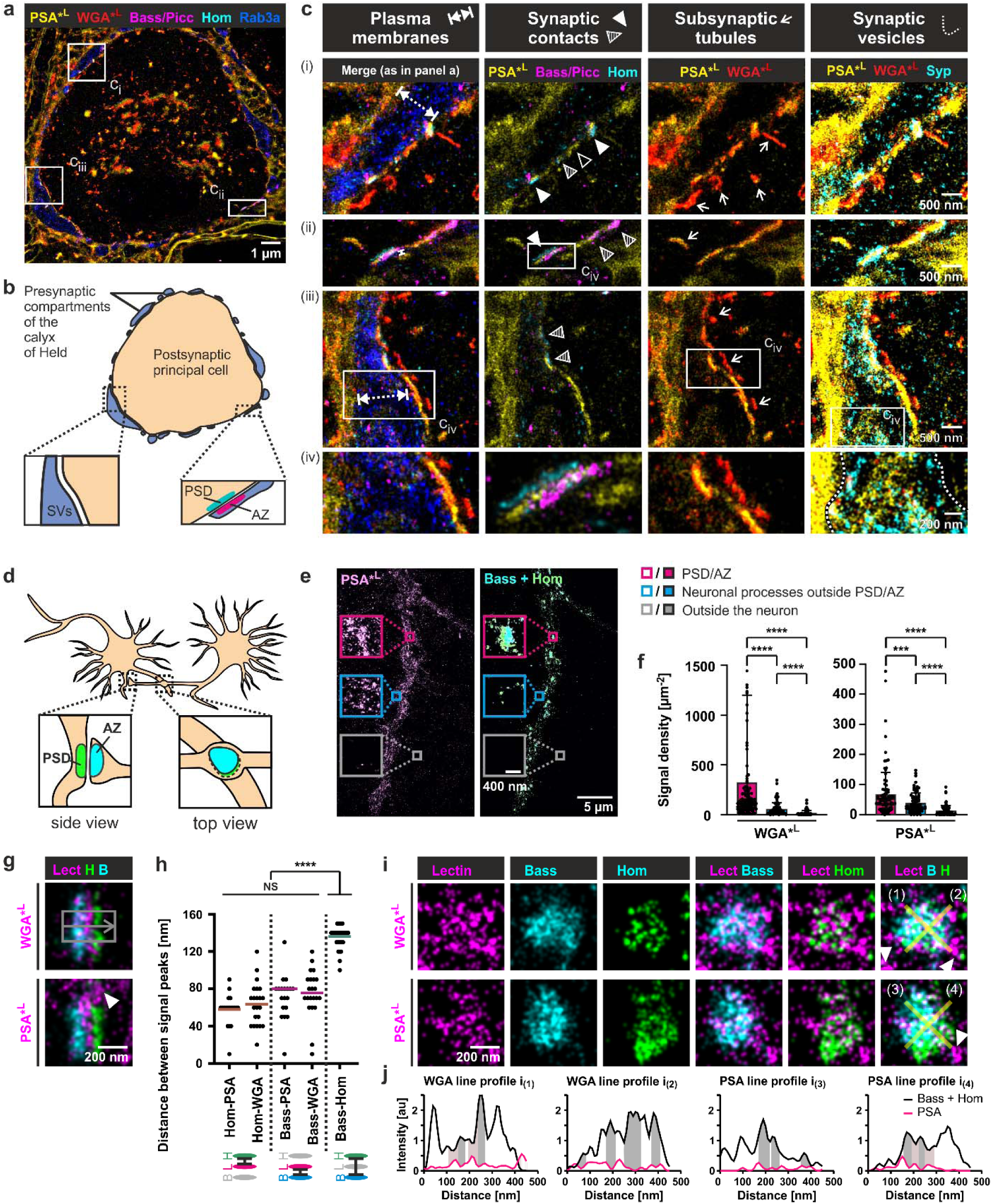
Visualization of synaptic structures by WGA and PSA in neuronal tissue and hippocampal neurons. **a-c**, Neuronal tissue. **a,** Glyco-STORM image with fluorescently labeled lectins PSA and WGA and antibody stainings of Bassoon (Bass), Piccolo (Picc), Homer (Hom), and Rab3a. Boxes denote magnified areas shown in **c**. **b,** Schematic illustration of the calyx of Held synapse corresponding to the image shown in **a**. Abbreviations: SVs, synaptic vesicles; AZ, active zone; PSD, postsynaptic density. **c,** Regions (i-iii) defined in panel **a** and additional magnification (iv) of regions boxed in ii or iii. 1^st^ column i-iv: PSA and WGA delineate plasma membranes (double-headed arrows) surrounding the presynaptic marker Rab3a. 2^nd^ column i-iv: PSA and WGA are enriched at synaptic contacts between PSD marker Homer and AZ markers Bassoon/Piccolo (filled arrowheads), at sites positive for either Bassoon/Piccolo or Homer (striped arrowheads), and at membrane stretches lacking PSD and AZ marker signal (open arrowheads). 3^rd^ column i-iv: PSA and WGA reveal subsynaptic tubules at the postsynaptic side of most synaptic contacts. 4^th^ column i-iv: PSA and WGA are present in areas occupied by SV marker Synaptophysin (Syp; presynapse outlined by dashed lines). **d-i,** Hippocampal neurons. **d,** Illustration of hippocampal neurons forming synaptic contacts that can be imaged either from the side or the top view orientation (*en face*). **e,** Highly clustered PSA signal at synaptic contacts (magenta box), sparser clusters found along neuronal processes negative for synaptic contact markers (blue box), and no measurable signal was present at the glass surface (gray box). **f,** Quantification of Homer-positive areas of *en face* synaptic contacts confirming high PSA and WGA signal density at these sites. WGA, *n* = 114 synaptic contacts from *N* = 8 individual experiments; PSA, *n* = 83, *N* = 5; Data is shown as mean ±SD, asterisks denote statistical significance (****p ≤ 0.0001, ***p ≤ 0.001; Mann-Whiteney U test). **g,** Synaptic contacts imaged from the side with WGA or PSA in the synaptic cleft. Arrow: PSA signal beyond synaptic contacts. **h,** Mean distances between line profile peaks of antibody marker and lectin signals revealing that PSA and WGA are enriched in the center of the synaptic cleft. Hom and Bass: *η* = 110 synaptic interfaces in total with 3 to 10 averaged per image resulting in *n* = 31 data points from *N* = 12 individual experiments; WGA: *η* = 91, *n* = 22, *N* = 8; PSA: *η* = 32, *n* = 19, *N* = 5; one-way ANOVA with Tukey’s multiple comparisons test (****p ≤ 0.0001, NS = not significant). Cartoons illustrating measurements (bottom). **i,** *En face* synaptic contacts with clustered PSA and WGA signals that show partial anti-correlation with Homer or Bassoon staining. **j,** Exemplary line profiles laid through synaptic contacts shown in **i** with Bassoon and Homer signals pooled together. Gray- and pink-shaded areas denote anti-correlation between lectins and Bassoon/Homer.

Interestingly, almost every synaptic contact was accompanied by strand-shaped or round structures at the rim of the postsynaptic principal cell that were positive for WGA and PSA (Fig. 4c, 3^rd^ column, arrows). Although these structures were often found in proximity with CHC17, many of them showed only minor overlap with it (Fig. S11a,b), suggesting that these are not only vesicles related to clathrin-mediated endocytosis but also other specializations, e.g., involved in delivery of glycans to the postsynaptic membrane. Interestingly, some of these structures showed traces of GM130 or TGN38 signal (Fig. S11a-d, filled arrowheads), suggesting that certain sub-synaptic, tubular-shaped glycan structures could represent Golgi outposts or satellites.

Co-imaging with synaptic vesicle markers, i.e., Synaptophysin, revealed high abundance of PSA and, to a lesser degree, WGA within synaptic vesicle clouds (Fig. 4c, 4^th^ column). Clarifying the precise relationship of WGA and PSA with synaptic vesicles will facilitate investigations into whether different functional pools of SVs carry distinct glycosylation signatures.

To further investigate the synaptic distribution of WGA and PSA and demonstrate their universal applicability in diverse neuronal models, we performed multiplexed *d*STORM experiments in hippocampal neurons (Fig. 4d-j) and evaluated glycan distribution at synaptic contacts. Both, WGA and PSA were significantly enriched (5.7× for WGA, 1.8× for PSA) at regions occupied by Homer1b/c, as compared to extra-synaptic contact regions within neuronal processes (Fig. 4e,f). Synaptic contacts imaged in profile reveal that WGA and PSA were both localized at the synaptic cleft (Fig. 4g), with WGA and PSA exhibiting some signals beyond the contact sites, protruding into pre- or postsynaptic regions (arrowheads in Fig. 4g,i). Line profiling and Gaussian fitting revealed a distance of 136.1 ± 12.56 nm between signal peaks of Homer1b/c and Bassoon (Fig. 4h), aligning with previous SMLM studies; e.g., by Pauli *et al.* reporting 149 ± 12 nm in hippocampal mossy fibers^44,45^. The N-terminal Bassoon antibody used in our study previously demonstrated a signal peak in ∼70 nm from the plasma membrane^46^; and Homer1 was proposed to peak at ∼52 nm^47^. Given the 20 nm width of the synaptic cleft^48^, the center of the synaptic cleft is expected at a distance of approximately 80 nm from the Bassoon peak or at a distance of 62 nm from Homer1b/c. Both lectins displayed a location at the center of the cleft, with WGA detected at 63.64 ± 23.21 nm and PSA at 57.89 ± 16.86 nm from the Homer peak (Fig. 4h, Fig. S12a,b), indicating that pre- or post-synaptic sides as well as the extracellular matrix could contribute to the signal.

Overlaying WGA or PSA signals with antibodies against Homer1b/c and Bassoon (Fig. 4i) of synaptic contacts imaged *en face* highlights the broad distribution of both lectins in the area determined by AZ/PSD markers. Notably, WGA and PSA nanoclusters often exhibited minimal overlap with Bassoon and Homer1b/c, instead occupying Bassoon- and Homer1b/c-devoid areas, as evident from line profile curves (Fig. 4j, anti-colocalization in areas with peaking antibody or lectin signal is exemplarily denoted by gray- or pink-shaded areas). The anti-colocalization was more pronounced in the case of Homer1b/c (5^th^ panel column in Fig. 4i). Given that Homer1b/c is a scaffold protein that is involved in neurotransmitter receptor organization, WGA and PSA could reflect the positions of glycosylated receptors intermitted (or indirectly interconnected) by Homer1b/c (for details and alternative interpretation, see section 4.5 of Supplementary Discussion).

In conclusion, we observed that PSA and WGA signals were enriched precisely at the midpoint between synaptic membranes within the synaptic clefts, in SV-occupied regions, and at specializations in close proximity to PSDs. The lectins revealed a novel tubular structure at the postsynaptic side, in the proximity of synaptic contacts. Furthermore, the alternating signals of Homer1b/c and WGA demonstrate that glycosylation could serve as a blue-print to decipher synaptic organization. The concomitant binding of WGA and PSA is associated with mature glycans, indicating that specific glycosylation patterns contribute to synaptic maturity and organization.

### Probing biological questions with Glyco-STORM

Building on the close examination of individual glycosylation patterns across various organelles and structures, we aimed to explore the broader capabilities of Glyco-STORM. High-resolution, multiplexed glycosylation mapping via Glyco-STORM enabled detailed glycan co-profiling, advancing the field of spatial glycomics, as illustrated for brain tissue in Fig. 5a (for more examples, see Fig. S13).

**Fig. 5.**
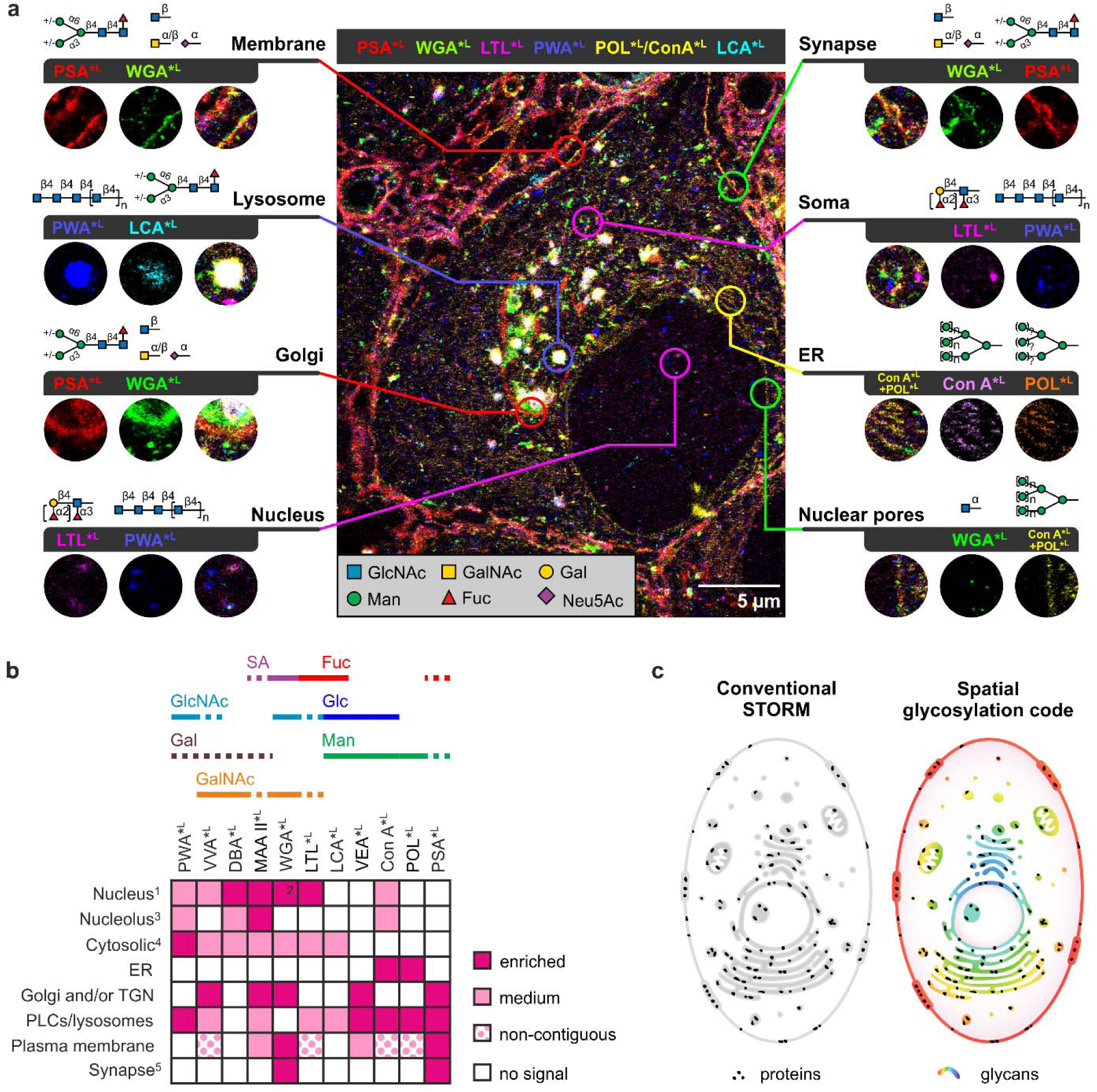
Profiling the glycan landscape using multiplexed lectin-based *d*STORM. **a**, Glycosylation map from a multiplexed Glyco-STORM experiment in brain tissue highlighting binding of lectins with different carbohydrate specificities to distinct cellular structures. **b,** Performance matrix of 11 lectins evaluated in the Glyco-STORM approach, showing their binding across diverse cellular structures and organelles. Solid lines highlight predominant sugar specificity; dashed lines denote sugar moieties that are primarily bound as part of a glycan motif. For a complete overview of all screened lectins, including those tested only by confocal screening, see Fig. S14. ^1^Nucleus staining refers to clustered or dispersed signal in the nucleoplasm if not stated otherwise. ^2^In addition to the clustered signal in the nucleoplasm, WGA stains the nuclear pores. ^3^Nucleolar signal was denoted as present if detected in a subset of cells. ^4^Cytosolic localization refers to clustered or dispersed signal patterns. ^5^Synaptic localization refers only to brain tissue model. Abbreviations: ER, endoplasmic reticulum; Fuc, fucose; Gal, galactose; GalNAc, N-acetylgalactosamine; Glc, glucose; GlcNAc, N-acetylglucosamine; Man, mannose; PLCs, pre-lysosomal compartments; SA, sialic acid; TGN, *trans*-Golgi network. **c,** Glyco-STORM allows to super-resolve organelles and specializations with common glycosylation signatures throughout the cell, revealing new organization patterns and obscured relationships.

To provide a comprehensive overview of lectin specificities, we validated lectin signals by specific antibody markers in at least three independent staining experiments for each lectin in neuronal tissue. In addition, for some lectins, we verified lectin signal consistency in U2OS cells and hippocampal neurons (e.g., see Fig. S8, Fig. S10, Fig. S13, Tables S9 and S10). The binding profiles of the best-studied lectins tested with Glyco-STORM are summarized in a performance matrix in Fig. 5b, while a comprehensive matrix of all screened lectins, including those tested only by confocal screening, is provided in Fig. S14.

This study provided new insights on glycosylation in different organelles and specializations of neuronal and non-neuronal cells, triggering new questions. In Table 1 we have compiled a guide for future lectin-aided studies on the roles of glycans in cellular and neurological functions and summarized scientific questions and hypotheses that Glyco-STORM can address.

**Table 1.**
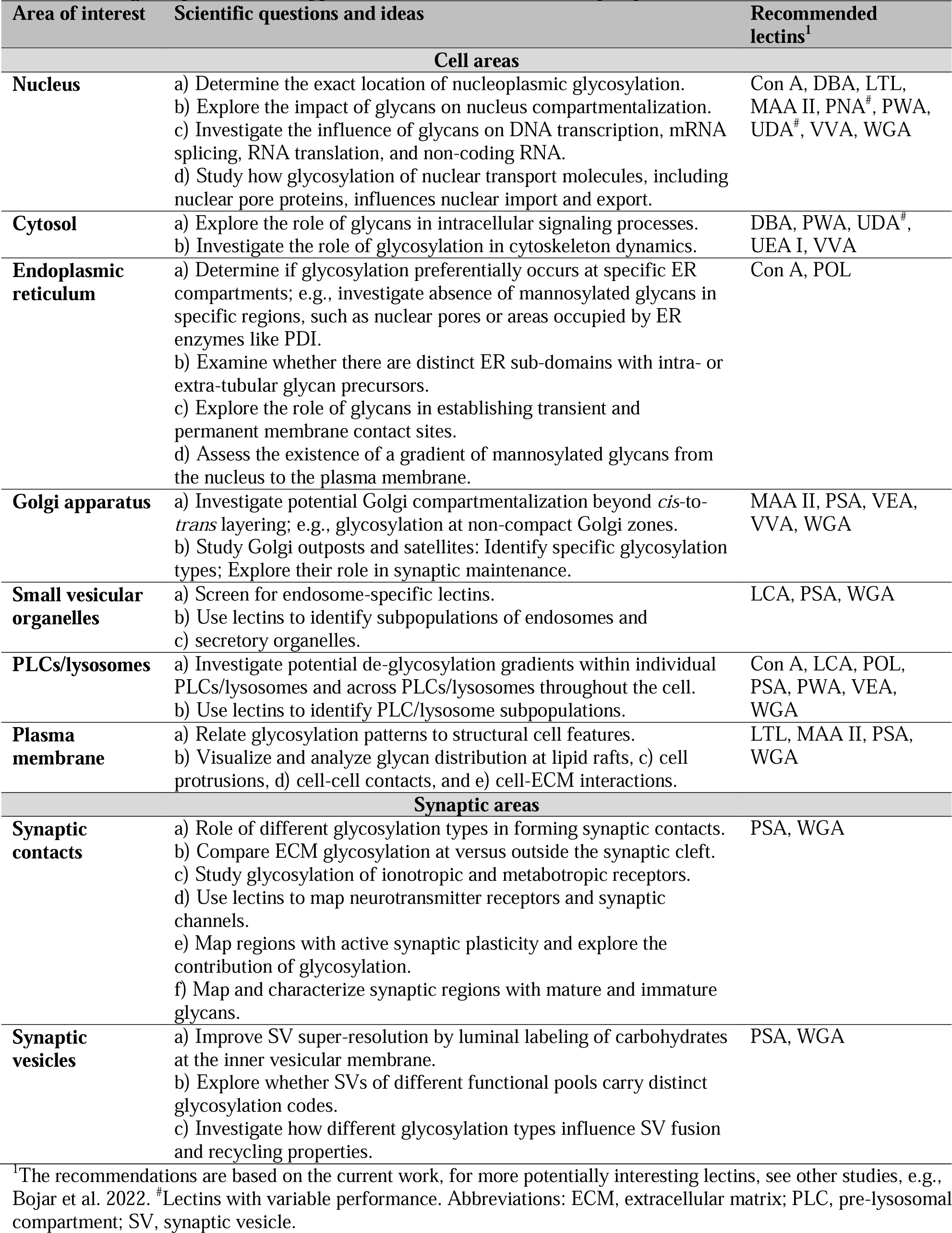
Biological questions and hypotheses that can be addressed by Glyco-STORM.

| Area of interest | Scientific questions and ideas | Recommended lectins<sup>1</sup> |
| --- | --- | --- |
| **Cell areas** |  |  |
| **Nucleus** | a) Determine the exact location of nucleoplasmic glycosylation.<br>b) Explore the impact of glycans on nucleus compartmentalization.<br>c) Investigate the influence of glycans on DNA transcription, mRNA splicing, RNA translation, and non-coding RNA.<br>d) Study how glycosylation of nuclear transport molecules, including nuclear pore proteins, influences nuclear import and export. | Con A, DBA, LTL, MAA II, PNA<sup>#</sup>, PWA, UDA<sup>#</sup>, VVA, WGA |
| **Cytosol** | a) Explore the role of glycans in intracellular signaling processes.<br>b) Investigate the role of glycosylation in cytoskeleton dynamics. | DBA, PWA, UDA<sup>#</sup>, UEA I, VVA |
| **Endoplasmic reticulum** | a) Determine if glycosylation preferentially occurs at specific ER compartments; e.g., investigate absence of mannosylated glycans in specific regions, such as nuclear pores or areas occupied by ER enzymes like PDI.<br>b) Examine whether there are distinct ER sub-domains with intra- or extra-tubular glycan precursors.<br>c) Explore the role of glycans in establishing transient and permanent membrane contact sites.<br>d) Assess the existence of a gradient of mannosylated glycans from the nucleus to the plasma membrane. | Con A, POL |
| **Golgi apparatus** | a) Investigate potential Golgi compartmentalization beyond *cis*-to-*trans* layering; e.g., glycosylation at non-compact Golgi zones.<br>b) Study Golgi outposts and satellites: Identify specific glycosylation types; Explore their role in synaptic maintenance. | MAA II, PSA, VEA, VVA, WGA |
| **Small vesicular organelles** | a) Screen for endosome-specific lectins.<br>b) Use lectins to identify subpopulations of endosomes and<br>c) secretory organelles. | LCA, PSA, WGA |
| **PLCs/lysosomes** | a) Investigate potential de-glycosylation gradients within individual PLCs/lysosomes and across PLCs/lysosomes throughout the cell.<br>b) Use lectins to identify PLC/lysosome subpopulations. | Con A, LCA, POL, PSA, PWA, VEA, WGA |
| **Plasma membrane** | a) Relate glycosylation patterns to structural cell features.<br>b) Visualize and analyze glycan distribution at lipid rafts, c) cell protrusions, d) cell-cell contacts, and e) cell-ECM interactions. | LTL, MAA II, PSA, WGA |
| **Synaptic areas** |  |  |
| **Synaptic contacts** | a) Role of different glycosylation types in forming synaptic contacts.<br>b) Compare ECM glycosylation at versus outside the synaptic cleft.<br>c) Study glycosylation of ionotropic and metabotropic receptors.<br>d) Use lectins to map neurotransmitter receptors and synaptic channels.<br>e) Map regions with active synaptic plasticity and explore the contribution of glycosylation.<br>f) Map and characterize synaptic regions with mature and immature glycans. | PSA, WGA |
| **Synaptic vesicles** | a) Improve SV super-resolution by luminal labeling of carbohydrates at the inner vesicular membrane.<br>b) Explore whether SVs of different functional pools carry distinct glycosylation codes.<br>c) Investigate how different glycosylation types influence SV fusion and recycling properties. | PSA, WGA |

<sup>1</sup>The recommendations are based on the current work, for more potentially interesting lectins, see other studies, e.g., Bojar et al. 2022. <sup>#</sup>Lectins with variable performance. Abbreviations: ECM, extracellular matrix; PLC, pre-lysosomal compartment; SV, synaptic vesicle.

## Discussion

By combining fluorescently labeled lectins with multiplexed super-resolution microscopy, Glyco-STORM reveals the spatial distribution of glycosylation within subcellular and suborganellar compartments at the nanoscale. This approach provides a unique perspective on the cellular landscape by visualizing glycosylation as a distinct level of organization, revealing a ‘spatial glycosylation code’ that complements protein-focused models on cellular organization (Fig. 5c). Critical findings from our study include the delineation of suborganellar glycan domains within the ER, Golgi apparatus, and PLCs/lysosomes, which were previously challenging to resolve. The following examples illustrate glycan domains found using this approach: POL and Con A bound mannose residues in and around ER tubules partially sparing nanodomains occupied by the ER enzyme PDI; at the Golgi, lectins not only allowed to resolve the transversal organization of Golgi stacks but also unveiled lateral domains; and WGA staining in combination with clathrin antibody staining revealed polarized transport vesicles in the vicinity of PLCs/lysosomes. These findings have enriched our understanding of the compartmentalization of glycosylation processes and their contributions to protein maturation, trafficking, and function. Furthermore, the identification of distinct glycosylation patterns associated with synaptic architecture offers novel perspectives on the modulation of synaptic transmission and plasticity. In particular, we demonstrate that PSA and WGA can be used to study clathrin-coated endosomes and distal glycosylation stations, such as Golgi outposts or satellites, and their role for the maintenance of the synaptic contacts; glycosylation of synaptic vesicles and its role for their release probability; and the role of glycosylation for synaptic contact scaffolding, trans-synaptic alignment, and neurotransmission. To our knowledge, so far this is the first super-resolution study on synaptic glycosylation. Looking ahead, the integration of live cell capabilities with lectin-aided super-resolution imaging presents exciting possibilities for future studies. This advancement could ideally be applied to explore the dispatch of Golgi satellites and outposts, to investigate the glycan-dependent organization of glutamate receptors, and to assess the impact of glycosylation on synaptic vesicle (SV) release.

Despite the advances of Glyco-STORM, challenges remain in fully exploiting its potential. Some lectins showed heterogeneous staining patterns (Fig. S2). This can be explained by (1) their binding to O-glycans that are predominantly present at soluble proteins with a potentially higher likelihood of being washed out, (2) the instability of their target glycans, or (3) the reduced affinity between these lectins and their targets. Carbohydrate-stabilizing fixatives, such as periodate-lysine-paraformaldehyde could help to enhance the longevity of these molecules^49^, addressing the first two issues. For lectins with reduced affinity, employing alternative strategies in super-resolution microscopy, such as PAINT, could exploit their weaker binding properties^50^. For example, WGA conjugated to a photoactivatable fluorophore has been already successfully used in PAINT imaging^17^. However, the PAINT approach requires a very laborious optimization of buffer conditions for each lectin, coupled with its inherent demand for extended image acquisition times, limiting its routine applicability.

In the past years super-resolution microscopy has been substantially advanced by groundbreaking techniques like MINFLUX^51^. Fluorophores used for *d*STORM, i.e., AF647 and CF680, are well suited for MINFLUX nanoscopy. Employing directly labeled lectins for MINFLUX opens up new resolution scales below 10 nm. Multiplexed imaging of lectin and immunohistochemically stained samples at this resolution will allow to precisely map glycosylation motifs to individual proteins and other glycoconjugates. While individual lectins typically detect multiple sugar moieties or carbohydrate motifs, creating uncertainty about the precise carbohydrate sequence attached to a molecule, multiplexing different lectins in combination with exceptional resolution will allow for the specification or even full reconstruction of complex carbohydrate chains in individual glycans. With the accumulation of sufficient data for deep learning techniques, spatial glycomics and interaction analyses will facilitate the comparison of glycosylation profiles of diseased versus healthy or treated versus untreated samples.

Altered glycosylation patterns play critical roles in diseases such as cancer and neurodegeneration. In cancer, dysregulated glycosylation could be a key driver of tumor invasion and progression^10^. While single-lectin application with acquisition at limited resolution provides an incomplete view of the glycosylation landscape, super-resolution and multiplexing together amplify our capacity to dissect the complex glycosylation signatures associated with cancer and to map them to small organelles and nanodomains. This paves the way for deeper insights into the mechanisms of cancer development and for the identification of novel biomarkers and possible therapeutic targets. In neurodegenerative diseases, such as Alzheimer’s, Parkinson’s, and multiple sclerosis, altered glycosylation is linked to disease progression^52^. Traditional approaches, focusing on the pathways of proteins without considering their glycosylation status, have proven insufficient for fully comprehending the intricacies of these diseases, leading to unsuccessful treatment outcomes. The minute nature of synaptic specializations, such as SVs, AZs, PSDs, and other nanodomains, necessitates the use of super-resolution techniques. By enabling precise visualization of where and how altered glycosylation occurs at these critical neural junctures, Glyco-STORM can uncover the underlying mechanisms leading to synaptic failure, offering new avenues for effective treatment strategies.

In conclusion, Glyco-STORM has established a new paradigm in glycoscience, offering unprecedented insights into the spatial organization of glycans within cells, enabling insights into their functional implications. As we expand the toolkit of lectins and refine our understanding of glycan functions, we anticipate that Glyco-STORM will continue to illuminate the roles of glycosylation in health and disease, paving the way for novel diagnostic and therapeutic strategies.

## Methods

### Animals

For primary hippocampal cell cultures, pregnant Sprague-Dawley rats at the 19^th^ day of gestation were terminally anesthetized by intraperitoneal injection of sodium-pentobarbital (500 mg/kg Narcoren; Boeringer Ingelheim Vetmedica, Germany), and their embryos were euthanized. Both procedures were carried out in compliance with the German animal welfare guidelines according to protocol T-33/21 approved by animal welfare authorities of Heidelberg University.

For tissue preparation, 12-/13-day-old rats were anesthetized by intraperitoneal injection of 100 mg/kg Ketamine (Bremer Pharma, Germany) and 5 mg/kg Xylazine (Ecuphar, Germany) and transcardially perfused with 20 ml phosphate buffered saline (PBS; Sigma-Aldrich, MO, USA: P3813), followed by 30 ml 4% paraformaldehyde (PFA) in PBS. Perfusion experiments were conducted according to the German animal welfare guidelines – protocol 35-9185.81/G-214/20 – approved by the Regierungspraesidium Karlsruhe.

### U2OS cells

U2OS human osteosarcoma cells were cultured and stored in a mixture of Dulbecco’s Modified Eagle Medium (DMEM) and Ham’s F-12 (DMEM/F-12; Gibco, CA, USA: 11320033) supplemented with 10 % FBS, 100 U/mL penicillin and streptomycin, and 2 % GlutaMAX (Gibco: 35050061) at 37.5 °C in 5% CO_2_. Cells were cultured in 10 mm dishes and split two times per week. For splitting and re-seeding, cell dishes were washed with 5 ml PBS, incubated with 1 ml of 0.5 % Trypsin for 1.5 min at 37.5 °C, and filled with 9 ml medium to stop trypsinization. For imaging experiments, cells were seeded onto fibronectin-coated glass-bottom dishes (MatTek, MA, USA: P35G-0-14-C). After 24 hours in the incubator, cells were twice washed with cytoskeleton-stabilizing buffer (160 mM piperazinediethanesulfonic acid pH 6.8, 10 mM EGTA, 4 mM MgCl_2_) and chemically fixed with 4% PFA/4% sucrose in cytoskeletal buffer for 10 min at 37 °C. Subsequently, cells were washed three times with Tris buffer consisting of 50 mM Tris-HCl and 150 mM NaCl (Sigma-Aldrich: 94158) and stored at 4°C until further usage.

### Hippocampal cells

Hippocampal tissues were excised from E19 embryo brains in ice-cold Dulbecco’s Phosphate Buffered Saline (DPBS; Sigma-Aldrich: D8537) and collected in Hank’s Balanced Salt Solution (HBSS; Invitrogen, CA, USA: 14170088). Hippocampi were then treated with 0.25% trypsin in HBSS for 20 min at 37°C to induce cell dissociation. The enzymatic activity was halted by introducing warm high-glucose DMEM (Invitrogen: 41966029) enriched with 10% FBS, antibiotics (100 U/mL penicillin and streptomycin), and 2 mM L-glutamine at a 2:1 volume ratio for 5 min. Sedimented hippocampal cells were washed and resuspended in 2 ml DMEM, followed by filtering through a cell strainer (Greiner Bio-One, Frickenhausen, Germany: 542000). The cells were then centrifuged at 500 g for 5 min, resuspended in DMEM with added supplements (10% FBS, 100 U/mL penicillin/streptomycin, and 2 mM L-glutamine), and plated onto poly-L-lysine-coated glass-bottom dishes (MatTek, MA, USA: P35G-0-14-C). After 5 h in a 37°C incubator with 5% CO_2_, DMEM was replaced by Neurobasal medium (Invitrogen: 21103-049) supplemented with 1x B-27 (Gibco: 17504044), 2 mM L-glutamine, and 100 U/mL penicillin and streptomycin. Hippocampal neurons were grown for 21 days in the incubator and fixed for experiments as described for U2OS cells.

### Semi-thin brain tissue cryosections

Cryosections were cut, based on the Tokuyasu method^53^ and adapted for *d*STORM imaging^18^. In short, 200 µm thick vibratome sections from the rat brain stem were sliced and the medial nucleus of the trapezoid body (MNTB) region with the calyx of Held synapses was excised under a binocular using a scalpel. The brain-tissue-blocks were infiltrated in 2.1 M sucrose in 0.1 M cacodylic acid buffer overnight at 4°C. Brain tissue samples were mounted on specimen holder pins and plunge-frozen in liquid nitrogen. Subsequently, mounted samples were transferred to the cryo-ultramicrotome chamber (Leica, Wetzlar, Germany: Ultracut S with cryo-chamber EM FCS) and sectioned into 400 nm slices. The slices were transferred to poly-L-lysine-coated glass-bottom dishes. Transfer was realized using a metal loop harboring a droplet of 2.3 M sucrose in 0.1 M cacodylic acid buffer mixed with 2% methylcellulose in a 1:1 ratio. Sucrose-mounted samples were stored at 4 °C until further usage. Prior to staining, tissue samples underwent three consecutive 10-minute washes with Tris buffer to ensure thorough removal of the viscous sucrose/methylcellulose mounting medium.

If multiplex experiments were conducted, nanodiamonds were applied to the coated glass-bottom dishes before cell seeding or tissue application. For a detailed protocol on fiducial conjugation, see Methods section ‘Multiplexed *d*STORM imaging’.

### Lectin and antibody staining

Stainings for *d*STORM imaging and confocal microscopy followed a standardized protocol. U2OS or hippocampal cells were permeabilized by an incubation in a 0.1 % triton solution for 10 min at room temperature (RT). Subsequent procedures were conducted in a similar fashion for all cell and tissue samples. To prevent unspecific antibody staining, 5 % fetal calf serum (FCS) with 5 % glycine in Tris buffer was applied for 10 min onto tissue samples or 30 min onto cell samples at RT. The samples were briefly washed with Tris buffer. For confocal microscopy, samples were incubated for 10 min with a DAPI solution in Tris buffer and washed two times. In the first staining step, primary antibodies diluted in 0.5 % FCS in Tris buffer were applied for 45 min at RT, followed by three washes of 5 min each. In the second staining step, samples were incubated for 45 min with secondary antibodies, directly labeled primary antibodies or other direct labels, and lectins, all diluted in Tris buffer. If only lectins or other direct labels were used, the staining procedure was performed in a single 45 min incubation step. Subsequently, the sample was washed three times and used for imaging. The initial staining was performed manually while all subsequent staining rounds were realized via the automated pipetting robot (see ‘Multiplexed *d*STORM imaging)’.

### Lectin screening

Initial screening in tissue was performed by capturing wide-field images on the maS³TORM setup with the 661 nm laser at approx. 1 W/cm² and an exposure time of 200 ms. For further validation of the 17 selected lectins, the Leica TCS SP8 confocal microscope with a 63× oil objective (NA 1.4) was used. This microscope is equipped with four lasers with wavelengths of

405 nm, 488 nm, 568 nm, and 638 nm and enables simultaneous imaging of different fluorophores. All images were captured with a speed of 100 voxels/second, resulting in a pixel dwell time of 4.88 µs and a frame rate of 0.015 per second. During the line by line scanning, the ‘line average’ was set to 4 for noise reduction. If no substantial staining variation among multiple samples was observed in the initial screening, lectin staining was further characterized by co-stainings with established antibodies that were selected to confirm initial assumptions for lectin localization. The following antibodies were used for orientation: PDI for the ER, GM130 for the Golgi apparatus, cathepsin D and LAMP1,2,3 for lysosomes, and Synaptophysin for synaptic vesicles (Table S3). DAPI was used as a nucleus marker. To prevent any bleed-through of signals between channels, multi-color images were acquired in sequential mode. All lectins with reliable staining performance were screened across a minimum of three different biological replicates, specifically cell samples from different passages or tissue samples from different animals, with images acquired from at least five cells per replicate. Additionally, at least one z-stack per cell sample was captured to appreciate the 3D distribution of the lectin signal.

### Elution experiments with competing carbohydrates

Elution experiments were performed fully automated using our maS³TORM setup equipped with a pipetting robot (Fig. S3b). Images were acquired in the wide-field mode with 100 ms exposure time at five preselected regions of interest per staining. Reference images (*E_R_*) were acquired before elution. Next, the elution buffer specific for individual lectins (Table S5) was applied for 45 min, followed by three 5 min washing steps with Tris buffer. Finally, images were acquired after the elution treatment (*E_T_*) and the post-/pre-elution ratio *E_R_*/*E_T_* was calculated. To correct for elution-unrelated effects such as bleaching from imaging and mechanical stress from washing, we performed a control experiment where the sample was incubated with Tris buffer instead of the sugar-containing elution buffer. In analogy to the elution condition, reference (*C_R_*) and post-treatment images (*C_T_*) were also acquired for the control condition; and the *C_R_/C_T_* ratio was used for normalization. The relative signal remaining after elution was expressed as:

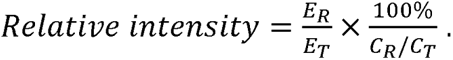

### Multiplexed *d*STORM experiments

#### Dish coating

For a precise registration, fiducials were immobilized by conjugation via an amide coupling reaction to the amino acid residues in the glass-bottom dish coating. To this end, glass-bottom dishes were coated with 0.01% poly-L-lysine solution (PLL, Sigma-Aldrich: P4832) for 30 min and then washed thoroughly with sterile water. Subsequently, nanodiamonds (Adamas Nanotechnologies, NC, USA: NDNV100nmHi2ml) 1:8000-10,000 suspended in water were added for 10 min. The solution was replaced by 6.25 mM EDC for 30 min. The dishes were washed 5-7 times with water before letting them dry completely for 60 min. For U2OS cells, instead of PLL, 1 mg/ml fibronectin (Sigma-Aldrich: 11051407001) was applied for 60 min. The used dilution resulted in 5-10 fiducials per field of view. For cell experiments, conjugation was done under sterile bench conditions.

#### Experiment preparation

After the initial staining round, the sample-containing glass-bottom dish was placed onto the motorized stage. For automated-multiplexed experiments, the staining and imaging steps were programmed via the in-house developed Experiment Editor software^18^ that directed the coordinated performance of the custom-built *d*STORM setup and the PAL3 RTC (Axel Semrau, Sprockhövel, Germany) pipetting robot. Vials required for the long-term experiment were placed in the tray for the robot and prefilled with: imaging buffer components, elution buffer, blocking solution, re-staining solutions with diluted primary and secondary antibodies, lectins, and other labels. For experiments that lasted for more than one day, re-staining solutions and blocking buffers were freshly added each day. Buffers were supplied via the washing station. Five regions of interest (ROIs) and one reference position for the autofocus with 5 to 10 fiducials in the sample-free glass area were selected per experiment. The lateral coordinates and individual focus positions for the fiducials were transferred to the respective modules of the Experiment Editor. To prevent bleaching between close regions, ROIs were selected at a minimal distance of 100 nm. The experiment was executed by the Experiment Editor software in a fully automated manner.

#### Imaging buffer preparation and microscopy

The imaging buffer was freshly prepared using the pipetting robot. First, Mercaptoethylamine (MEA) powder was dissolved in Tris buffer with 50 mM Tris-HCl and 150 mM NaCl. Then, KOH was added to adjust the final concentrations to 85 mM MEA and 35 mM KOH, achieving a pH of 8 to 8.1.

After the imaging buffer was applied to the sample, the imaging series begun with the recording of the fiducial reference position for the automated refocusing. Then a standardized recording sequence was conducted for each of the five selected ROIs. First, a wide-field image was acquired with the 488 nm laser, followed by a pause time of 3 minutes. The autofluorescence image of the sample captured in the green channel helped to verify the correct position of the sample in the field of view, which allowed intervention in case of gross deviations. Second, a wide-field image was acquired with the 661 nm laser. In the third step, the *d*STORM acquisition was conducted using the 661 nm laser at a high power setting of ∼2 kW/cm^2^, with a 30 ms exposure time for 20,000 frames. Fourth, for image registration across imaging rounds, for each ROI a fiducial image series was acquired using the 561 nm laser at 0.08-0.12 kW/cm^2^, with an exposure time of 200 ms for 50 frames. Fiducial emission was collected via the far-red filter that is also used for *d*STORM imaging, allowing fiducial-based transformations without the need to correct for chromatic aberrations. This approach was repeated until all five ROIs were imaged.

#### Signal removal and re-staining

To remove residual fluorescence signal from previous rounds, the imaging buffer was exchanged by Tris buffer and all ROIs were subjected to high power illumination by the 661 nm laser at ∼2 kW/cm^2^ combined with the 405 nm laser at ∼0.7 kW/cm² for 4 min. If secondary antibodies directed against a host species that was already employed in the same multiplex experiment were to be used for the next re-staining, samples were subjected to chemical elution with 0.1 % SDS elution buffer at pH 13 for 15 min. The effectiveness of photobleaching for fluorophore destruction and partial photo-unbinding of fluorophore-conjugated labels, alongside with the SDS elution for antibody stripping was demonstrated in our previous work^18^.

Re-staining with a new set of labels was realized by the pipetting robot according to the protocol previously outlined for manual staining procedures. Subsequently, the entire cycle of refocusing, re-imaging, signal removal, and re-staining was repeated for the desired number of rounds until all targets were imaged.

### *d*STORM data processing

#### Localization fitting and image reconstruction

After every single SMLM imaging round, the acquired image stacks were automatically processed by RapidSTORM software for 3D single molecule localization with local relative thresholding. The resulting localization data files were further post-processed by an in-house developed Post-Processing Software written in Java. Post-processing encompassed, correction of lateral drift that occurred during the 10 min acquisition, linking of blinking events that remained in the ‘on’ state for multiple consecutive frames, and image rendering. Depending on the purpose, 3D data was either rendered for a 2D (projected onto a plane) or a 3D representation using a pixel size (for 2D) or voxel size (for 3D) of 10 nm.

#### Fiducial-based registration

Fiducial coordinates were extracted from fiducial images acquired after each *d*STORM session by RapidSTORM fitting. The transformation matrix was calculated between fiducial coordinates from the first imaging round and any subsequent round (n > 1) of the same ROI. This matrix was then applied to correctly align the SMLM image from the n^th^ imaging round with respect to the first imaging round. The transformation was done either manually as described in our previous publication^18^ or with our newly developed 3D Transformer software.

### *d*STORM data ah3nalyses

*Nucleus.* In 400 nm brain tissue sections, the identification of the nucleus of the principal cell was achieved through the detection of autofluorescence signals within the 488 nm laser channel, a method previously validated using WGA labeling of nuclear pores (N > 10 experiments, available upon request). Comparative analysis revealed that the nuclear area exhibits considerably reduced signal intensity relative to the cytoplasmic region. For comparative signal quantification, a 3500 nm × 3500 nm window was randomly placed in the nuclear area. The number of localizations was quantified in ImageJ from *d*STORM images rendered with intensity values reflecting the number of localizations. Rendering was done with our custom-written post-processing software. Density of clusters with the radius above the calculated resolution (>28 nm) was quantified using the ‘Analyze particles’ plugin of ImageJ.

#### Endoplasmic reticulum

To quantify the localization density, 400-1000 nm long and 100-150 nm wide selections were positioned with their centers matching the center of the PDI signal along ER strands. The signal density was quantified by dividing the number of localizations by the selection area. Signal outside the ER was determined by placing selections in areas devoid of PDI signal and excluding lysosome and Golgi regions. For colocalization analysis, images rendered with a 10 nm pixel size were blurred with a sigma of 2. Correlation between lectins and PDI in the ER selections was determined using the Pearson’s analysis in ImageJ. For the signal distribution analysis across ER tubules, 400 nm wide line profiles were laid perpendicularly to ER strands. Multiple line profiles were aligned according to the PDI peak and the averaged line profiles with standard deviations were calculated and plotted. Gaussian fitting was performed to determine the full width at half maximum values.

#### Golgi apparatus

To analyze the distribution of multiple conventional antibodies and lectins, rendered images (10 nm pixel size) were analyzed in ImageJ. The combination of GM130 and TGN signals allowed us to identify areas of the Golgi apparatus displaying the *cis*—*trans* distribution of markers across the imaging plane. Line profiles, 300 nm wide and 1600 nm long, were drawn across these Golgi regions perpendicularly to the GM130 signal, (see illustration in Fig. 3c). The intensity plot values were extracted from ImageJ and transferred to the customized MatLab script to obtain averaged line profiles. Alignment of multiple line profiles was realized based on the GM130 signal. For the averaged line profile plot, the global and local peaks were directly extracted from the averaged smoothed curve. For the bar plots, global and local peaks were determined from line profiles that were grouped and averaged per calyx of Held.

#### PLCs/lysosomes

For the line profile, a 2D rendering of the lysosome was analyzed using an ImageJ script where a line selection with a width of 50 nm was rotated in increments of 30°, resulting in five line profiles that were averaged to create the intensity curve plot. For the analysis of correlation between PWA and cathepsin D signal, wide-field images of brain tissue sections were used. Tissue sections were stained with PWA-AF647 and cathepsin D primary antibody and a AF488-labeled secondary antibody. Round selections with an area of 1.15 µm² were centered around the center of the PWA signal and the signal was quantified in ImageJ. To determine the signal in PWA-negative areas, the selections were placed in regions devoid of obvious PWA signal next to the PWA-positive areas. For quantification of localization density, lysosome areas were selected based on the PWA signal. Localization density was quantified from 2D-projected STORM images rendered with intensities correlated to the number of localizations.

#### Synaptic specializations

Quantification of lectin signals at synaptic contact sites was performed in hippocampal neurons. Synaptic contacts were chosen based on Homer1b/c and Bassoon signals and two groups were collected for analyses: contacts imaged in the *en face* orientation where Homer1b/c and Bassoon are superimposed in the x/y view and contacts imaged in profile with a visible gap between Homer1b/c and Bassoon signals. For the analysis of the enrichment of lectins at synaptic contacts, only *en face* synaptic contacts were used. In ImageJ, the synaptic contact area was segmented based on the Homer1b/c signal in 2D renderings. The number of localizations in these regions was measured for the antibodies and the lectin. To quantify the specificity of lectin binding to synaptic contacts, two further area categories were established: areas outside the AZ/PSD region within neuronal processes and areas outside the neuron (at the glass-bottom surface of the dish). The selections previously segmented based on Homer1b/c signal were placed directly besides the synaptic contact within the neuronal process or outside the neuron, resulting in an equal number of analyzed regions in all three categories. The analysis of the spatial distribution of lectins between the AZ and the PSD was conducted on synaptic contacts imaged in profile. Line selections of 300 nm width were placed across the synaptic contact. Intensity profiles were extracted from ImageJ and averaged in the custom-written Matlab script as described for the Golgi analysis.

*Imaging quality evaluation.* Localization precision was determined from the first 10,000 frames of the raw localization data using the nearest neighbor analysis in LAMA-software^54^. Spatial resolution was quantified on fiducial-free *d*STORM image areas through decorrelation analysis^21^. Images were rendered based on intensity counts from all acquired frames (20,000), with both the pixel size and the sigma set to 10 nm.

## Data availability

Source data tables underlying all quantifications shown in this paper are provided as a Source Data files. All raw and processed images used in this work will be provided upon request.

## Code availability

This study did not involve the development of new relevant code. All relevant code utilized was previously published and publicly accessible^18^.

## Supporting information

Supplementary Discussion

## Acknowledgements

We thank Claudia Koksch, Franziska Gleiche, and Michaela Kaiser for their excellent technical assistance. We thank Dr. Frank Hermannsdoerfer, Rory Power, and Gerald Bendner for their help with the realignment of the optical path of the maS³TORM setup. We thank Aleksandar Stojic for kindly providing additional test tissue samples. The authors gratefully acknowledge the data storage service SDS@hd supported by the Ministry of Science, Research and the Arts Baden-Württemberg (MWK) and the German Research Foundation (DFG) through grant INST 35/1503-1 FUGG.

## Author contributions

M.K. conceptualized this work and designed all experiments with advice from T.K. and M.H.. M.K. and H.S. wrote the manuscript with contributions from T.K. and M.H.. M.K. did all animal perfusion experiments and created 400 nm sections. H.S. and M.K. collected and analyzed the data. M.K. wrote the analysis algorithms. S.S. established the protocol for the application of nanodiamond fiducials and provided STORM images from brain tissue sections that demonstrate synaptic performance of WGA and PSA. All authors edited, read, and approved the manuscript.

## Supplementary information

This paper contains extended data, including supplementary figures, tables, and discussion.

