## Supplementary Figures for "Glyco-STORM: Nanoscale Mapping of the Cellular Glycosylation Landscape"

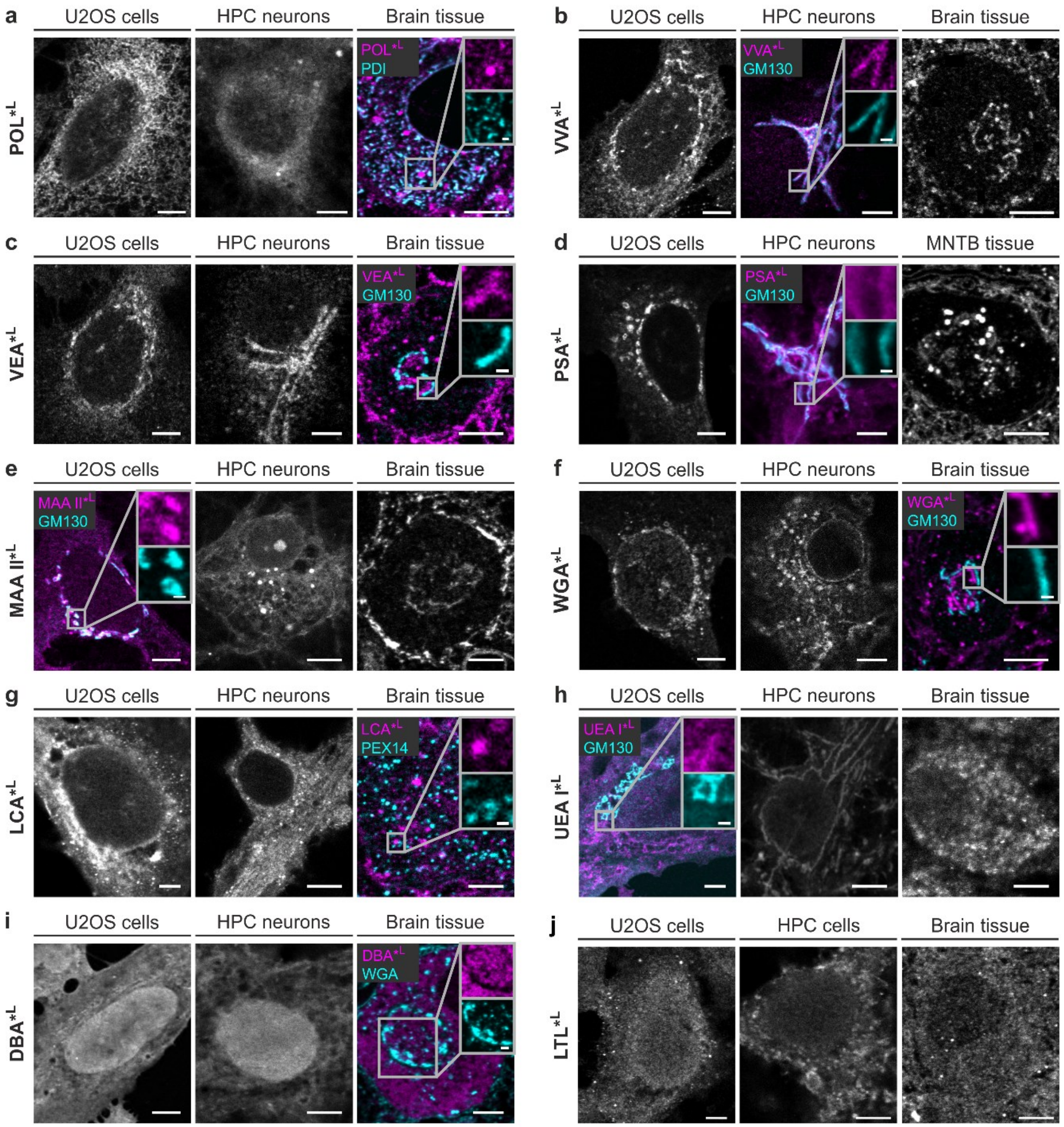


**Figure S1 | Extended results of lectin screening at the confocal microscope.** Lectin staining in U2OS cells, hippocampal (HPC) neurons and brain tissue. **a,** POL colocalizes with endoplasmic reticulum marker PDI. **b-f,** VVA, VEA, PSA, MAA II, and WGA exhibit staining at or close to the Golgi apparatus, as confirmed by spatial proximity with *cis*-Golgi maker GM130. **g,** LCA shows staining of small round clusters that do not colocalize with peroxisome marker PEX14. **h,** UEA I displays filamentous staining in hippocampal neurons and U2OS cells with only sparse signal in GM130-positive areas. No filamentous signal was observed in brain tissue. **i,** DBA shows abundant nuclear and cytosolic staining but is clearly absent in the Golgi area in principal cells in brain sections. **j,** LTL displays broad, partially clustered nucleocytoplasmic staining. Scale bars correspond to 5 µm in all overview images and 1 µm in the insets.

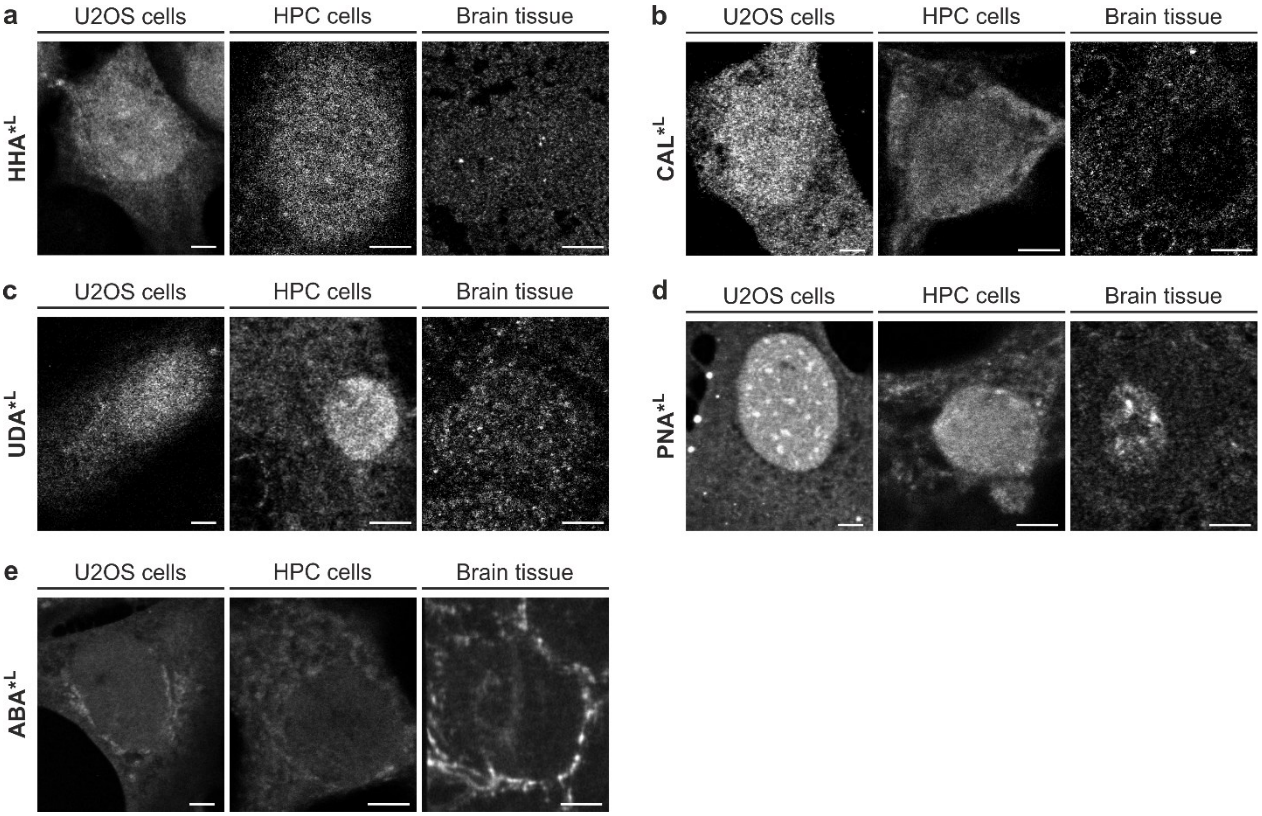


**Figure S2 | Lectins with weak signals or variable performance. a-c,** HHA, CAL, and UDA showed diffuse cytoplasmic and nuclear staining in most cell/tissue models, often with low-intensity signals. **d,e,** Image panels represent the predominant staining patterns for PNA and ABA, which varied across experiments. Scale bars correspond to 5 µm.

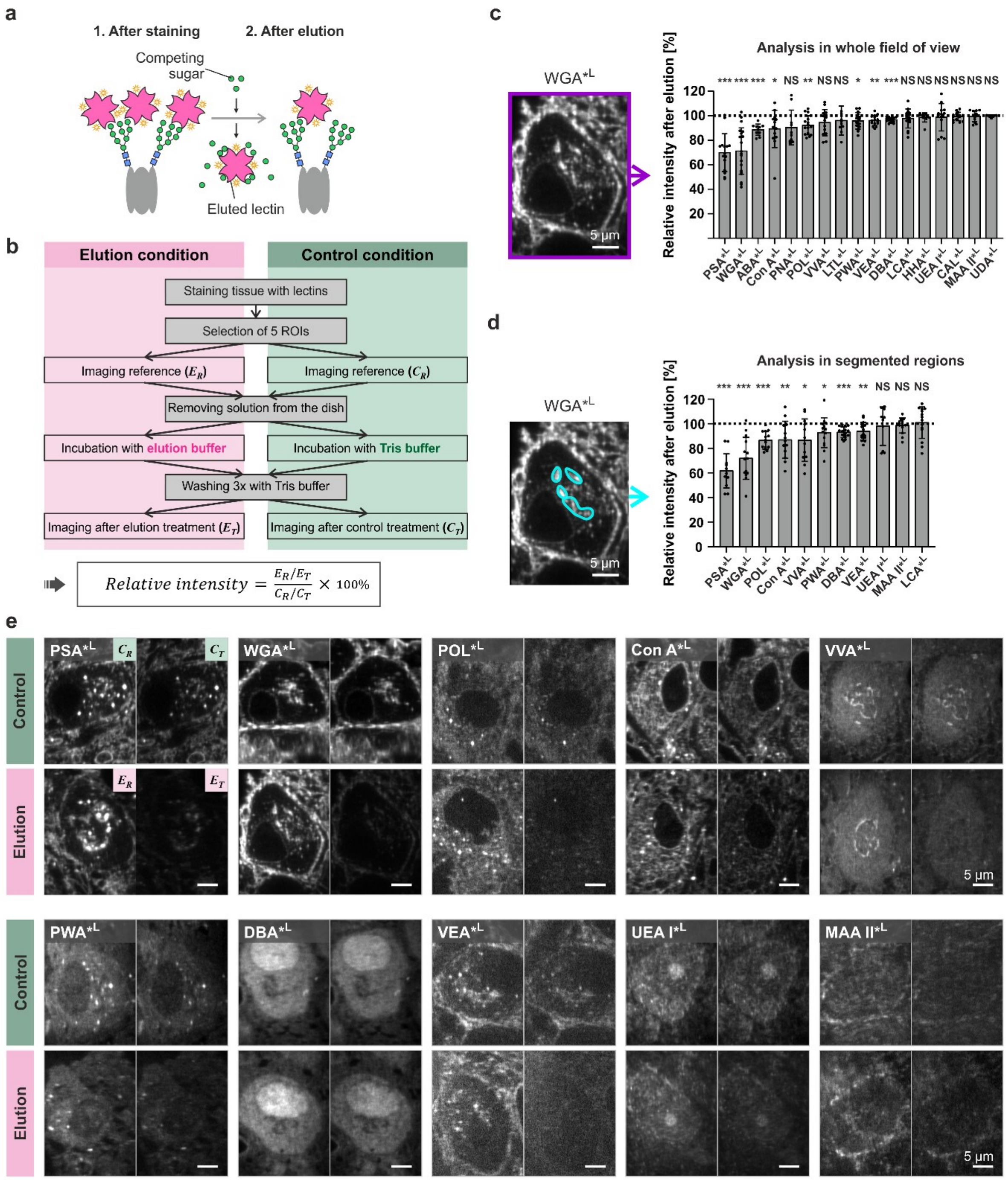


**Figure S3 | Characterization of lectin-specificity by elution experiments. a,** Elution experiments using sugars competing with carbohydrates recognized by the respective lectins were performed to characterize lectin specificity. Example illustrating elution of a mannose-binding lectin using mannosides. **b,** Experimental workflow for elution experiments. **c-d,** Signal intensity was either quantified in full-frame images containing whole cells (**c**) or in specifically segmented cell areas (**d**). Data represent mean intensity percentage ±SD from $n = 5$ to 15 analyzed principal cells per lectin. Asterisks denote statistical significance (***$p \leq 0.0001$, **$p \leq 0.01$, *$p \leq 0.05$, NS = not significant; two-sided one sample t-test test versus 100%). **e,** Images showing loss of signal after elution with competing sugars compared to control experiments visualizing the effect of bleaching and washing without sugar treatment (used as 100% reference).

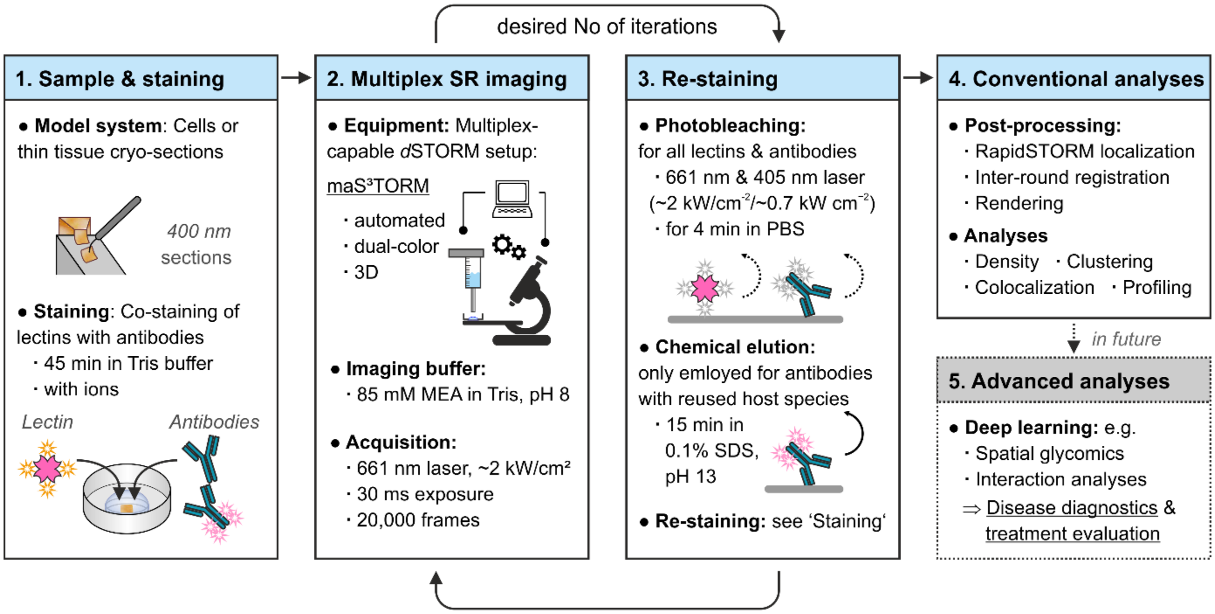


**Figure S4 | Glyco-STORM concept.** Strategy for Glyco-STORM imaging. (1) Staining of thin tissue or cells with lectins is realized in Tris buffer supplemented with ions and can be done together with classical immunohistochemistry stainings. (2) Automated 3D super-resolution (SR) imaging and re-staining of samples is realized using the maS³TORM setup that allows dual-channel acquisition of two spectrally close far-red fluorophores (in our case Alexa fluor 647 and CF647). Any other STORM setup and re-staining method can be used. (3) For re-staining, signals from lectins and antibodies imaged in the previous round are removed by photobleaching. Photobleaching causes quenching of all fluorophores and is expected to also result in photo-unbinding of a certain fraction of fluorophore-conjugated labels[1],[2] (dashed curved arrow). Chemical elution results in reliable removal of antibodies and is only realized after rounds preceding the usage of secondary antibodies against antibodies from host species already employed in previous rounds (solid curved arrow). (4) Data undergoes post-processing, including localization fitting, fiducial-aided registration of images from different rounds, and image rendering. Classical analysis methods suitable for super-resolution data can be applied. (5) Future directions: With the accumulation of sufficient data amenable to deep learning techniques, spatial glycomics and interaction analyses will facilitate the comparison of glycosylation profiles of diseased versus healthy, or treated versus untreated samples.

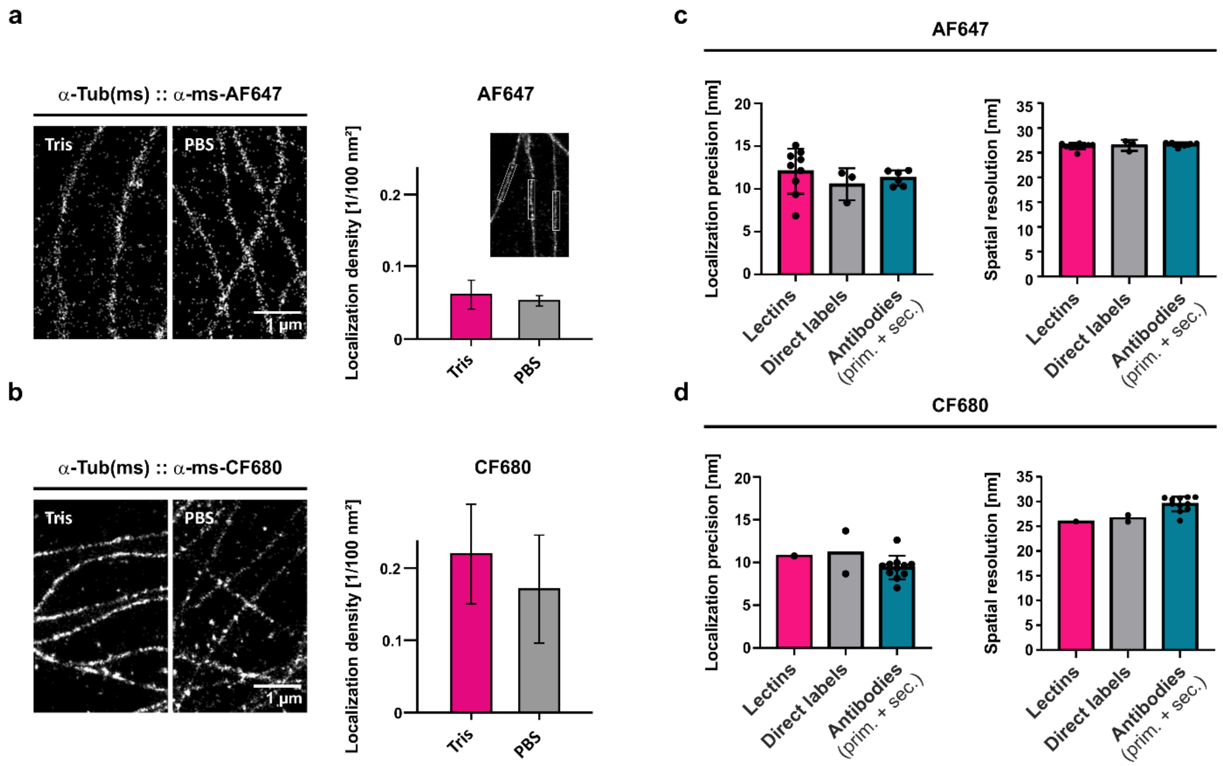


**Figure S5 | Buffer optimization and evaluation of imaging precision for Glyco-STORM. a,b,** Evaluation of Tris- versus PBS-based buffers for immunohistochemical staining and STORM imaging. Tris-based buffers (staining: 50 mM Tris-HCl, 150 mM NaCl; imaging: 85 mM MEA in Tris, 35 mM KOH, pH 8) were used for Glyco-STORM to prevent ion precipitation in PBS (staining: PBS; imaging: 100 mM MEA in PBS, 20 mM KOH, pH 8). The effectiveness of Tris-based buffers was validated through classical immunohistochemical stainings, demonstrating similar STORM image quality and localization density to PBS-based buffers, previously employed for maS³TORM[2]. Buffers were tested using the anti-α-Tubulin primary antibody from mouse and (**a**) the anti-mouse-AF647 or (**b**) the anti-mouse-CF680 secondary antibody. Bars represent mean density ±SD, $n$ = 10 to 36 quantified microtubule segments (exemplified in the inset in **a**). **c,d,** Lectins result in localization precision (left panels) and spatial resolution (right panels) comparable to direct labels such as phalloidin or directly labeled primary antibodies and nanobodies. Each data point corresponds to a distinct label and is the average value from 3 to 5 analyzed STORM images per label. Bar graphs represent mean ±SD.

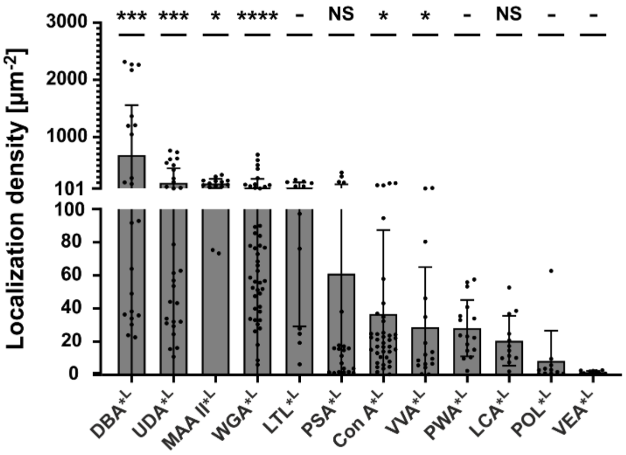


**Figure S6 | Signal density of lectins in the nucleus.** Density of localizations within the nuclear area. Bar plots represent means ±SD from 3 to 8 independent experiments per lectin (3 ≤ *N* ≤8) and 3 to 8 measured areas per experiment (3 ≤ *n* ≤8). Asterisks denote statistical significance (****p ≤ 0.0001, ***p ≤ 0.001, *p ≤ 0.05, NS = not significant; Mann-Whitney U test) versus nonspecific signal on glass surface (not shown for the sake of clarity). Dashes (-) indicate that no reference for nonspecific signal is available for the respective lectins due to the absence of regions lacking tissue in these experiments.

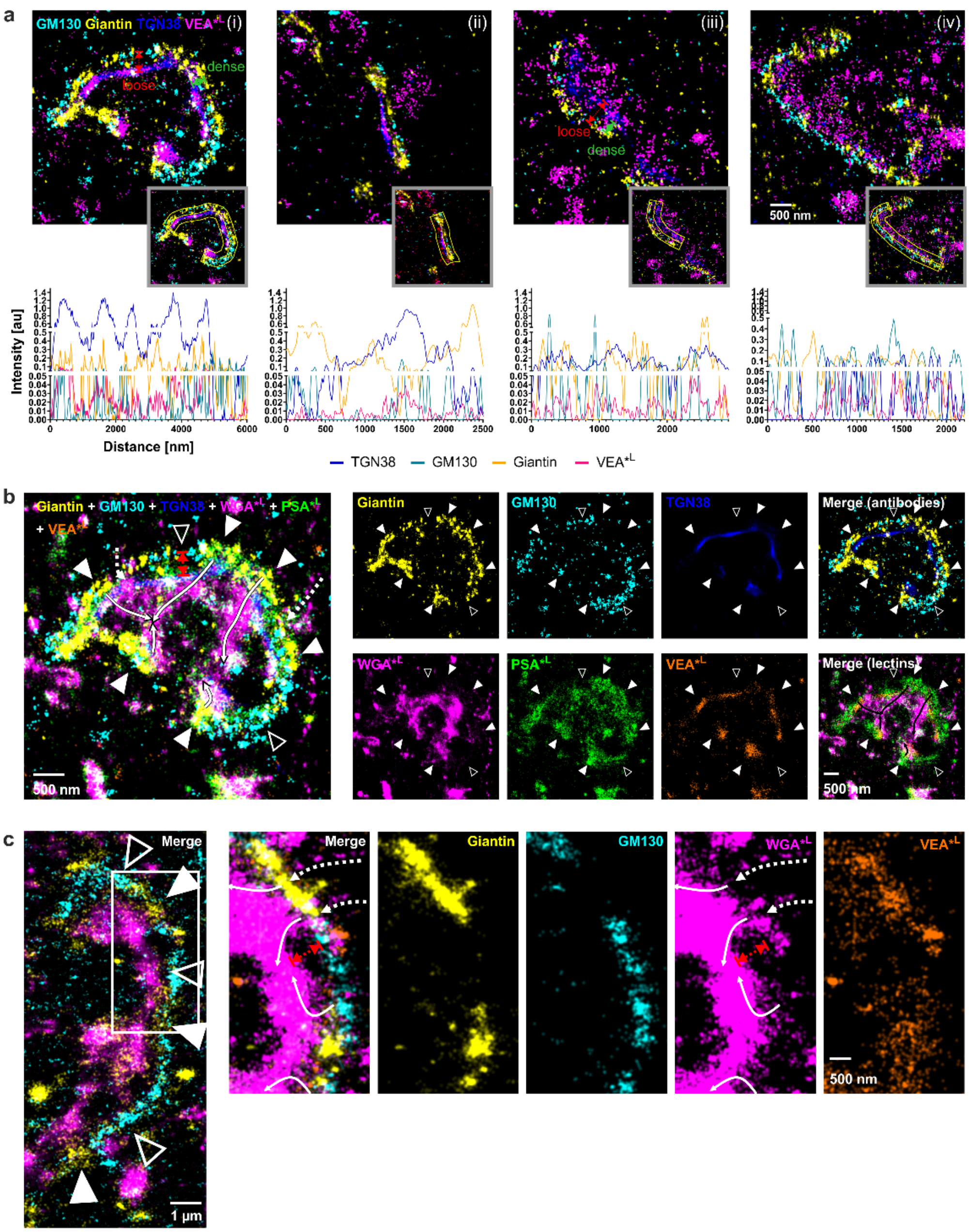


**Figure S7 | Lateral Golgi organization in brain tissue neurons. a-c,** Additional examples illustrating the lateral periodic distribution of Golgi antibody markers GM130, Giantin, and TGN38 and lectins. GM130 and Giantin signals are periodically alternating along the lateral Golgi ribbon axis (open and filled arrowheads in **b** and **c**). In many experiments, VEA signal positively correlates with the Giantin signal. **a,** Top row shows example images, while the bottom row presents line profiles taken along the lateral Golgi axis (shown in insets of the top-row images). Often Golgi regions overarched by Giantin are more densely packed along the *cis*-to-*trans* axis (denoted by green double-headed arrows between GM130 and TGN38 markers in i and iii) compared to the more loose packaging in GM130-rich regions with low or absent Giantin content (red double-headed arrows). **b,c,** Two examples illustrating how WGA-positive vesicles could transverse Giantin-overarched regions (dashed and solid white curved arrows), eventually merging with the TGN region. We propose that lectins such as VEA and WGA that appear enriched in Giantin-enclosed Golgi areas. These areas could represent non-compact Golgi regions and potentially facilitating bidirectional transport that bypasses Golgi stacks.

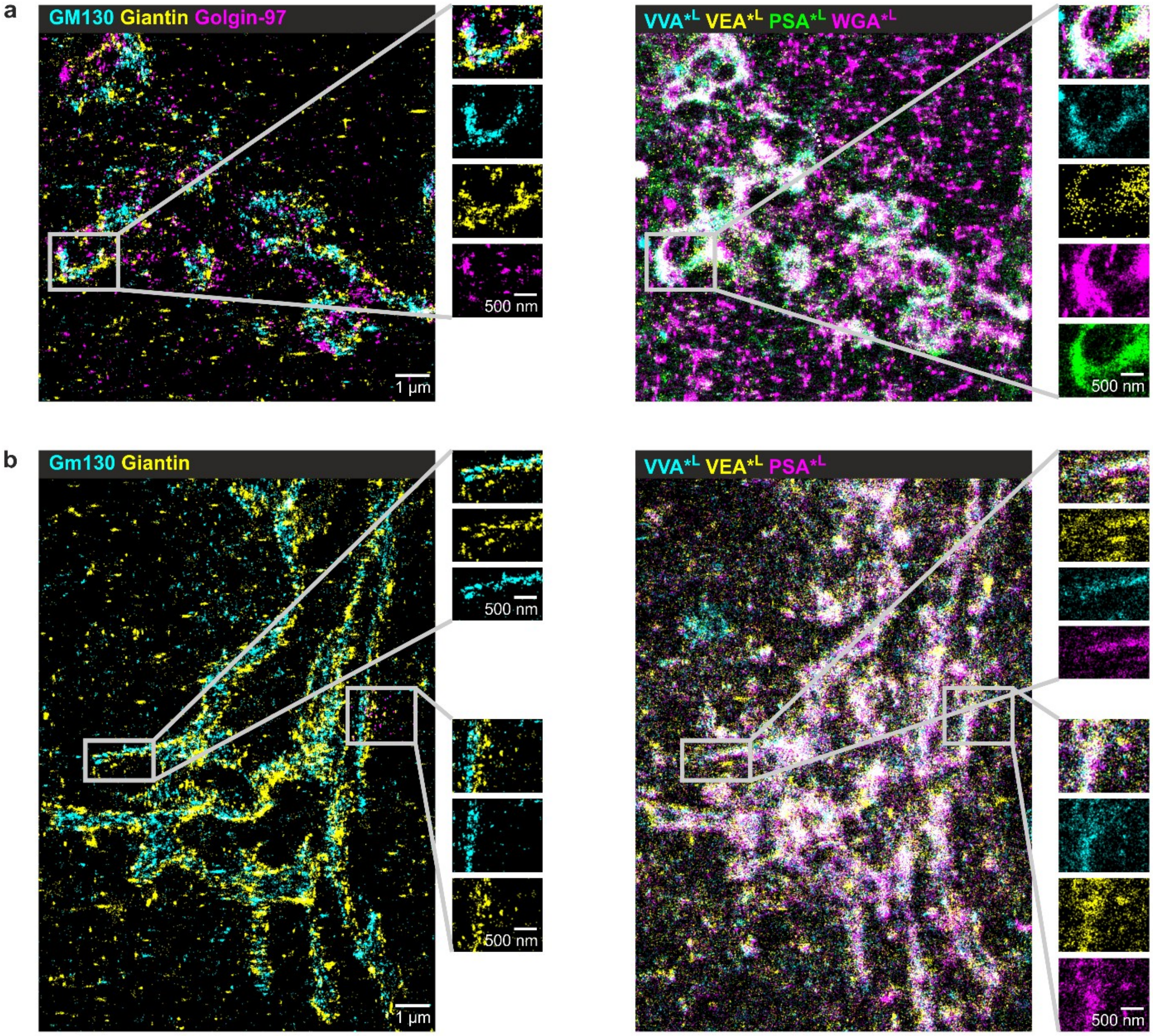


**Figure S8 | Golgi in U2OS and hippocampal neurons.** Golgi apparatus ultrastructure visualized by conventional antibody markers and lectins in **a**, human osteosarcoma cells (U2OS) and **b,** hippocampal neurons.

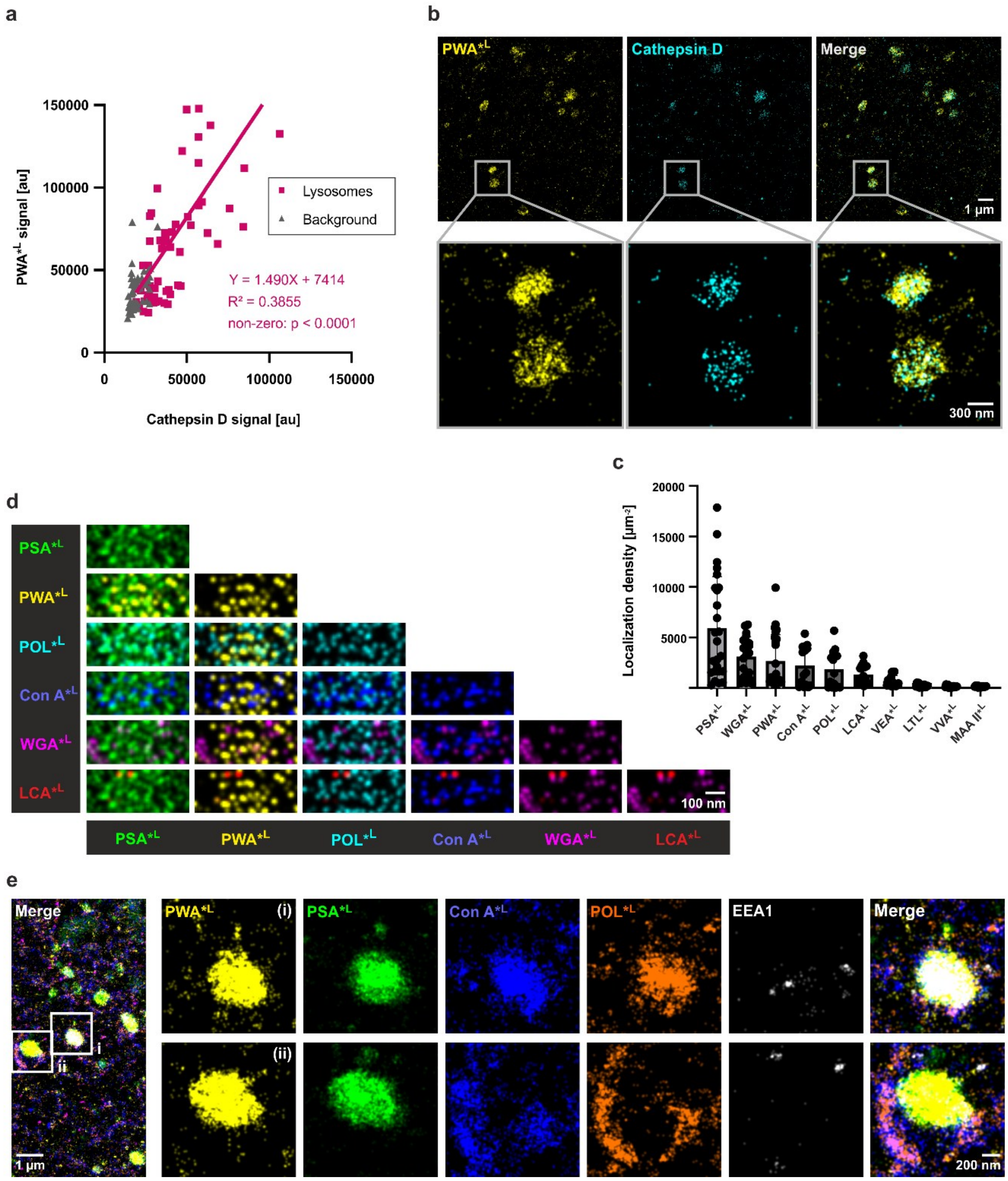


**Figure S9 | Pre-lysosomal compartments/lysosomes in brain tissue cell somata. a,** PWA signal correlates with pre-lysosomal compartment (PLC) and lysosome marker cathepsin D. Clusters were chosen based on areas enriched for PWA signal from wide-field images. Total number of selections $n = 59$ from 5 experiments ($N = 5$). **b,** Glyco-STORM image illustrating that PWA is specifically present in lysosomes identified by the cathepsin D antibody. However, PWA and cathepsin D nanodomains do not overlap. **c**, Density of localizations per lysosome quantified from *d*STORM images. Number of selections and individual experiments: PWA, PSA – $n = 25$, $N = 5$; Con A, POL, LCA, VEA, LTL, VVA, MAA II – $n = 15$, $N = 3$; WGA – $n = 20$, $N = 4$. **d,** Pairwise merging of lectin signals from an optical cross-section through a 3D image of the lysosome. **e,** Two examples of PWA-positive PLCs/lysosomes, showing distinct staining patterns of the mannose-binding lectins Con A and POL: (i) lumenal signal within the organelle and (ii) surrounding signal that forms a clasp-like structure around the organelle.

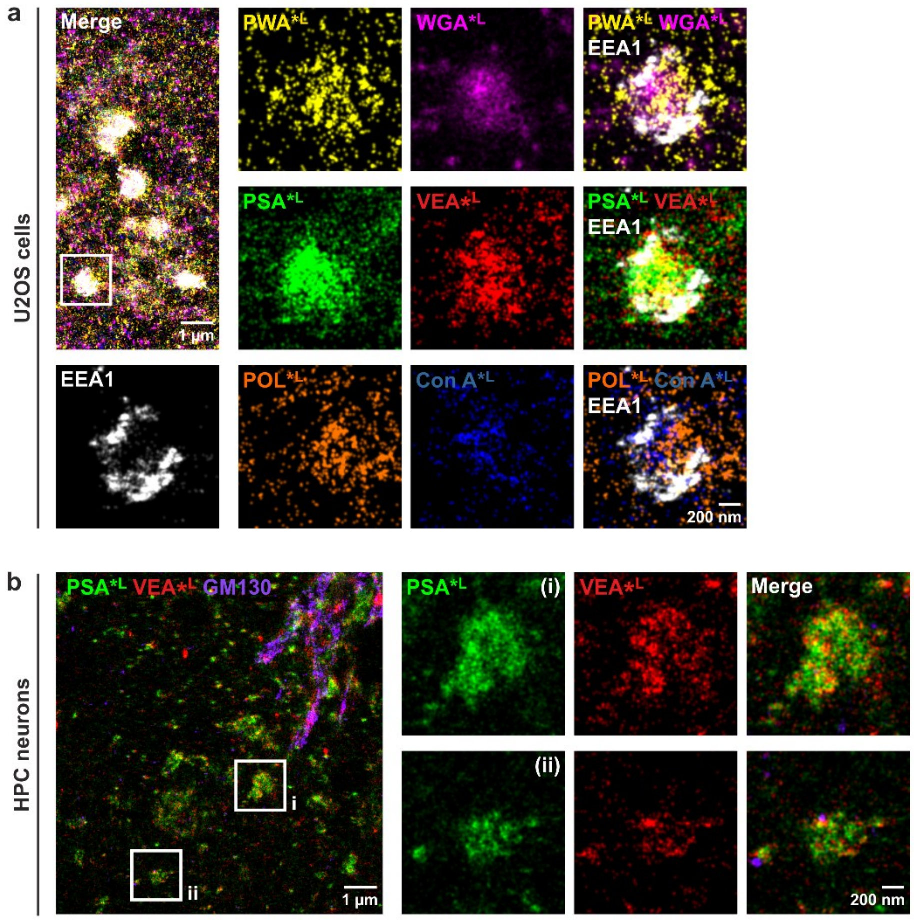


**Figure S10 | Glyco-STORM of pre-lysosomal compartments/lysosomes in U2OS and hippocampal cells. a,** In U2OS cells, multiple lectins stained large spherical organelles, with most of these organelles being abundantly coated by the early endosome marker EEA1. **b,** In hippocampal (HPC) neurons, spherical organelles of different sizes, located in areas distinct from the main Golgi body indicated by GM130 staining, were positive for PLC/lysosome-binding lectins PSA and VEA.

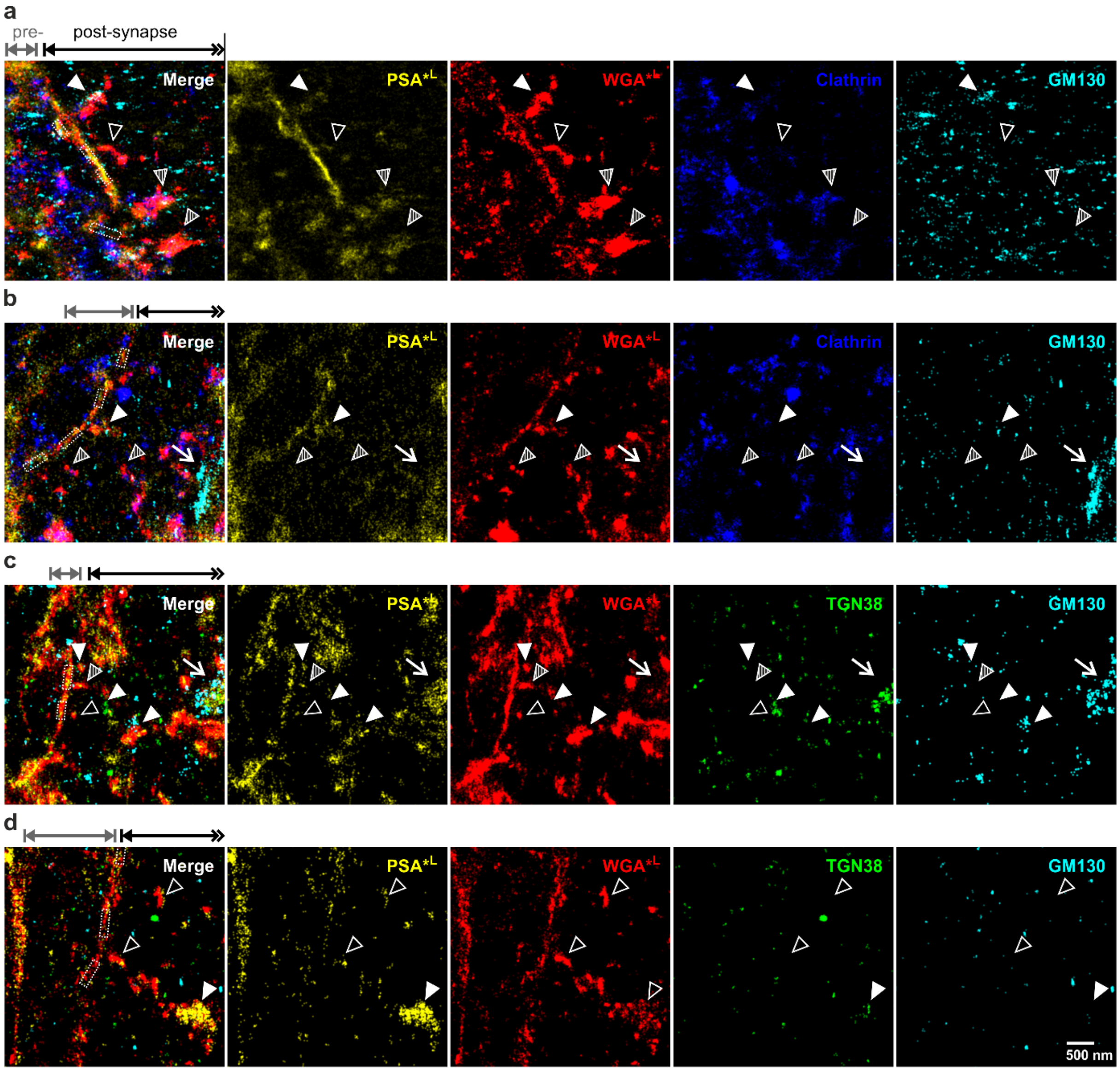


**Figure S11 | Tubular glycan-rich perisynaptic structures in the postsynaptic principal cell of the giant glutamatergic synapse in brain tissue. a-d,** Examples of postsynaptic glycans in proximity to synaptic contacts (dashed boxes; approximated based on Homer signal, not shown here). Some peri-synaptic glycan structures contain traces of Golgi markers GM130 and/or TGN38 (filled arrowheads), while other do not coincide with these Golgi markers (open arrowheads). Many peri-synaptic glycan structures also contain clathrin traces (**c**,**d**). Arrows point to Golgi marker signal the main Golgi apparatus whose parts are included in some images (**b**,**c**).

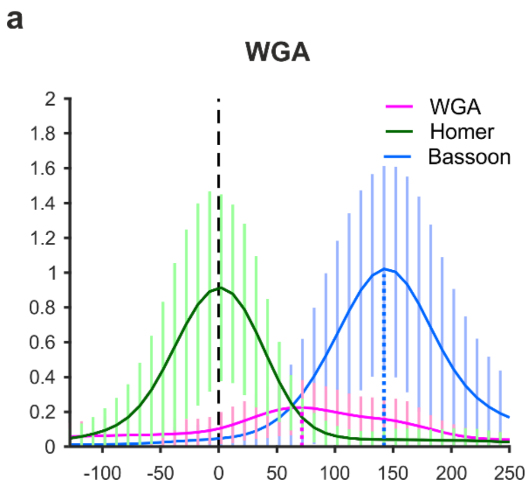


**Figure S12 | Distribution of glycans at the synaptic cleft of hippocampal neurons.** Averaged line profiles demonstrating that both WGA (**a**) and PSA (**b**) peak in the center between the active zone and the postsynaptic density. Line profiles for Homer and Bassoon were averaged from $n$ = 31 intensity profiles that are in turn averages for 3-10 synaptic interfaces per image acquired in $N$ = 12 individual experiments ($\eta$ = 110 synaptic interfaces in total); WGA: $\eta$ = 91, $n$ = 22, $N$ = 8; PSA: $\eta$ = 32, $n$ = 19, $N$ = 5. Vertical lines represent SD. The plotted peaks of the averaged profiles are displayed in the main Fig. 4h.

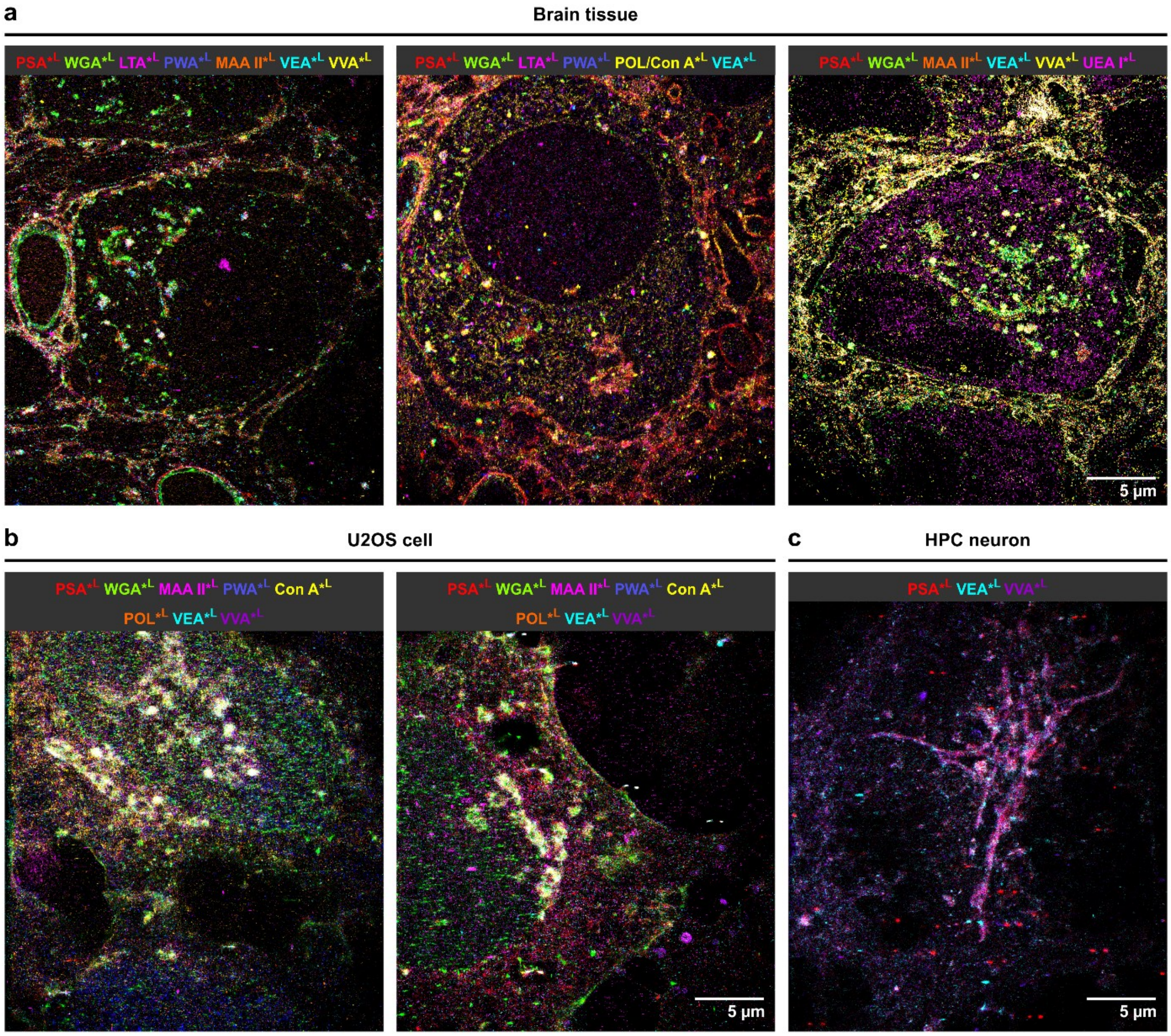


**Figure S13 | Glycosylation maps. a,** Examples from three individual experiments conducted with different lectin subsets in neuronal tissue. **b,c,** Exemplary glycosylation landscapes of two U2OS cells (**b**) and a hippocampal neuron (**c**).

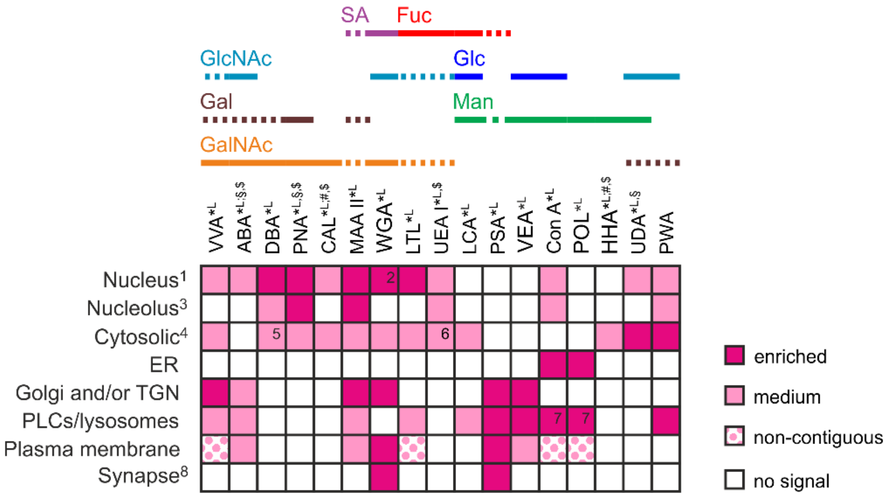


**Figure S14 | Performance matrix for all screened lectins.** Lectin performance matrix for 17 tested lectins, including those with low-intensity signals (#), variable performance ($), and those that were evaluated predominantly based on confocal screening (§). Note that a subset of lectins shown here is also highlighted in Fig. 5b as the best-studied examples tested by Glyco-STORM. [1]Nucleus staining refers to clustered or dispersed signal in the nucleoplasm if not stated otherwise. [2]In addition to the clustered signal in the nucleoplasm, WGA stains the nuclear pores. [3]Nucleolar signal was denoted as present if detected in a subset of cells. [4]Cytosolic localization refers to clustered or dispersed signal patterns. [5]DBA showed abundant cytosolic signal sparing distinct areas, including the Golgi apparatus. [6]UEA I showed streaky signal in U2OS cells and hippocampal neurons and clustered round signal in the principal cell soma in brain tissue. [7]For some pre-lysosomal compartments/lysosomes, Con A and POL showed signals within the lumen of the organell, while in others their signals were located at the periphery. [8]Synaptic localization refers only to brain tissue model. Solid lines highlight predominant sugar specificity; dashed lines denote sugar moieties that are primarily bound as part of a glycan motif. Abbreviations: ER, endoplasmic reticulum; Fuc, fucose; G1al, galactose; GalNAc, N-acetylgalactosamine; Glc, glucose; GlcNAc, N-acetylglucosamine; Man, mannose; PLCs, pre-lysosomal compartments; SA, sialic acid; TGN, *trans*-Golgi network.
