## Supplementary Tables for "Glyco-STORM: Nanoscale Mapping of the Cellular Glycosylation Landscape"

### Supplementary Table 1 | Properties of lectins used in this study

| **Lectin;**<br>**Organism** | **Size [kDa];**<br>**Subunits** | **Preferred binding motifs** |
|---|---|---|
| ABA;<br>*Agaricus bisporus* | 60; 4[1] | N-glycans: biantennary, β-GlcNAc terminated[2];<br>N-glycans: biantennary, LacNAc terminated[2];<br>O-glycans: core 2 (GlcNAcβ1-6(Galβ1-3)GalNAc)[2] |
| CAL;<br>*Cicer arietinum* | 43; 2[3] | N-glycans: complex carbohydrates[3];<br>GalNAc[4] |
| Con A;<br>*Canavalia ensiformis* | 104; 4[5] | N-glycans: α-Man, terminal ($Man_3$ to $Man_9$)[2];<br>N-glycans: biantennary with most extensions (no terminus-proximal α-Fuc)[2] |
| DBA;<br>*Dolichos biflorus* | 111; 4[6] | Forssman antigen (GalNAcα1-3GalNAcβ1-3Galα1-4Galβ1-4Glc)[2,7]; GalNAcα1-3GalNAc[7];<br>β-GalNAc[2]; α-GalNAc[7,8];<br>GalNAcα1-3(Fucα1-2)Gal[7]; |
| HHA;<br>*Hippeastrum hybrid* | 48; 4[9] | N-glycans: Man, terminal: Manα1-6(Manα1-3)Man[2];<br>$Man_5$>$Man_6$>$Man_7$[2] |
| LCA;<br>*Lens culinaris* | 48; 4[9] | N-glycans: Fuc, core (Fucα1-6)[2];<br>Man, terminal (Manα1-2)[2] |
| LTL;<br>*Lotus tetragonolobus* | 105; 4[10] | $Lewis^{X}$: Galβ1-4(Fucα1-3)GlcNAc[2];<br>Fucα1-3[2];<br>$Lewis^{y}$: Fucα1-2Galβ1-4(Fucα1-3)GlcNAc[2] |
| MAA II;<br>*Maackia amurensis* | 130; 4[11] | O-Glycan: SA α2-3-linked to Galβ1-3GalNAc[2];<br>3' sulfated β-Gal[2] |
| PNA;<br>*Arachis hypogaea* | 110; 4 | O-Glycan: Galβ1-3GalNAc, terminal[2]; |
| POL;<br>*Polygonatum odoratum* | 48; 4[12] | N-glycans: $Man_3$ (Manα1-3(Manα1-6)Man[12]);<br>$Man_2$>$Man_1$[12] |
| PSA;<br>*Pisum sativum* | 50; 4[13],[14] | Fuc, core (Fucα1-6), tolerating SA, Gal, GlcNAc[2];<br>Man[15] |
| PWA;<br>*Phytolacca americana* | 32; 1[16] | Chitin oligomers $(GlcNAc\beta1\text{-}4)_n$ (n ≥ 4), terminal[2]<br>poly-LacNAc[17] |
| UDA;<br>*Urtica dioica* | 9; 1[18] | Manα1-6, terminal $Man_3$ to $Man_9$, with chitobiose core[2];<br>(poly-)LacNAc[2];<br>Chitin oligomers $(GlcNAc\beta1\text{-}4)_n$[2] |
| UEA I;<br>*Ulex europaeus* | 63; 2[19] | Type 2 blood group H (Fucα1-2Galβ1-4GlcNAc)[2];<br>Fuc (Fucα1-2Galβ1-4Glc)[2];<br>$Lewis^{y}$ (Fucα1-2Galβ1-4(Fucα1-3)GlcNAc)[2] |
| VEA;<br>*Vicia ervilia* | 60; 4[20] | α-Man[20];<br>α-Glc[20,21] |
| VVA A+B;<br>*Vicia villosa* | 136; 4 &<br>144; 4[9] | GalNAcβ1-4GlcNAc[2];<br>α/β-GalNAc, terminal[2];<br>LacNAc, terminal, multiantennary[2] |
| WGA;<br>*Triticum aestivum* | 36; 2[22] | β-GlcNAc, terminal[2]; α-GlcNAc, terminal[2];<br>α/β-GalNAc[2]; SA: α-Neu5Ac[24Glc] |

The first entry for each lectin indicates the predominant binding motif. Abbreviations: Fuc, fucose; Gal, galactose; GalNAc, N-acetylgalactosamine; Glc, glucose; GlcNAc, N-acetylglucosamine; LacNAc, N-acetyllactosamine; Man, mannose; Neu5Ac, N-acetylneuraminic acid; SA, sialic acid.

**Supplementary Table 2 | Lectin-fluorophore conjugates used in this study**

| Lectin | Conju-gate | Used dilution | Recommended/ applied ions | Order № | Lot № |
|---|---|---|---|---|---|
| ABA | AF647 | 1:20 | - | 21511446[a] | L20061809ZH |
| CAL | AF647 | 1:20 | $Ca^{2+}$, $Mg^{2+}$ | 21511589[a] | L20110609ZH |
| Con A | AF647 | 1:20 | $Ca^{2+}$, $Mn^{2+}$ | 21511462[a] | L20042403ZH |
| Con A | CF680 | 1:200 | $Ca^{2+}$, $Mn^{2+}$ | 29020[b] | 21C0720-1164044 |
| DBA | AF647 | 1:20 | $Ca^{2+}$, $Mg^{2+}$, $Mn^{2+}$, $Zn^{2+}$ | 21511470[a] | L20110605ZH |
| HHA | AF647 | 1:5 | - | 21511503[a] | L20090406ZH |
| LCA | AF647 | 1:20 | $Ca^{2+}$, $Mg^{2+}$ | 21511511[a] | L20110606ZH |
| LTL | AF647 | 1:20 | $Ca^{2+}$, $Mn^{2+}$ | 21511594[a] | L20110612ZH |
| MAA II | AF647 | 1:20 | - | 21511531[a] | L20110607ZH |
| PNA | AF647 | 1:20 | $Ca^{2+}$, $Mg^{2+}$ | 21511454[a] | L20110604ZH |
| POL | AF647 | 1:20 | $Ca^{2+}$, $Mn^{2+}$ | 21511595[a] | L20110613ZH |
| PSA | AF647 | 1:100 | $Ca^{2+}$, $Mn^{2+}$ | 21511543[a] | L20110611ZH |
| PWA | AF647 | 1:20 | - | 21511588[a] | L20110608ZH |
| UDA | AF647 | 1:100 | $Zn^{2+}$ | 21511581[a] | L20052206ZH |
| UEA I | DL649 | 1:20 | $Ca^{2+}$, $Mn^{2+}$, $Zn^{2+}$ | BL-21068[c] | ZD1130 |
| VEA | AF647 | 1:20 | - | 21511591[a] | L20110610ZH |
| VVA A+B | AF647 | 1:5 | $Ca^{2+}$, $Mn^{2+}$ | 21511590[a] | L21080900CRCR |
| WGA | CF680 | 1:1000 | $Ca^{2+}$ | 29029[b] | 20W0527-1122158 |

Lectins were purchased as fluorophore conjugates from: [a]BioWORLD, Dublin, OH, USA, [b]Biotium, Fremont, CA, USA, or [c]Vector laboratories, Burlingame, CA, USA.

**Supplementary Table 3 | Primary antibodies and labels used in this study**

| Antibody/ Label | Sub-type | Host | Clonality, № | Used dilution | Provider | Order № | Lot № |
|---|---|---|---|---|---|---|---|
| **Fluorophore-conjugated primary antibodies or other direct labels** | | | | | | | |
| CHC17-AF647 | IgG1 | ms | mono, X22 | 1:30 | Novusbio | NB300-613AF647 | VL315162-070121-AF647 |
| DAPI | toxin | - | - | 1:5000 of 50 mg/ml | Invitrogen | D1306 | - |
| Hoechst-JF646 | bisbinzi-midine | - | - | 100 pM | Luke Lavis | - | - |
| Phalloidin-AF647 | toxin | - | - | 1:80 | ThermoFisher | A22287 | 1750839 |
| Phalloidin-AF680 | toxin | - | - | 1:80 | ThermoFisher | A22286 | 1709895 |
| VGlut1-AF647 | sdAb | ll | mono, Nb9 | 1:200 | NanoTag | N1605 | 220602 |
| VGlut1-CF680 | sdAb | ll | mono, Nb9 | 1:200 | NanoTag | N1605 | 12190101 |
| **Unconjugated primary antibodies** | | | | | | | |
| Bassoon | IgG2aκ | ms | mono, SAP7F407 | 1:200 | Enzo Life Sciences | ADI-VAM-PS003 | 02012005 |
| Cathepsin D | IgG | rb | poly | 1:200 | Cell Signalling | 69854 | 1 |
| EEA1 | IgG | rb | poly | 1:200 | Abcam | ab2900 | - |
| GLT1 | serum | gp | poly | 1:400 | Millipore | AB1783 | 3526283 |
| Fibrillarin | IgG | rb | poly | 1:200 | Abcam | ab5821 | GR293483-2 |
| Giantin | serum | gp | poly | 1:200 | SySy | 263004 | 263004/1 |
| GM130 | IgG1 | ms | mono, 35 | 1:200 | BD Biosciences | 610822 | 1145724 |
| Golgin97 | IgG | rb | poly | 1:70 | ThermoFisher | PA5-83719 | - |
| Homer1b/c | IgG | rb | poly | 1:400 | SySy | 160023 | 160023/2-11 |
| Lamin a/c | IgY | ch | poly | 1:200 | Biosensis | C1698-100 | C-1698-300-201307-SH |
| LAMP1 | IgG1 | ms | mono, H4A3 | 1:200 | Abcam | ab25630 | GR3395210-1 |
| LAMP1 | IgG | rb | poly | 1:200 | Abcam | ab24170 | GR3395210-1 |
| LAMP2A | IgG | gp | poly | 1:400 | SySy | 437005 | - |
| LAMP3 | IgG | gp | poly | 1:200 | SySy | 391005 | - |
| LAMP5 | IgG | gp | poly | 1:200 | SySy | 412005 | - |
| MAP2 | IgY | ch | poly | 1:500 | SySy | 188006 | 188006/1 |
| PDI | IgG1 | ms | mono, 1D3 | 1:400 | Enzo Life Sciences | ADI-SPA-891 | 03022019 |
| PEX14 | IgG | rb | poly | 1:200 | Novusbio | NPB2-33455 | A119064 |
| Piccolo | IgG | rb | poly | 1:200 | SySy | 142113 | 142113/1-3 |
| Rab3a | IgG1 | ms | mono, 42.2 | 1:200 | SySy | 107111 | 107111/1-20 |
| Rab5 | IgG | rb | poly | 1:200 | Abcam | ab18211 | 793878 |
| Rab7 | IgG | rb | mono, n.a. | 1:200 | Abcam | ab137029 | GR155792-64 |
| S100B | IgY | ch | poly | 1:400 | SySy | 287006 | 287006/1-4 |
| SCAMP1 | IgG | rb | poly | 1:200 | SySy | 121003 | - |
| Synapto-physin | IgG | rb | poly | 1:200 | SySy | 101002 | 101002/26 or 1-41 |

| | | | | | | | |
|---|---|---|---|---|---|---|---|
| Syntaxin6 | serum | rb | poly | 1:100 | SySy | 110062 | - |
| SV2 a/b/c | IgG1 | ms | mono | 1:200 | DSHB | AB2315387 | ABC214744 |
| TGN38 | IgG1 | ms | mono, Clone 2 | 1:100-1:200 | BD Biosciences | 610898 | 8274719 |
| TGN38 | serum | rb | poly | 1:100 | | NB1-03495SS | RB0705-060908-WS |
| VAMP1 | serum | rb | poly | 1:500 | SySy | 104002 | 104002/1-16 |
| VAMP2 | IgG | rb | mono | 1:500 | SySy | 104008 | 104002/1-5 |
| VAMP4 | serum | rb | poly | 1:100 | SySy | 136002 | - |
| VAMP7 | IgG | ms | mono, 158.2 | 1:100 | SySy | 232011 | 232011/1-8 |
| VGlut1 | serum | gp | poly | 1:100 | Millipore | AB5905 | 3836436 |
| VGlut2 | IgY | ch | poly | 1:200 | SySy | 135416 | 135416/1-6 |

Abbreviations: AF647/680, Alexa Fluor 647/680; ch, chicken; gp, guinea pig; JF646, Janelia Fluor 646; ll, llama; mono, monoclonal; ms, mouse; n.a., not available; poly, polyclonal; rb, rabbit; sdAB, single domain antibody (nanobody); SySy, Synaptic Systems.

**Supplementary Table 4 | Secondary antibodies used in this study**

| Target/ Conjugate | Sub-type | Appli-cation | Host | Used dilution | Provider | Order № | Lot № |
|---|---|---|---|---|---|---|---|
| anti-ms IgG/ AF488 | $F(ab')_2$ | WF | gt | 1:400 | Invitrogen | A11017 | 262519 |
| anti-ms IgG/ AF647 | IgG (H+L) | WF | gt | 1:400 | Invitrogen | A21235 | 2369184 |
| anti-ms IgG/ AF647 | $F(ab')_2$ | *d*STORM | dn | 1:400 | Abcam | ab181292 | GR3310174-2 |
| anti-ms IgG/ CF680 | IgG (H+L) | *d*STORM | gt | 1:400 | Sigma | SAB4600199 | 18C0130 |
| anti-rb IgG/ AF488 | IgG (H+L) | WF | gt | 1:400 | Invitrogen | A11008 | 2420731 |
| anti-rb IgG/ AF568 | IgG (H+L) | WF | gt | 1:400 | Invitrogen | A11011 | 2500544 |
| anti-rb IgG/ AF647 | IgG (H+L) | WF | gt | 1:400 | Invitrogen | A21245 | 264997 |
| anti-rb IgG/ AF647 | $F(ab')_2$ | *d*STORM | dn | 1:400 | Abcam | ab181347 | GR3361186-1 |
| anti-rb IgG/ CF680 | $F(ab')_2$ | *d*STORM | gt | 1:400 | Sigma | SAB4600362 | 16C0721 |
| anti-gp IgG/ AF488 | IgG (H+L) | WF | gt | 1:400 | Invitrogen | A11073 | 1990462 |
| anti-gp IgG/ CF680 | IgG (H+L) | *d*STORM | gt | 1:400 | Biotium | 20499 | 21C0104 |
| anti-ch IgY/ AF488 | IgG (H+L) | WF | gt | 1:400 | Invitrogen | A11039 | 2566343 |
| anti-ch IgY/ AF647 | IgG (H+L) | *d*STORM | gt | 1:400 | Invitrogen | A21449 | 1806124 |

All secondary antibodies used in this study were polyclonal. AF488/568/647, Alexa Fluor 488/568/647; ch, chicken; dn, donkey; gp, guinea pig; gt, goat; ms, mouse; rb, rabbit; WF, wide-field.

**Supplementary Table 5 | Composition of lectin elution buffers**

| Eluted lectin | Competing sugars | Order № |
|---|---|---|
| ABA | 0.2 M Gal-GalNAc[a] | 21511278-1 |
| CAL | 0.2 M Bovine fetuin[a] | 21511283-1 |
| Con A, HHA, PSA, LCA | 0.2 M α-Methylmannoside[a] | 21511337-1 |
| DBA | 0.2 M GalNAc[a] | 21511031-1 |
| MAA II | 0.2 M Lactose[a] | 21511261-1 |
| PNA | 0.2 M Galactose[a] | 21511342-1 |
| POL | 0.1 M Mannose[b] | 20120110-1 |
| PWA, WGA | 0.2 M Chitin hydrolysate[a] | 21511023-3 |
| UDA | 0.2 M Chitobiose[a] | 21511315-1 |
| UEA I, LTL | 0.2 M L-Fucose[a] | 21511316-1 |
| VEA | 0.2 M Glucose[a] | 21511318-1 |
| VVA | 0.1 M GalNAc[b] | 21510884-1 |
| WGA | 0.1 M GlcNAc[b] | 21510885-1 |

Competing sugars were purchased in following buffers:
[a] 10 mM Tris with 0.1 M or 1 M NaCl and 1 mM or 2 mM EDTA
[b] 1 M NaCl and 2 mM EDTA

**Supplementary Table 6 | Areas segmented for region-specific elution analysis**

| Eluted lectin | Segmented area |
|---|---|
| Con A | Endoplasmic reticulum, round organelles |
| DBA | Cytosol |
| LCA | Round organelles |
| MAA II | Golgi apparatus, plasma membrane |
| POL | Endoplasmic reticulum, round organelles |
| PSA | Golgi |
| PWA | Round organelles (lysosomes) |
| UEA I | Round organelles |
| VEA | Golgi apparatus |
| VVA | Golgi apparatus |
| WGA | Endoplasmic reticulum, round organelles, nuclear pores |

**Supplementary Table 7 | Number of experiments and confocal or wide-field images acquired during initial lectin screening and elution**

| Lectin | Brain tissue | Hippocampal neurons | U2OS cells |
|---|---|---|---|
| ABA | N = 7 | N = 2 | N = 2 |
| CAL | N = 2 | N = 2 | N = 1 |
| Con A | N = 22 | N = 7 | N = 4 |
| DBA | N = 12 | N = 4 | N = 6 |
| HHA | N = 4 | N = 1 | N = 1 |
| LCA | N = 10 | N = 3 | N = 5 |
| LTL | N = 7 | N = 3 | N = 5 |
| MAA II | N = 21 | N = 3 | N = 9 |
| PNA | N = 5 | N = 2 | N = 7 |
| POL | N = 11 | N = 8 | N = 3 |
| PSA | N = 16 | N = 13 | N = 11 |
| PWA | N = 11 | N = 5 | N = 9 |
| UDA | N = 6 | N = 4 | N = 1 |
| UEA I | N = 7 | N = 4 | N = 4 |
| VEA | N = 17 | N = 12 | N = 11 |
| VVA A | N = 21 | N = 12 | N = 11 |
| WGA | N > 30 | N = 15 | N = 11 |

N is the total number of experiments with 5 – 7 cells each.

**Supplementary Table 8 | Workflows for Glyco-STORM experiments in brain tissue**

| Round | Label in channel 1 | Label in channel 2 | Signal removal (post-imaging) |
|---|---|---|---|
| ***Experiment 1*** *used in Fig. 2a-c, Fig. S6 (nucleus)* | | | |
| 1 | MAA II-AF647 | - | Bleaching |
| 2 | (n.r.) | WGA-CF680 | Bleaching |
| 3 | Con A-AF647 | Fibrillarin::rb-CF680 | Bleaching |
| 4 | UDA-AF647 | Bassoon::ms-CF680 | Bleaching |
| 5 | (n.r.) | VGlut1::gp-CF680 | Elution, Bleaching |
| 6 | (n.r.) | Synaptophysin::rb-CF680 | Bleaching |
| 7 | Hoechst-JF646 | - | - |
| ***Experiment 2*** *used in Fig. 2c, Fig. S6 (nucleus)* | | | |
| 1 | DBA | Fibrillarin::rb-CF680 | Bleaching |
| 2 | Phalloidin-AF647 | WGA-CF680 | Bleaching |
| 3 | UDA-AF647 | SV2::ms-CF680 | Bleaching |
| 4 | Con A-AF647 | (n.r.) | Elution, Bleaching |
| 5 | (n.r.) | Bassoon::ms-CF680 | - |
| ***Experiment 3*** *used in Fig. 2c, Fig. S6 (nucleus); Fig. 2f-j (ER); Fig. 3c, Fig. S7a (Golgi); Fig. S9c (PLC/lysosome)* | | | |
| 1 | VVA-AF647 | - | Bleaching |
| 2 | VEA-AF647 | (n.r.) | Bleaching |
| 3 | PSA-AF647 | GM130::ms-CF680 | Bleaching |
| 4 | CHC17-AF647 | Giantin::gp-CF680 | Elution, Bleaching |
| 5 | PWA-AF647 | TGN38::ms-CF680 | Elution, Bleaching |
| 6 | MAA II-AF647 | (n.r.) | Bleaching |
| 7 | Con A-AF647 | WGA-CF680 | Elution, Bleaching |
| 8 | POL-AF647 | PEX14::rb-CF680 | Elution, Bleaching |
| 9 | PDI::ms-AF647 | - | - |
| ***Experiment 4*** *used in Fig. 2c, Fig. S6 (nucleus)* | | | |
| 1 | VVA-AF647 | Bassoon::ms-CF680 | Bleaching |
| 2 | GM130::ms-AF647 | Golgin97::rb-CF680 | Elution, Bleaching |
| 3 | (n.r.) | WGA-CF680 | Elution, Bleaching |
| 4 | Con A-AF647 | Synaptophysin::rb-CF680 | - |
| ***Experiment 5*** *used in Fig. 2b,c, Fig. S6 (nucleus); Fig. 2 f,g,j (ER); Fig. 3b,c, Fig. S7c (Golgi); Fig. 3d-g, Fig. S9c,d (PLC/lysosome); Fig. 5a, Fig. S13(i) (glycosylation map); Fig. S5 (precision analysis); Fig. S11a,b (perisynaptic structures)* | | | |
| 1 | LCA-AF647 | - | Bleaching |
| 2 | Phalloidin-AF647 | LAMP1::rb-CF680 | Bleaching |
| 3 | PWA-AF647 | LAMP3::gp-CF680 | Bleaching |
| 4 | CHC17-AF647 | WGA-CF680 | Elution, Bleaching |
| 5 | VGlut2::ch-AF647 | Bassoon::ms-CF680 | Elution, Bleaching |
| 6 | LTL-AF647 | EEA1::rb-CF680 | Elution, Bleaching |
| 7 | VAMP7::ms-AF647 | Homer1b/c::rb-CF680 | Elution, Bleaching |
| 8 | POL-AF647 | Rab5::rb-CF680 | Elution, Bleaching |
| 9 | PSA-AF647 | GM130::ms-CF680 | Elution, Bleaching |
| 10 | Con A-AF647 | Rab7::rb-CF680 | - |

| | | | |
|---|---|---|---|
| ***Experiment 6*** | | | |
| *used in Fig. 2b,c, Fig. S6 (nucleus); Fig. 2f,g (ER)* | | | |
| 1 | Con A-AF647 | - | Bleaching |
| 2 | (n.r.) | Bassoon::ms-CF680 | Bleaching |
| 3 | Rab7::rb-AF647 | VGlut1::gp-CF680 | Elution, Bleaching |
| 4 | LAMP1::rb-AF647 | (n.r.) | Elution, Bleaching |
| 5 | PDI::ms-AF647 | WGA-CF680 | - |
| ***Experiment 7*** | | | |
| *used in Fig. 2b,c, Fig. S6 (nucleus); Fig. 2e-g,j (ER); Fig. S9c,e (PLC/lysosome)* | | | |
| 1 | PSA-AF647 | LAMP1::ms-CF680 | Bleaching |
| 2 | LCA-AF647 | LAMP5::gp-CF680 | Bleaching |
| 3 | PWA-AF647 | SCAMP1::rb-CF680 | Bleaching |
| 4 | LTL-AF647 | LAMP2a::gp-CF680 | Elution, Bleaching |
| 5 | Con A-AF647 | EEA1::rb-CF680 | Elution, Bleaching |
| 6 | POL-AF647 | PDI::ms-CF680 | - |
| ***Experiment 8*** | | | |
| *used in Fig. 2b,c, Fig. S6 (nucleus); Fig. 3c, Fig. S7a (Golgi); Fig. S13a(ii) (glycosylation map); Fig. S9c (PLC/lysosome); Fig. S11c,d (perisynaptic structures)* | | | |
| 1 | VEA-AF647 | TGN38::ms-CF680 | Bleaching |
| 2 | VVA-AF647 | WGA-CF680 | Bleaching |
| 3 | PSA-AF647 | Giantin::gp-CF680 | Bleaching |
| 4 | PWA-AF647 | Homer1b/c::rb-CF680 | Elution, Bleaching |
| 5 | SV2::ms-AF647 | (n.r.) | Elution, Bleaching |
| 6 | MAA II-AF647 | GM130::ms-CF680 | Elution, Bleaching |
| 7 | LTL-AF647 | (n.r.) | - |
| ***Experiment 9*** | | | |
| *used in Fig. 2c, Fig. S6 (nucleus); Fig. 3c, Fig. S7a,b (Golgi); Fig. S13a(iii) (glycosylation map); Fig. S9c (PLC/lysosome)* | | | |
| 1 | VVA-AF647 | - | Bleaching |
| 2 | VEA-AF647 | (n.r.) | Bleaching |
| 3 | PSA-AF647 | WGA-CF680 | Bleaching |
| 4 | MAA II-AF647 | Giantin::gp-CF680 | Elution, Bleaching |
| 5 | Golgin97::rb-AF647 | TGN38::ms-CF680 | Elution, Bleaching |
| 6 | CHC17-AF647 | VAMP4::rb-CF680 | Elution, Bleaching |
| 7 | UEA I-DL649 | GM130::ms-CF680 | - |
| ***Experiment 10*** | | | |
| *used in Fig. 2c, Fig. S6 (nucleus); Fig. S9c (PLC/lysosome)* | | | |
| 1 | PSA-AF647 | - | Bleaching |
| 2 | Bassoon::ms-AF647 | (n.r.) | Bleaching |
| 3 | LCA-AF647 | WGA-CF680 | Elution, Bleaching |
| 4 | GM130::ms-AF647 | LAMP1::rb-CF680 | Elution, Bleaching |
| 5 | Synaptophysin::rb-AF647 | GLT1::gp-CF680 | - |
| ***Experiment 11*** | | | |
| *used in Fig. 2c, Fig. S6 (nucleus)* | | | |
| 1 | ABA-AF647 | - | Bleaching |
| 2 | Bassoon::ms-AF647 | Homer1b/c::rb-CF680 | Bleaching |
| 3 | Synaptophysin::rb-AF647 | GLT1::gp-CF680 | Bleaching |
| 4 | S100B::ch-AF647 | WGA-CF680 | Elution, Bleaching |
| 5 | UDA-AF647 | Rab5::rb-CF680 | - |

| | | | |
|---|---|---|---|
| ***Experiment 12*** | | | |
| *used in Fig. 2c, Fig. S6 (nucleus); Fig. S5 (precision analysis)* | | | |
| 1 | - | Bassoon::ms-CF680<br>Piccolo::gp-CF680 (cumulative) | Bleaching |
| 2 | UDA-AF647 | - | Bleaching |
| 3 | (n.r.) | WGA-CF680 | - |
| ***Experiment 13*** | | | |
| *used in Fig. 2b,c, Fig. S6 (nucleus)* | | | |
| 1 | UDA-AF647 | - | Bleaching |
| 2 | MAP2::ch-AF647 | Bassoon::ms-CF680 | Bleaching |
| 3 | - | WGA-CF680 | Bleaching |
| 4 | Phalloidin-AF647 | Rab5::rb-CF680 | - |
| ***Experiment 14*** | | | |
| *used in Fig. 2b,c, Fig. S6 (nucleus); Fig. 2d,f,g (ER); Fig. S5 (precision analysis)* | | | |
| 1 | Con A-AF647 | - | Bleaching |
| 2 | - | Piccolo::gp-CF680 | Bleaching |
| 3 | PDI::ms-AF647 | WGA-CF680 | Bleaching |
| 4 | Synaptophysin::rb-AF647 | - | - |
| ***Experiment 15*** | | | |
| *used in Fig. 2c, Fig. S6 (nucleus)* | | | |
| 1 | UDA-AF647 | WGA-CF680 | - |
| ***Experiment 16*** | | | |
| 1 | - | WGA-CF680 | - |
| ***Experiment 17*** | | | |
| 1 | PWA-AF647 | - | Bleaching |
| 2 | PSA-647 | WGA-CF680 | - |
| ***Experiment 18*** | | | |
| *used in Fig. 2c, Fig. S6 (nucleus)* | | | |
| 1 | DBA-AF647 | - | Bleaching |
| 2 | Synaptophysin::rb-AF647 | PDI::ms-CF680 | Bleaching |
| 3 | MAP2::ch-AF647 | Phalloidin-AF680 | - |
| ***Experiment 19*** | | | |
| *used in Fig. S5 (precision analysis)* | | | |
| 1 | VVA-AF647 | - | Bleaching |
| 2 | Bassoon::ms-AF647 | (n.r.) | Bleaching |
| 3 | VGlut2::ch-AF647 | WGA-CF680 | Bleaching |
| 4 | VEA-AF647 | Giantin::gp-CF680 | Elution, Bleaching |
| 5 | GM130::ms-AF647 | Homer1b/c::rb-CF680 | Elution, Bleaching |
| 6 | (n.r.) | LAMP3::gp-CF680 | Elution, Bleaching |
| 7 | VAMP7::ms-AF647 | TGN38::rb-CF680 | Elution, Bleaching |
| 8 | MAA II-AF647 | Golgin97::rb-CF680 | - |
| ***Experiment 20*** | | | |
| *used in Fig. 4a,c (synapse)* | | | |
| 1 | VGlut1::gp-AF647 | Bassoon::ms-CF680<br>Piccolo::rb-CF680 (cumulative) | Bleaching |
| 2 | VGlut2::ch-AF647 | Homer1b/c::rb-CF680 | Elution, Bleaching |
| 3 | CHC17-AF647 | WGA-CF680 | Elution, Bleaching |
| 4 | Synaptophysin::rb-AF647 | (n.r.) | Elution, Bleaching |
| 5 | VAMP2::rb-AF647 | SV2::ms-CF680 | Elution, Bleaching |
| 6 | VAMP1::rb-AF647 | Rab3a::ms-CF680 | Elution, Bleaching |
| 7 | PSA-AF647 | (n.r.) | - |

| | | | |
|---|---|---|---|
| ***Experiment 21*** *(synapse)* | | | |
| 1 | VGlut1::gp-AF647 | Bassoon::ms-CF680<br>Piccolo::rb-CF680 (cumulative) | Bleaching |
| 2 | PSA-647 | WGA-CF680 | Elution, Bleaching |
| 3 | VGlut2::ch-AF647 | Homer1b/c::rb-CF680 | Elution, Bleaching |
| 4 | CHC17-AF647 | VGlut-CF680 | Elution, Bleaching |
| 5 | Synaptophysin::rb-AF647 | (n.r.) | Elution, Bleaching |
| 6 | VAMP2::rb-AF647 | SV2::ms-CF680 | Elution, Bleaching |
| 7 | VAMP1::rb-AF647 | Rab3a::ms-CF680 | - |
| ***Experiment 22*** *used in Fig. 2b,c, Fig. S6 (nucleus)* | | | |
| 1 | Bassoon::ms-AF647 | Piccolo::rb-CF680 | Elution, Bleaching |
| 2 | (n.r.) | Homer1b/c::rb-CF680 | Elution, Bleaching |
| 3 | Synaptophysin::rb-AF647 | WGA-CF680 | Elution, Bleaching |
| 4 | DBA-AF647 | (n.r.) | - |
| ***Experiment 23*** *used in Fig. S5 (precision analysis)* | | | |
| 1 | Piccolo::rb-AF647 | Bassoon::ms-CF680 | Elution, Bleaching |
| 2 | (n.r.) | Homer1b/c::rb-CF680 | Bleaching |
| 3 | VGlut-AF647 (NB) | VGlut-CF680 (NB) | Elution, Bleaching |
| 4 | Synaptophysin::rb-AF647 | WGA-CF680 | - |
| ***Experiment 24*** *used in Fig. S9a,b (PLC/lysosome)* | | | |
| 1 | PWA-AF647 | Cathepsin D::rb-CF680 | Elution, Bleaching |
| ***Experiment 25*** *(PLC/lysosome)* | | | |
| 1 | PWA-AF647 | Cathepsin D::rb-CF680 | Elution, Bleaching |

In label names, a dash '-' symbolizes direct conjugation and a double colon '::' symbolizes binding by a secondary antibody or nanobody. Abbreviations: AF647/680, Alexa Fluor 647/680; ch, chicken; gp, guinea pig; JF646, Janelia Fluor 646; ms, mouse; NB, nanobody; n.r., not relevant (if label did not work or has no relevance for this study); rb, rabbit.

**Supplementary Table 9 | Workflows for Glyco-STORM experiments in U2OS cells**

| Round | Label in channel 1 | Label in channel 2 | Signal removal (post-imaging) |
|---|---|---|---|
| ***Experiment 26;*** *used in Fig. S10a, Fig. S13b* | | | |
| 1 | VVA-AF647 | Golgin97::rb-CF680 | Bleaching |
| 2 | VEA-AF647 | GM130::ms-CF680 | Bleaching |
| 3 | PSA-AF647 | Giantin::gp-CF680 | Bleaching |
| 4 | CHC17-AF647 | WGA-CF680 | Elution, Bleaching |
| 5 | PWM-AF647 | TGN38::ms-CF680 | Elution, Bleaching |
| 6 | LAMP1::rb-AF647 | PDI::ms-CF680 | Elution, Bleaching |
| 7 | Con A-AF647 | EEA1::rb-CF680 | Elution, Bleaching |
| 8 | MAA II-AF647 | LAMP3::gp-CF680 | Elution, Bleaching |
| 9 | POL-AF647 | Rab7::rb-CF680 | - |
| ***Experiment 27;*** *used in Fig. S8a* | | | |
| 1 | VEA-AF647 | TGN38::ms-CF680 | Bleaching |
| 2 | VVA-AF647 | Golgin97::rb-CF680 | Bleaching |
| 3 | PSA-AF647 | GM130::ms-CF680 | Bleaching |
| 4 | CHC17-AF647 | Giantin::gp-CF680 | Bleaching |
| 5 | PWM-AF647 | WGA-CF680 | - |
| ***Experiment 28*** | | | |
| 1 | PSA-AF647 | Golgin97::rb-CF680 | Bleaching |
| 2 | VVA-AF647 | GM130::ms-CF680 | Bleaching |
| 3 | VEA-AF647 | Giantin::gp-CF680 | Elution, Bleaching |
| 4 | MAA II-AF647 | EEA1::rb-CF680 | Elution, Bleaching |
| 5 | CHC17-AF647 | WGA-CF680 | - |
| ***Experiment 29*** | | | |
| 1 | VEA-AF647 | Giantin::gp-CF680 | Bleaching |
| 2 | PSA-AF647 | GM130::ms-CF680 | - |
| ***Experiment 30*** | | | |
| 1 | VVA-AF647 | Giantin::gp-CF680 | Bleaching |
| 2 | PSA-AF647 | GM130::ms-CF680 | Bleaching |
| 3 | VEA-AF647 | Golgin97::rb-CF680 | Elution, Bleaching |
| 4 | MAA II-AF647 | Syntaxin6::rb-CF680 | - |
| ***Experiment 31*** | | | |
| 1 | DBA-AF647 | - | Bleaching |
| 2 | PNA-AF647 | WGA-CF680 | Bleaching |
| 3 | Lamin a/c::ch-AF647 | (n.r.) | Bleaching |
| 4 | UDA-AF647 | Fibrillarin::rb-CF680 | Bleaching |
| 5 | MAA II-AF647 | Con A-CF680 | Bleaching |
| 6 | Hoechst-JF646 | - | - |
| ***Experiment 32*** | | | |
| 1 | Con A-AF647 | EEA1::rb-CF680 | Bleaching |
| 2 | POL-AF647 | PDI::ms-CF680 | Bleaching |
| 3 | LCA-AF647 | LAMP5::gp-CF680 | Bleaching |
| 4 | PWM-AF647 | WGA-CF680 | Elution, Bleaching |
| 5 | UEA I-AF649 | LAMP1::ms-CF680 | Elution, Bleaching |
| 6 | CHC17-AF647 | Syntaxin6::rb-CF680 | Elution, Bleaching |
| 7 | SCAMP1::rb-AF647 | VAMP7::ms-CF680 | - |

In label names, a dash '-' symbolizes direct conjugation and a double colon '::' symbolizes binding by a secondary antibody or nanobody. Abbreviations: AF647, Alexa Fluor 647; ch, chicken; gp, guinea pig; JF646, Janelia Fluor 646; ms, mouse; NB, nanobody; n.r., not relevant (if label did not work or has no relevance for this study); rb, rabbit.

**Supplementary Table 10 | Workflows for Glyco-STORM experiments in hippocampal neurons**

| Round | Label in channel 1 | Label in channel 2 | Signal removal (post-imaging) |
|---|---|---|---|
| ***Experiment 33*** *used in Fig. S8b, S10b, S13c* | | | |
| 1 | PSA-AF647 | TGN38::ms-CF680 | Bleaching |
| 2 | VVA-AF647 | GM130::ms-CF680 | Bleaching |
| 3 | VEA-AF647 | Giantin::gp-CF680 | - |
| ***Experiment 34*** | | | |
| 1 | PSA-AF647 | Golgin97::rb-CF680 | Bleaching |
| 2 | VVA-AF647 | GM130::ms-CF680 | - |
| ***Experiment 35*** | | | |
| 1 | PSA-AF647 | Bassoon::ms-CF680 | - |
| ***Experiment 36*** | | | |
| 1 | POL-AF647 | Bassoon::ms-CF680 | - |
| ***Experiment 37*** | | | |
| 1 | Bassoon::ms-AF647 | Con A-CF680 | - |
| ***Experiment 38*** | | | |
| 1 | Bassoon::ms-AF647 | WGA-CF680 | - |
| ***Experiment 39*** | | | |
| 1 | Homer1b/c::rb-AF647 | WGA-CF680 | - |
| ***Experiment 40*** | | | |
| 1 | Con A-AF647 | Homer1b/c::rb-CF680 | - |
| 2 | Bassoon::ms-AF647 | - | |
| ***Experiment 41*** *used in Fig.4f,h, Fig. S12* | | | |
| 1 | PSA-AF647 | Homer1b/c::rb-CF680 | - |
| 2 | Bassoon::ms-AF647 | | |
| ***Experiment 42*** | | | |
| 1 | POL-AF647 | Homer1b/c::rb-CF680 | - |
| ***Experiment 43*** | | | |
| 1 | Homer1b/c::rb-AF647 | Con A-CF680 | - |
| ***Experiment 44*** | | | |
| 1 | POL-AF647 | (n.r.) | Elution, Bleaching |
| 2 | Bassoon::ms-AF647 | Homer1b/c::rb-CF680 | Elution, Bleaching |
| 3 | (n.r.) | WGA-CF680 | Elution, Bleaching |
| 4 | POL-AF647 | - | - |
| ***Experiment 45*** *used in Fig.4f* | | | |
| 1 | POL-AF647 | (n.r.) | Elution, Bleaching |
| 2 | Bassoon::ms-AF647 | Homer1b/c::rb-CF680 | Elution, Bleaching |
| 3 | (n.r.) | WGA-CF680 | - |
| ***Experiment 46*** *used in Fig. 4f* | | | |
| 1 | PSA-AF647 | (n.r.) | Elution, Bleaching |
| | Bassoon::ms-AF647 | Homer1b/c::rb-CF680 | - |
| ***Experiment 47*** *used in Fig. 4f,h, Fig. S12* | | | |
| 1 | PSA-AF647 | (n.r.) | Elution, Bleaching |
| 2 | Bassoon::ms-AF647 | Homer1b/c::rb-CF680 | - |
| ***Experiment 48*** *used in Fig. 4f-h, Fig. S12* | | | |
| 1 | Con A-AF647 | Bassoon::ms-CF680 | Bleaching |
| 2 | Homer1b/c::rb-AF647 | WGA-CF680 | - |

| | | | |
|---|---|---|---|
| ***Experiment 49*** <br> *used in Fig. 4f,h-j, Fig. S12* | | | |
| 1 | POL-AF647 | Homer1b/c::rb-CF680 | Bleaching |
| 2 | Bassoon::ms-AF647 | WGA-CF680 | - |
| ***Experiment 50*** <br> *used in Fig. 4f* | | | |
| 1 | POL-AF647 | Bassoon::ms-CF680 | Bleaching |
| 2 | Homer1b/c::rb-AF647 | WGA-CF680 | Elution, Bleaching |
| 3 | (n.r.) | Con A-CF680 | - |
| ***Experiment 51*** <br> *used in Fig. 4f* | | | |
| 1 | POL-AF647 | Bassoon::ms-CF680 | Bleaching |
| 2 | Homer1b/c::rb-AF647 | WGA-CF680 | Elution, Bleaching |
| 3 | (n.r.) | Con A-CF680 | - |
| ***Experiment 52*** <br> *used in Fig.4f,h, Fig. S12* | | | |
| 1 | Con A-AF647 | Bassoon::ms-CF680 | Bleaching |
| 2 | Homer1b/c::rb-AF647 | WGA-CF680 | - |
| ***Experiment 53*** <br> *used in Fig.4f,h, Fig. S12* | | | |
| 1 | PSA-AF647 | (n.r.) | Bleaching |
| 2 | Bassoon::ms-AF647 | Homer1b/c::rb-CF680 | - |
| ***Experiment 54*** <br> *used in Fig. 4f-j, Fig. S12* | | | |
| 1 | Homer1b/c::rb-AF647 | WGA-CF680 | Bleaching |
| 2 | PSA-AF647 | Bassoon::ms-CF680 | - |
| ***Experiment 55*** <br> *used in Fig. 4f,h, Fig. S12* | | | |
| 1 | Homer1b/c::rb-AF647 | WGA-CF680 | Bleaching |
| 2 | POL-AF647 | Bassoon::ms-CF680 | - |

In label names, a dash ‘-‘ symbolizes direct conjugation and a double colon ‘::’ symbolizes binding by a secondary antibody or nanobody. Abbreviations: AF647, Alexa Fluor 647; gp, guinea pig; ms, mouse; NB, nanobody; n.r., not relevant (if label did not work or has no relevance for this study); rb, rabbit.

**Supplementary Table References**


1. Sueyoshi, S., Tsuji, T. & Osawa, T. Purification and characterization of four isolectins of mushroom (agaricus bisporus). *Biol. Chem. Hoppe. Seyler.* **366**, 213–222 (1985).

2. Bojar, D. *et al.* A Useful Guide to Lectin Binding: Machine-Learning Directed Annotation of 57 Unique Lectin Specificities. *ACS Chem. Biol.* **17**, 2993–3012 (2022).

3. Kolberg, J., Michaelsen, T. E. & Sletten, K. Properties of a Lectin Purified from the Seeds of Cicer arietinum * Brought to you by | New York University Bobst Li Authenticated Download Date Brought to you by | New York University Bobst Lib Authenticated. 655–664 (1983).

4. Gautam, A. K., Gupta, N., Narvekar, D. T., Bhadkariya, R. & Bhagyawant, S. S. Characterization of chickpea (Cicer arietinum L.) lectin for biological activity. *Physiol. Mol. Biol. Plants* **24**, 389–397 (2018).

5. RCSB PDB - 3CNA: STRUCTURE OF CONCANAVALIN A AT 2.4 ANGSTROMS PRECISION ANALYSIS. doi:https://doi.org/10.2210/pdb3CNA/pdb

6. THE STRUCTURE OF THE DOLICHOS BIFLORUS SEED LECTIN IN COMPLEX WITH THE FORSSMAN DISACCHARIDE. doi:https://doi.org/10.2210/pdb1LU1/pdb

7. Wu, A. M., Dudek, A. & Chen, Y. L. Recognition factors of Dolichos biflorus agglutinin (DBA) and their accommodation sites. *Glycoconj. J.* **40**, 383–399 (2023).

8. Etzler, M. E. & Kabat, E. A. Purification and characterization of a lectin (plant hemagglutinin) with blood group A specificity from Dolichos biflorus. *Biochemistry* **9**, 869–877 (1970).

9. Damme, E. J. M. Van, Peumans, W. J., Barre, A. & Rougé, P. Plant Lectins: A Composite of Several Distinct Families of Structurally and Evolutionary Related Proteins with Diverse Biological Roles. *CRC. Crit. Rev. Plant Sci.* **17**, 575–692 (1998).

10. RCSB PDB - 2EIG: Lotus tetragonolobus seed lectin (Isoform). doi:https://doi.org/10.2210/pdb2EIG/pdb

11. Kawaguchi, T., Matsumoto, I. & Osawa, T. Studies on hemagglutinins from Maackia amurensis seeds. *J. Biol. Chem.* **249**, 2786–2792 (1974).

12. Yang, Y. *et al.* Characterization, molecular cloning, and in silico analysis of a novel mannose-binding lectin from Polygonatum odoratum (Mill.) with anti-HSV-II and apoptosis-inducing activities. *Phytomedicine* **18**, 748–755 (2011).

13. RCSB PDB - 2LTN: DESIGN, EXPRESSION, AND CRYSTALLIZATION OF RECOMBINANT LECTIN FROM THE GARDEN PEA (PISUM SATIVUM). doi:https://doi.org/10.2210/pdb2LTN/pdb

19. Frost, R. G., Reitherman, R. W., Miller, A. L. & Brien, J. S. O. Crude extracts of. **179**, 170–179

(1975).

20. Fornstedt, N. & Porath, J. Characterization studies on a new lectin found in seeds of Vicia ervilia. *FEBS Lett.* **57**, 187–191 (1975).

21. Goldstein, I. J. & Poretz, R. D. Isolation, Physicochemical Characterization, and Carbohydrate-Binding Specificity of Lectins. in *The Lectins* 33–247 (Elsevier, 1986). doi:10.1016/B978-0-12-449945-4.50007-5

22. Wright, C. S. Crystallographic elucidation of the saccharide binding mode in wheat germ agglutinin and its biological significance. *J. Mol. Biol.* **141**, 267–291 (1980).
