## Supplementary Discussion for "Glyco-STORM: Nanoscale Mapping of the Cellular Glycosylation Landscape"

This Supplementary Discussion contextualizes experimental findings and methodological nuances on lectin validation, characterization (section 1), and performance in Glyco-STORM imaging (section 2). Section 3 primarily focuses on to correlating the precise subcellular binding locations of individual lectins with previous data, integrating present observations into a broader biological context and suggesting directions for future research.

#### 1 Lectin characterization and validation

##### 1.1 Evaluation of lectin specificity

To rigorously evaluate lectin reproducibility, each lectin was investigated in at least five independent wide-field microscopy experiments each encompassing a minimum of five brain tissue samples. For all lectins selected for super-resolution microscopy, at least three confocal microscopy-based experiments were conducted in hippocampal neurons and U2OS cells (Table S7). Lectins that showed variable performance within the same type of sample were excluded from Glyco-STORM experiments (section 1.2). All other lectins demonstrated overall universal staining properties across three biological models, encompassing samples with high (hippocampal and brain stem neurons) and low (U2OS cells) secretory activity. This consistent performance set the foundation for a general recommendation to use these lectins in high-resolution microscopy applications across sample types. Sugar specificities for the different lectins will be discussed in section 4.

To verify whether lectin staining emerged from a particular sugar specificity, we employed elution buffers composed of monomeric or oligomeric carbohydrates that compete with the cellular binding of the respective lectins (Fig. S3a; Table S5). For these experiments, we used brain tissue sections as they provide optimal accessibility of epitopes, therefore not requiring detergent-based permeabilization. We quantified the signal remaining after elution, accounting for the unspecific effects of sugar-free washing and bleaching (Fig. S3b-d). For eight lectins, application of competing sugars resulted in a significant reduction of signal (Fig. S3d, analyzed regions listed in Table S6), with PSA and WGA showing the highest signal loss of ~40% and ~30% with a significance of $p \leq 0.001$ when eluted with α-methylmannoside and N-acetylglucosamine (GlcNAc), respectively (Fig. S3d,e; Supplementary Discussion, section 1.1). However, not all lectins showed significant signal reduction upon sugar-based elution. This outcome was anticipated, due to the overall lower affinity of most lectins to free monomeric or oligomeric carbohydrates as compared to *in situ* glycosylation sites where binding could be strengthened by neighboring sugar moieties within the more complex polysaccharides and protein parts to which the carbohydrate chains are attached (Van Damme, 2011). For example, although α-methylmannoside is considered as one of the most efficient eluting sugars for Con A, its affinity to Con A is considerably lower compared to a trimannoside (Mandal and Brewer, 1993). The specificity of most lectins used in this study was previously demonstrated by elution experiments (Goldstein et al., 1997), enzymatic assays (Sabbieti et al., 2000), reproducible results from histochemistry (Manning et al., 2017), affinity chromatographies (Itakura et al., 2017), microarrays (Wang et al., 2014), and machine learning-based analysis of microarrays (Bojar et al., 2022).

To monitor and identify possible cross-reactions between lectins and antibodies, experiments were strategically designed such that the order of label application and the label pairs co-stained in one round varied among biological replicates (Table S8). For each lectin, specific experiments were conducted where it was applied alone in the initial round of labeling, ensuring the complete avoidance of cross-talk and potential cross-reactions with other lectins or antibodies. This approach allowed for the exclusion of any lectins or antibodies that demonstrated inconsistent performance as compared to the single-label reference, ensuring that only reliable candidates were used in further analyses.

Our results, together with published data, provide strong evidence supporting the specificity of the lectin signals obtained in Glyco-STORM experiments.

##### 1.2 Lectins with variable performance

Confocal microscopy identified variability in the performance of some lectins (Fig. S2). HHA and CAL showed a low fluorescence intensity, while UDA exhibited variations in signal strength. PNA and ABA exhibited inconsistent staining patterns across individual experiments. PNA only occasionally showed pronounced nuclear staining. ABA displayed peri-calyceal signals that partially colocalized with WGA (data available upon request). Due to low signal or heterogeneity in signal and staining, HHA, CAL, PNA, and ABA were not further used in Glyco-STORM experiments. UEA I was excluded because the commercially available conjugate was labeled with the fluorophore DyLight-649, which was incompatible with our imaging and analysis pipelines. A more detailed discussion on the performance of these lectins deduced from confocal microscopy experiments and possible explanations for the observed heterogeneity are summarized in section 4.

#### 2 Lectin performance and Glyco-STORM technique

In the following, key aspects leveraging lectins as optical probes for super-resolution microscopy are listed:

- ***In situ* glycosylation mapping**
  - Nano-scale in situ glycome mapping for direct correlation with cellular function
  - Analysis of glycosylation dysfunction in samples from deficient organisms/cells
- **Revelation of new structural features**
  - Distribution of glycan-shielded molecules at the nano-scale
  - Exploration of suborganellar architecture
- **Small probe size: 3-7 nm**
  - Minimized linkage error
- **Direct fluorophore labeling**
  - Streamlined application: single-step, reduced time
  - Combinability for multiplex experiments
- **High binding diversity**
  - Large selection of lectins available, among these >100 characterized for their targeted glycosylation motifs

- **Sugar binding specificity**
  - Enabling glyco-centric labeling
  - Broad labeling of abundant and/or diverse glycans can provide environmental context
- **Diverse affinities suitable for various techniques**
  - High-affinity lectins: for methods like STORM, SIM, STED, MINFLUX
  - Low-affinity lectins: for methods like PAINT and as weak-affinity labels for other microscopy methods such as STED, confocal and single-particle tracking (Albertazzi and Heilemann, 2023)
- **Cost-effectiveness**

#### 3 Limitations of this study

The most relevant limitations are discussed in the main manuscript, here we want to point out additional considerations.

First, lectins bind to several carbohydrate motifs, and to each of these with different affinities, some of which can be very low. These weak interactions may explain the minor diffuse signals observed for certain lectins.

Second, biological variability of lectin preparations can affect imaging results. Most lectins used in this study are plant-purified, and their exact protein structure and expressed isoforms can vary based on breeding effects or harvesting times. Such variations may alter binding properties and overall specificity, introducing heterogeneities in lectin performance. In consequence, lectin performance may vary by vendor and lot number.

Third, if multiplexed super-resolution imaging following the maS³TORM concept is used, the detergent-based chemical elution used in some rounds to remove antibodies could affect membrane integrity and thereby influence glycan abundance and/or accessibility of membrane-associated glycans. To minimize bias from variable cell integrity, we alternated the order of labeling across experiments and carefully monitored lectin performance before and after elution, observing no significant differences.

Fourth, since commercially available lectins are typically fluorophore-labeled by amine-reactive crosslinker chemistry, the degree of labeling depends on the size of the lectin and the number of amino-reactive residues. This limits direct quantitative comparisons between different lectins. Interestingly, compared to antibody markers, the fluorescence signal recorded for most lectins was lower (e.g., see Fig. 2f and Fig. S12). Multiple reasons can explain this: (1) Higher protein abundance in the respective regions as compared to the lectin-binding motifs, (2) higher degree of labeling of secondary antibodies as compared to lectins, (3) signal enrichment due to multiple epitopes in the case of polyclonal primary and/or secondary antibodies, (4) a generally lower binding affinity of lectins that might be even lower upon fluorophore conjugation.

Lastly, the selection of model samples needs consideration. We chose three different cell/tissue models, with two of our models – hippocampal neurons and brain sections – of neuronal origin. Although the presynaptic calyx of Held is glutamatergic, it is unusual due to its large size, hundreds of active zones, and fast-firing properties. The postsynaptic principal cell on the other hand features inhibitory GABAergic projections. Given the diversity among neurons, studies on other neuronal cells would help to expand our understanding of universal principles of neuronal glycosylation. U2OS cells are an immortal cancer cell line and may not fully represent native glycosylation patterns due to cancer-related alterations. Expanding the use of lectins to other cell models will further enrich the characterization of glycan binding and provide a broader perspective.

### 4 Cellular glycosylation patterns and biological implications revealed by Glyco-STORM

#### 4.1 Nucleus- and cytoplasm-associated glycosylation

The majority of tested lectins displayed strong to sporadic signals in both the nucleoplasm and the cytoplasm, suggesting a broad sugar repertoire in these areas. In this section, the nucleoplasmic and cytoplasmic staining patterns of these lectins are discussed with respect to the specific sugar binding properties.

*Variable performance of lectins binding nucleocytoplasmic targets*

Many lectins that showed signal in the nucleus or cytoplasm showed variable staining performance. Possible reasons include factors previously discussed in the main text – such as their binding to O-glycosylated soluble proteins prone to washout, the instability of their target glycans, or reduced lectin affinity. Another notable factor for the volatile signal could be the different state of the cultured cells or cells within the tissue. For example, it has been demonstrated in an osteoblast cell line that PNA preferentially recognizes nuclei of mitotically active cells (Sabbieti et al., 2000). Among lectins whose nuclear signals were analyzed using super-resolution data, UDA and, to a lower extent, DBA showed variations in fluorescence intensity. Four lectins – HHA, CAL, PNA, and ABA – that were excluded from all Glyco-STORM analyses due to their variable performance, showed dispersed signals in the cytoplasm and the nucleus.

*Gal-and GalNAc-binding lectins*

We observed that many lectins with high nuclear signal were specific (not exclusively) for galactose- (Gal) or N-acetylgalactosamine- (GalNAc) containing motifs (Table S1). These included DBA, UDA, MAA II, WGA, VVA, and PWA, whose staining was assessed by localization density (Fig. S6) and cluster analysis in Glyco-STORM images (Fig. 2c). Interestingly, also lectins with low intensity or inconsistent performance (CAL, UDA, PNA, and ABA), showed strong signal in the nucleus in some experiments (Fig. S2). All of the above listed lectins also showed cytoplasmic staining (Fig. 1d-f and Fig. S1, S2). Increasing evidence indicates that the nucleus contains a substantial number of proteins that are modified by O-linked α-GalNAc (M. West et al., 2022), such as lamins (Cejas et al., 2020, 2019) or the tumor suppressor p53 (Xu et al., 2020). Besides the nucleus, many of the O-GalNAcylated proteins are also found in the cytoplasm that can explain the concurrent signal of the same lectins in both compartments.

DBA is a lectin that was found to specifically target the Forssman antigen and α- and β-linked GalNAc sugar moieties (Bojar et al., 2022). In this study, DBA displayed strong nucleocytoplasmic signals (Fig. S1i) and exhibited the highest localization density in the nucleus (Fig. S6). An earlier study uncovered DBA binding sites in the cytoplasm of osteoblast cells that are presumably attributable to α-linked GalNAc as β-Galactosidase digestion did not influence DBA affinity (Sabbieti et al., 2000). This study also showed that permeabilizing cells before fixation reduced cytosolic DBA staining in favor of nuclear signal. Likewise, WGA, besides other specificities (discussed in upcoming sub-sections), may interact with nuclear and cytoplasmic glycans due to its GalNAc-binding properties.

MAA II, specific for α2-3 sialylated Galβ1-3GalNAc, showed signals throughout the cytoplasm and nucleoplasm (Fig. S1e; Fig. 2a,b) and displayed the highest density of clusters (radius >28 nm) in the nucleus (Fig. 2c). Interestingly, the clustered signal in the nucleolus showed nano-scale correlation with the nucleolus marker fibrillarin (Fig. $2a_{ii}$, Fig. S1e, second panel). This is in agreement with a recent study reporting the presence of SA-capped Galβ1,3-GalNAc O-glycans in the nuclei of HeLa cells (Cejas et al., 2020). More specifically, the authors identified histone and ribosomal proteins as being constitutively modified by Galβ1,3-GalNAc (Cejas et al., 2020, 2019). This suggests that nucleolar MAA II signals may correspond to ribosomal proteins linked to the ribosomal RNA (rRNA). Hence, clustered nuclear MAA II signal may help to explore the precise localization of nuclear glycans and their role in ribosomal assembly and chromatin organization.

VVA preferentially binds GalNAcβ1-4GlcNAc and terminal α/β-GalNAc (Bojar et al., 2022). Our data showed sparsely distributed clustered signal in the nucleus (Fig. 2b,c) and the cytosol (Fig.

S3b). In line with our observations, Cejas et al. demonstrated that VVA targets α-GalNAc O-glycans in the nuclei of HeLa cells (Cejas et al., 2019), where endogenous O-GalNAc signals appeared as confined clusters detectable by confocal imaging. These findings align with the sparse signals we observed using Glyco-STORM (Fig. 2b,c). Some proteins modified by O-linked α-GalNAc, such as p53, are known to localize in both the nucleus and cytoplasm, supporting our observations.

Nucleus-specific staining of O-GalNAc by PNA was previously demonstrated in human cells (Cejas et al., 2020). Previous research in Py1a osteoblasts indicated that PNA binds to nuclear glycans specifically in mitotic cells (Sabbieti et al., 2000). Collectively, this implies a dynamic shift in the presence or accessibility of PNA-reactive glycan structures across the cell cycle, pointing towards a role of O-GalNAcylation in cell division regulation. This may also account for the inconsistent staining of PNA and other O-GalNAc-binding lectins across biological replicates (Fig. S2d). While there is substantial evidence that O-G**lc**NAcylation regulates the cell cycle, the role of O-G**a**lNAcylation remains to be further investigated.

Several lectins that displayed nucleocytoplasmic signal (such as VVA, PWA**,** UDA, and ABA) have a specificity for N-acetyllactosamine (LacNAc) or other Gal-GlcNAc-containing motifs. However, it is unclear whether this sugar motif contributes to their binding, given that LacNAc attachment typically occurs in the Golgi apparatus and resulting modified glycans are usually transported to the plasma membrane. Although such glycans are generally not expected in the nucleus, some proteins with these glycosylation motifs, such as NCAM, have been reported within the nuclear compartment (Westphal et al., 2017, 2016).

In summary, the observation that many lectins that are specific for O-linked Gal, GalNAc, or Gal-GalNAc stain the nucleus and cytoplasm is consistent with the increasing evidence that O-glycosylation, specifically involving these sugar residues, occurs in these compartments. This supports the role of O-glycosylation in intracellular processes, including nuclear and cytoplasmic signaling.

*Sialic acid-binding lectins*

Two of the Gal- and GalNAc-recognizing lectins that showed nuclear signal – MAA II and WGA – also specifically target sialic acid (SA). While the nucleoplasm is the location of free sialic acid (Neu5Ac) where it is converted to cytidine monophosphate-sialic acid (CMP-SA), it is unlikely that these lectins are binding to free Neu5Ac or CMP-SA as they were reported to primarily recognize

SA residues that are α-linked to other sugars (specifically, α2-3-linked to Galβ1-3GalNAc2 in case of MAA II).

MAA II preferentially binds to SA in specific carbohydrate structures on O-glycans. Although conclusive evidence for sialylated O-glycans in the nucleus remains limited, several studies provide supportive indications. For example, the lectin SNA preferentially binding to α2-6 sialic acid linked to LacNAc was demonstrated to bind nuclear pore proteins p62 and nup153 in neuroblastoma cells (Emig et al., 1995). Although the nucleoplasmic signal was not explicitly discussed, a close-up look onto the Western blot revealed a signal around 35 kDa that was comparable to the plasma membrane fraction detected by SNA. Another study that was focused on mammary morphogenesis, demonstrated high levels of α2-6-linked SA (determined by SNA staining) in a malignant human breast epithelial cell line (Bhat et al., 2016), whereas non-malignant cells also exhibited signal for α2-6-linked SA, but at lower levels. Recent studies suggest that some sialylated glycoproteins, such as fragments of the neuronal adhesion molecule (NCAM) can get endocytosed, transported to the cytoplasm, and internalized into the nucleus and act as transcription factors in neurons and CHO cells (Westphal et al., 2017, 2016). However, given its specificity for diverse sugar motifs, WGA could also bind to other glycoconjugates in the nucleus, e.g., through its above-mentioned GalNAc recognition capabilities (see sub-section *"Gal- and GalNAc-binding lectins"*). Further explorations of the binding targets of MAA II and WGA combined with super-resolution techniques could reveal novel insights into nuclear functions and dynamics of glycans in neuronal and non-neuronal cells.

*GlcNAc-binding lectins*

O-GlcNAcylation is one of the most prevalent forms of glycosylation in the cytoplasm and nucleoplasm, where it modifies proteins involved in critical cellular processes such as transcription, signal transduction, and cytoskeletal organization. This dynamic and reversible modification, regulated by the enzymes O-GlcNAc transferase (OGT) and O-GlcNAcase (OGA), plays a key role in controlling cellular responses to stress and regulating gene expression. Multiple lectins with Gal/GalNAc specificity also bind GlcNAc-containing motifs. For the lectins PWA, VVA, UDA, and ABA, GlcNAc could be recognized as part of the LacNAc binding motif (Galβ1-4GlcNAc) that could be attached to the core O-GalNAc (described in the "Gal-and GalNAc-binding lectins" subsection). ABA also has a high preference for β-GlcNAc, and β-linked GlcNAc is typically found in case of O-GlcNAcylation, which is a well-known and crucial process within the nucleus as well as in the cytoplasm (Bond and Hanover, 2015; Dupas et al., 2023). Finally, UDA and PWA also have chitin specificity that, however, is not expected to be present in the nucleoplasm or cytoplasm of mammalian cells.

Specificity of WGA to the O-linked β-GlcNAc modification of nuclear pore complex proteins has been well-documented (Li and Kohler, 2014) and its binding at the nuclear pore was previously reported using electron microscopy (Panté and Aebi, 1996) and super-resolution microscopy (Löschberger et al., 2012). We confirmed WGA's clustered perinuclear staining, revealing an interval corresponding matching the expected distance between nuclear pores in all three model systems: brain tissue (Fig. 2a,d), hippocampal neurons (Fig. S1f) and U2OS cells (Fig. 1f). In U2OS cells, we previously validated WGA's nuclear pore localization via Nup133 co-staining (Klevanski et al., 2020). Interestingly, besides staining nuclear pores, WGA also showed a clustered signal throughout the nucleoplasm (Fig. 2a-c) and the cytoplasm. In the nucleus, WGA could target histone components or other chromatin-associated proteins that have been shown to be modified by O-GlcNAcylation in the nucleus or the cytoplasm with subsequent import to the nucleus (Dupas et al., 2023; Hart et al., 2007). Given the robustness of the nuclear WGA signal, this suggests a use of WGA in combination with labels for chromosome-binding molecules to investigate the nucleoplasmic glycome and its relation to chromatin structure.

*Fucose-binding lectins*

Several lectins that exhibited strong nuclear and cytoplasmic staining possess fucose specificity. This includes LTL, which binds α1-3-linked Fucose (Fuc), and UEA I, which binds α1-2-linked Fuc. LCA and PSA are specific to core-fucosylation (Fucα1-6) and showed weaker and sparser staining (Fig. 2b,c; Fig. S6). Previous studies have reported fucosylation of nuclear proteins (Lipińska et al., 1994) and also LTL and UEA I signal in the cytoplasm and nucleus of some in Py1a osteoblast cells (Sabbieti et al., 2000). The nucleoplasmic localization of LCA has previously been attributed to its binding to chromatin in sea urchin cells (Hart et al., 1989). More recently, nucleolin, a protein involved in chromatin condensation, was shown to be fucosylated in cultured bovine endothelial cells and malignant human A431 cells (Aldi et al., 2009). Limited evidence from experiments using bovine serum albumin (BSA) conjugated to monosaccharides suggests that both fucosylation and mannosylation could act as nuclear localization signals (Duverger et al., 1993).

*Mannose-binding lectins*

Mannose-specific lectins also exhibited nucleoplasmic and cytoplasmic staining, supporting the notion that both fucosylation and mannosylation may function as nuclear localization signals.

Among the lectins with significant nuclear signal (Fig. S6), Con A and UDA preferentially bind terminal mannose (Man). Con A that preferentially binds glycans containing mannose-oligosaccharides with three to nine terminal mannose moieties ($Man_3$ to $Man_9$) showed sparse clustered signal in the nucleus (Fig. 2a-c). Previous research demonstrated that Con A can bind inside the nucleus (Hart et al., 1989) and proteins in nuclear fractions of hamster liver and hepatoma (Lipińska et al., 1994). However, the nuclear binding of Con A has been controversially discussed in the literature, with some studies failing to confirm these findings (Sabbieti et al., 2000). Other mannose-binding lectins, such as POL that preferentially binds $Man_3$ and VEA with α-Man specificity, showed lowest nuclear signal. In summary, given that the presence of mannosylated proteins in the nucleus is not supported by the current body of literature, whether the nuclear signal of Con A and UDA is due to mannose binding remains an open question.

In conclusion, Glyco-STORM is an effective method to spatially map the glycosylation of nucleus-organizing molecules *in situ*. This approach not only confirmed the presence of previously proposed glycans but also revealed the binding of lectins with specificities not typically associated with the nucleus, challenging conventional understanding of glycan distribution and glycosylation processes. This approach enables direct correlation of the glycosylation states of nuclear molecules with their specific sub-nuclear locations, facilitating the development of comprehensive spatio-functional models and uncovering potential roles of glycosylation in key nuclear processes.

##### 4.2 Endoplasmic reticulum glycosylation

Glyco-STORM imaging experiments showed that Con A and POL signals are enriched in the endoplasmic reticulum (ER), as confirmed by co-staining with a monoclonal antibody against the protein disulfide isomerase (PDI/PDIA1). Evaluation of the nanoscale spatial distribution of Con A, POL, and PDI revealed a partial anti-colocalization between the lectins and PDI (Fig. 2d,e,h,i). Several explanations can account for this finding. First, although PDI is reported to bind both non-glycosylated and glycosylated proteins (Rutkevich et al., 2010), its exact substrate preference is not known. In contrast, another PDI family member – ERp57/PDIA3 – preferentially binds glycosylated proteins (Maattanen et al., 2006). This specificity is conferred by ERp57 associating with the intrinsic lectins calnexin or calreticulin while PDI lacks the domain required for the interaction with these lectins (Zapun et al., 1998). Interestingly, while calnexin or calreticulin binding to ERp57 increased its activity, it led to the reduction of PDI binding to its substrate. A possible interpretation is that residual ER regions lacking PDI may be occupied by the presumably more abundant ERp57 (according to The Human Protein Atlas ("The Human Protein Atlas," n.d.)), and could hence explain a lower lectin signal in the PDI-occupied regions. Another possible reason for the anti-correlated signal of lectins and PDI staining is that the antibody used in this study (Table S3) is directed against the C-terminus of PDI, located near its active site. Therefore, it cannot be excluded that the antibody epitope at enzymatically active PDI molecules is masked by their protein substrates, potentially leading to the antibody detecting only the inactive PDI pool. Finally, it is conceivable that other glycan-binding enzymes, such as mannose-trimming enzymes, or lectins involved in ER-associated degradation, could mask Con A and POL binding sites of glycoproteins in proximity to PDI (Zhang et al., 2021). The partial anti-colocalization of lectins and the PDI marker is presumably not due to molecular crowding caused by antibodies or lectins, as these labels were used in different staining rounds. In most experiments (e.g., Experiment 3, Table S8), PDI antibodies were applied after multiple rounds of chemical elution and photo-bleaching, presumably causing photo-unbinding (demonstrated for some antibodies (Heinze et al., 2009; Klevanski et al., 2020)). Also the two ER-binding lectins Con A versus POL showed a low degree of colocalization. A possible explanation could be different binding propensities to different sugar motifs by Con A as compared to POL.

A recent report that used dual-color *d*STORM demonstrated that the ER features two distinct tubule forms in COS-7 cells, U2OS cells, and cultured astrocytes (Wang et al., 2022). The two ER tubule forms comprised an ultrathin and a thick ribbon-shaped structure. Proteins with large intraluminal domains were found excluded from the ultrathin tubules. For example, two interactors of the ER isomerase ERp57 – calnexin and calreticulin – that have large intraluminal domains were found in thicker tubules interrupted by ultrathin ER stretches, resulting in a fragmented signal similar to the PDI signal that we observed in the current study. The fragmented PDI signal is also in line with previous observations in human umbilical vein endothelial cells (Lippok et al., 2016). Given that PDI is a 57 kDA protein localized in the ER lumen, it would be expected in the thick ER compartments. Indeed, the full width at half maximum (FWHM) of 77.5 nm that we measured for PDI is markedly higher than ~50 nm FWHM measured by Wang et al. for the ultrathin ER tubule stretches using primary and secondary antibodies against the outer epitope of the Rtn5 protein. As PDI is a luminal ER protein, we expect that the linkage error is spatially restricted by the ER tubule membranes. Thus, with the lipid bilayer measuring approximately 4 nm and assuming that the antibodies remain confined within the tubule interior, the PDI signal could approximately indicate that the native size of the thick tubule stretches is about 85.5 nm.

Furthermore, we observed that Con A and POL signals in the ER showed a FWHM of 184.5 nm and 143.9 nm in ER stretches that were encompassed by the 400 nm wide transversal selection centered around the PDI clusters (only PDI-containing stretches were transversally measured in the current study). Even considering the largest reported ER tubule thickness of 100 nm, we would not expect such high FWHM values if the lectin staining would be restricted solely to the lumen of the ER tubules. Conversely, if the lectin staining would be confined entirely to the outer membrane, we would anticipate a bimodal distribution that we did not observe. Thus, we hypothesize that Con A and POL could bind both in the ER tubule lumen and the exterior where lectins could bind to dolichol-linked oligo-mannose. However, considering Con A's maximal size of 7 nm, and POL's likely smaller dimensions (POL ~48 kDa vs. Con A ~104 kDs; POL lacks a known 3D structure), we would not expect a FWHM surpassing 120 nm. The considerably higher FWHM values from POL and Con A suggest that they could also bind carbohydrates around the tubules. While this assumption requires further investigation, our hypothesis that lectins could recognize both oligo-mannose at the outer ER membrane and high-mannose glycans inside the ER lumen is supported by earlier electron microscopy studies where Con A has been shown to bind both the interior and the exterior of ER tubules (Guillouzo and Feldmann, 1977; Pinto Da Silva et al., 1981). More experiments are necessary to prove this possibility and explore the spatial organization of peri-ER carbohydrates.

##### 4.3 Glycosylation at the Golgi apparatus and post-Golgi regions.

*Axial Golgi organization*

The axial stratification of the Golgi stacks and the peak distances as visualized by antibodies and lectins (Fig. 3a-c) was in line with previously quantified super-resolution data based on antibody staining (Tie et al., 2018; van Bommel et al., 2023). For example, we measured a GM130-to-TGN38 distance of 230 nm for the primary TGN38 peak, which is in good agreement with previous data obtained from cultured mouse neurons with a GM130-to-TGN38 distance of 272 nm (van Bommel et al., 2023). However, in our line profile analysis, TGN38 also displayed a secondary peak at 370 nm. While the TGN38 signal did not show pronounced double-lines in individual super-resolution images, we observed that some Golgi regions were more densely packed in the *cis*—*trans* direction than others (red and green double-headed arrows in Fig. S7$a_{i,iii}$ denote the higher and lower distance between GM130 and TGN38, respectively). Thus, the secondary peak could correspond to Golgi stretches with larger spaces between the GM130 and the TGN38 signal, a consequence of averaging steps in image analysis independent of Golgi packaging.

Interestingly, Giantin was not only found at the *cis*-Golgi region where it is known to be delivered by COPI vesicles (Sönnichsen et al., 1998), but also coincided with TGN38. As the selection of areas for quantification mainly relied on the GM130 signal and did not account for the localization of Giantin, our analysis may include line profiles laid through the Golgi rimming regions or non-compact Golgi regions whose exact organization remains not fully clarified. Given that COPI vesicles are involved in bidirectional transport within the Golgi stacks, they could carry Giantin throughout the Golgi, including the *trans*-Golgi level, providing a plausible explanation for the overlap between Giantin and TGN38 (Popoff et al., 2011). This is supported by a previous electron microscopy study reporting Giantin's presence across all Golgi cisternae, from *cis* to *trans* (Koreishi et al., 2013).

While antibody stainings were more restricted to individual cisternae, glycan staining revealed broader and more heterogeneous patterns. All lectin signals showed clear upstrokes on the *cis*-directed side, reflecting the occurrence of their main binding motif along the stack. After peaking, signal intensity for most lectins remained elevated, which is expected since once modified, the glycans persist and continue their transport through the *trans*-Golgi network (TGN) to their final destinations. However, for some lectins, such as for VVA, the baseline signal dropped more distinctly in the *trans*-Golgi through TGN area and remained low. This can be attributed to the preference of VVA to unbranched terminal β-GalNAc, in contrast to more complex glycans modified by fucosylation or sialylation, which occur from the medial Golgi through the TGN and could inhibit VVA binding (Bojar et al., 2022; Puri et al., 1992).

Early studies reported that PSA is a mannose-binding lectin (Trowbridge, 1974), while recent machine-learning based analyses reveal core fucose as the binding determinant (Bojar et al., 2022). Our morphometric analysis aligns with the latter finding, as the trailing edge for PSA was found around the medial Golgi cisternae, where core fucosylation of N-linked glycans occurs. The PSA signal continued throughout the *trans*-Golgi and the TGN. In agreement with this preference for fucosylated glycans, we found almost no PSA staining at the ER, while Con A and POL were enriched in the ER. Binding of PSA to the Golgi area was previously proposed, amongst others based on confocal imaging of lectin histochemistry in HeLa cells (Morgan et al., 2013). An early electron microscopy study confirms our results and demonstrates that PSA signal predominated in penultimate cis-cisternae and transport vesicles and at the *cis*- and the *trans*-Golgi side of enterocytes (Pavelka and Ellinger, 1989).

WGA is long known to bind in the Golgi region in various cells (Hiller and Weber, 1982). Using electron microscopy, WGA binding has been precisely localized to the *trans*-Golgi, TGN, and lysosomes (Tartakoff and Vassalli, 1983). A chemical perturbation study leading to dispersal of post-Golgi vesicles demonstrated that WGA mainly binds to TGN-derived vesicles (Kanazawa et al., 2008).

While TGN-proximal clathrin-coated vesicles rich in WGA showed decreased PSA signal, distal clathrin-rich areas featured higher PSA and lower WGA signal. This indicates that TGN-derived vesicles are more likely to contain a higher proportion of newly β-GlcNAc-terminated or sialylated, presumably more mature, glycans. Lower WGA-reactive contents of distal clathrin-rich vesicles, on the other hand, could be explained by processes that can occur at the plasma membrane, such as glycan trimming or modifications that would reduce their accessibility to WGA and could represent glycans that are destined for degradation. Furthermore, whether fucosylated glycans are less abundant in clathrin-mediated transport routes from and to the TGN remains to be studied. Although MAA II also recognizes sialylated glycans, its distribution deferred from that of WGA and clathrin-rich regions. This could indicate that sialyl-Thomsen-nouveau antigen (sialyl-Tn; O-linked SAα2-6GalNAcα) carrying glycans recognized by MAA II are scarce in clathrin-coated vesicles and that the WGA signal associated with clathrin vesicles is mainly attributable to WGA's preference for GlcNAc-terminated glycans.

Moderate VEA signal (plateau around 0 nm in Fig. 3c) coincided with Giantin, suggesting that respective mannose-and glucose-containing motifs are present in COPI vesicles. The main signal peaks were found in the medial Golgi, *trans*-Golgi and TGN areas. While the signal dropped in clathrin-positive areas, it maintained an elevated baseline level and showed an ascending tendency in post-clathrin regions (from 1100 nm onwards; Fig. 3c). It is of note that although VEA shares similar sugar specificity with Con A, it did not show signal in the ER, suggesting that specific post-ER modifications influence its binding specificity. Interestingly, we find that VEA peaked in *cis*-most TGN regions while WGA signal was increased in more distal TGN regions. In contrast, clathrin-positive post-TGN areas with increasing WGA signal showed reduced VEA binding.

*Lateral Golgi organization*

We also studied the molecular content along the lateral axis parallel to the cisternae. We found alternating signals of Giantin and GM130, with the Giantin signal situated at the rim of presumed Golgi ribbons and in-between GM130-positive stretches likely marking the core of individual Golgi stacks (Fig. 3a,h; Fig. S7). While previous super-resolution microscopy studies in cells treated with drugs facilitating Golgi dispersal demonstrated Giantin's location at the rim of Golgi mini-stacks (Schueder et al., 2024; Tie et al., 2018), to our knowledge, the inter-stack location of Giantin within Golgi ribbons of intact cells has only been reported in one study using electron microscopy (Koreishi et al., 2013). This may be due to the high three-dimensional complexity of the Golgi apparatus. The thin (400 nm) brain tissue sections and 3D super-resolution imaging approach employed in this study, along with the ultra-thin sections used for electron microscopy (Koreishi et al., 2013), enable detailed delineation of this complex organelle.

In some experiments, we observed that the TGN38 signal was correlated with the Giantin pattern (Fig. S7$a_{i,iii}$, line profiles). Interestingly, in Giantin-overarched areas, TGN38 showed higher signal intensities and was found in closer proximity to the Giantin marker as compared to neighboring GM130-overarched and GM130-rich regions. Some lectins, including VEA and WGA, were more closely associated with Giantin-covered regions as compared to regions lacking Giantin. Based on previous studies demonstrating that Giantin could be (1) localized in the non-compact regions in-between adjacent stacks and (2) involved in organizing these regions (Satoh et al., 2019), we hypothesize that Giantin-rich stretches correspond to non-compact Golgi regions. These regions are characterized by a complex structure with fenestrated cisternae traversed by transport vesicles and microtubules and have been proposed to be areas of alternative transport through the Golgi, bypassing the tightly stacked region (Saraste and Prydz, 2019). Thus, lectin signal found in these areas could be associated with anterogradely or retrogradely transported glycans that are not processed via the classical cisternal Golgi 'glycan assembly line'. On the basis of this model, VEA signal could indicate high-mannose glycans that are not further modified by the intracisternal glycosylation machinery and are directly passed to the TGN for transport to their final destination. In case of WGA, the signal found in *cis*-face proximity could reflect retrogradely transported glycans (Dupas et al., 2023).

In summary, Glyco-STORM provided high-resolution spatial information of the organization of glycan assembly within the Golgi apparatus. Further studies, e.g., using electron microscopy or super-resolution microscopy combined with expansion microscopy (Bond et al., 2022), could further refine the nano-architecture of the Golgi glycan map.

##### 4.4 Glycosylation of vesicular organelles

*Endosome and lysosome markers*

To determine the identity of spherical organelles positive for PWA, we tested several antibody makers, including those for lysosomes (e.g., LAMP1, LAMP2A, cathepsin D), early endosomes (e.g., Rab5 and EEA1), late endosomes (e.g., Rab7, LAMP5), and clathrin (CHC17). While most antibody markers exhibited sparse signals in brainstem neurons (Fig. 3d, third column), precluding unequivocal organelle identification, cathepsin D showed high abundance and correlation with PWA-positive structures (Fig. S9a,b). As a protease abundant in pre-lysosomal compartments (PLCs), such as endosomes, and in lysosomes (Turk et al., 2012), cathepsin D effectively identifies these organelles as targets of PWA. While LAMP markers displayed particularly limited signal, Rab5 and EEA1 (in brain tissue: Fig. 3d; in U2OS cells: Fig. S10a), along with clathrin-coated bulges (Fig. $3g_{i,ii}$) were more frequently observed at the periphery of these organelles. These observations suggest that PWA-positive organelles are primarily early or late endosomes. Since LAMPs are increasingly considered insufficient as sole lysosome markers (Cheng et al., 2018), further confirmation is needed to estimate the fraction of PWA-positive organelles with lysosomal identity.

*Lectin-binding organelles of the endolysosomal system*

Glyco-STORM analysis revealed that compartments of the endolysosomal system featured a highly diverse glycosylation. The variability in lectin staining patterns and intensities across different experiments and at distinct PLCs/lysosomes further contributes to this complexity. At the same time, the results obtained with Glyco-STORM provide valuable insights into these organelles (Fig. 3d-g; Fig. S9b-e), as detailed in the following discussion paragraphs.

The high selectivity of PWA for PLCs/lysosomes, along with the absence of a pronounced PWA signal at the plasma membrane, suggest that its binding motifs likely arise during glycan degradation. This assumption aligns with a previous study demonstrating that removal of terminal fucose or sialic acid unmasks poly-LacNAc motifs targeted by PWA (Heiskanen et al., 2009). In addition, this is consistent with results obtained in extracellular vesicle uptake assays where lectin staining was used to assess the glycan specificity before and after treatment with different glycosyl transferases. The removal of sialic acid by neuraminidase treatment led to an increased PWA signal explained by the exposure of the underlying galactose residues (Williams et al., 2019).

PSA, which targets core fucose, was highly abundant in PLCs/lysosomes, with particular enrichment in their centers. This supports the interpretation that PWA-positive organelles are early endosomes or PLCs rather than endolysosomes, as core-fucosylation is not yet cleaved off therein. Although, to our knowledge, PSA binding to endosomes or lysosomes was not explicitly reported yet, PSA has been widely used as an acrosome marker for the evaluation of sperm morphology (Graham et al., 1990). Given the similarities between acrosomes and lysosomes, such as their enzymatic composition and many other features (Moreno and Alvarado, 2006), it is plausible that PSA also marks certain components of the endolysosomal system.

LCA, another core fucose-binding lectin, showed clustered signal in the center of PLCs/lysosomes and at their periphery (Fig. 3e,f). Interestingly, LCA coincided with only a small fraction of PSA (Fig. S9d), despite the very similar sugar specificity of these lectins (Table S1). The low overlap between LCA and PSA signal could have the following explanations: (1) In contrast to PSA, LCA does not tolerate additional fucosylation apart from core fucosylation, suggesting that PSA mainly bound multi-fucosylated glycans; (2) LCA binds to the PLCs/lysosomes because of its mannose rather than fucose specificity (Bojar et al., 2022). Besides their shared binding to PLCs/lysosomes, these lectins differ in cellular distribution, with PSA additionally strongly associating with the Golgi apparatus and synaptic membranes, supporting the distinct sugar-binding preferences of LCA and PSA.

In principal cell somata in brain tissue, POL and Con A showed a highly enriched signal in some PLCs/lysosomes (Fig. 3d, Fig. S9d,$e_i$), while in other PWA-positive organelles, POL and Con A showed minimal luminal signal and instead were confined to large domains clasping the PLCs/lysosomes (Fig. S9$e_{ii}$). Similarly, in U2OS cells, we observed Con A-rich domains at the periphery of PLCs/lysosomes, where they correlated with the distribution of the early endosome marker EEA1 (Fig. 10a). One plausible explanation is that these cup-shaped domains represent areas of high-mannose glycan recycling back to the ER. The proximity of Con A signal to the early endosome marker EEA1 in U2OS cells supports this idea, as early endosomes are known to play a role in glycan recycling pathways. Alternatively, these structures could reflect the delivery of mannose-6-phosphate-tagged enzymes to lysosomes. While Con A and POL are not expected to bind mannose-6-phosphate moieties directly, they may interact with non-phosphorylated portions of the modified glycans. Lastly, the cup-shaped structures could represent phagophores, given their morphology and their potential to engulf lysosomes for lysophagy. Their ER-like glycan signature aligns with evidence that phagophores predominantly originate from the ER, further supporting this possibility.

WGA was abundantly localized both in the PLC/lysosome periphery and interior. As WGA preferentially binds terminal β-linked GlcNAc, it could target GlcNAc in LacNAc structures exposed after sialic acid and galactose removal or core GlcNAc residues that are revealed after mannose removal. However, not only degradation products carry terminal GlcNAc, also non-digested

GlcNAc-terminated N- and O-glycans are targeted by WGA and thus could explain the peripheral staining observed. At the lysosomal periphery, WGA may bind to the glycoprotein coat on the luminal side, contributed by lysosomal proteins such as LAMPs, but also to peripheral glycans conferred by newly arrived transport vesicles, or cargoes sorted to the periphery for recycling. In addition, WGA binding to some glycosylated surface proteins of PLCs/lysosomes cannot be excluded. Early electron microscopy studies confirmed the binding of WGA to vesicular organelles, including multivesicular bodies in myeloma cells (Tartakoff and Vassalli, 1983) and lysosomes of the *Paramecium multimicronucleatum* (Allen et al., 1989), where it was found at lysosomal membranes and lumens.

Overall, Glyco-STORM experiments did not reveal a clear subdivision into distinct areas with different enzymatic activities, at least with the resolution limit of the method used. Apart from the LCA signal clusters that were yet uniformly distributed throughout PLCs/lysosomes, we did not observe systematically organized subdomains or large clusters with specific degradation products, nor did we detect a degradation gradient from the center to the periphery of the lysosomes. These findings suggest that, within the resolution limits of this study, the degradation process in endosomes and lysosomes is overall uniformly distributed without clear microdomains of enzymatic activity, although nanodomains like those observed for LCA are conceivable. Detailed analyses on PLCs and lysosomes at precisely defined stages of progression are required to accurately determine the distribution of degradation processes within these organelles.

*Clathrin decoration of endosomes/lysosomes*

Many organelles of the endolysosomal system are decorated by clathrin-coated bulges (Fig. 3d,$g_{i,ii}$). These structures could be TGN-derived enzymes that are delivered by a clathrin-mediated mechanism or vesicles emerging from the PLC/lysosome for recycling processes. Given the rapid removal of clathrin coats during transport shortly after vesicle formation, it is more plausible that these bulges are nascent recycling vesicles rather than newly fused TGN vesicles. In most cases, these evaginations showed WGA staining but lacked staining from other lectins, including PSA and Con A. In the context of recycling vesicles containing mannose-6-phosphate receptors, WGA may bind specifically to their terminal carbohydrate moieties. However, while mannose-6-phosphate receptors are known to be glycosylated at multiple sites, the exact glycan structures present on these receptors are not well characterized, and their affinity for WGA remains speculative. Alternatively, other cargo sorting processes would be conceivable.

Interestingly, on the PLC/lysosome surface, the clathrin coat appears to only partially cover the presumed protrusions of recycling vesicles (Fig. 3g$_{i,ii}$). Indeed, it has been proposed that clathrin may initiate budding from lysosomes, after which the initial bud is extended through tubulation. In this scenario, the clathrin coat could remain confined to the tip of the protrusion. Such tubular extrusions might be cleaved either at the tip to form a vesicle or at the neck, producing a tubular intermediate (Saffi and Botelho, 2019). The latter case could explain the polar distribution of clathrin and WGA in areas disconnected from large organelles (Fig. 3g$_{iii,iv}$). Extended tubules with clathrin at their tips have similarly been proposed in budding processes from the TGN or during retromer formation from endosomes (Anitei et al., 2010; Bonifacino and Hurley, 2008). While the idea of polarized clathrin distribution on protrusions remains speculative, Glyco-STORM could further elucidate the morphological nature of vesiculation and tubulation at the endolysosomal system.

#### 4.5 Synaptic glycosylation

Synapse formation, maintenance, plasticity, and transmission are highly organized processes that involve adhesion molecules and intricate protein machineries responsible for synaptic vesicle (SV) docking at the active zone (AZ), neurotransmitter release, and binding to receptors at the postsynaptic density (PSD). While structure-function relationships of many synaptic proteins are well understood, there are remaining gaps. Some of these gaps may be due to a limited exploration of synaptic glycosylation and its potential mechanistic contribution to synaptic function. Despite the known relevance of glycosylation for cell adhesion, endocytosis, and signal transduction, the unique glycome of the central nervous system is understudied. This is particularly significant, given that mutations affecting glycosylation are linked to numerous neurological diseases. Using Glyco-STORM, the precise distribution of glycan motifs at individual molecules and their specific functions can be effectively analyzed. In this study, we utilized lectins to map the glycosylation landscape at synaptic specializations.

*Glycosylation at the synaptic membrane*

WGA and PSA showed most pronounced staining at plasma membranes, i.e., the membranes surrounding the presynaptic calyx of Held and the soma of the postsynaptic principal cells. This observation is consistent with previous reports of WGA staining the glycocalyx and the extracellular space of different cells (Barker et al., 2004; Klevanski et al., 2024; Xie et al., 2020) and neuronal tissue (Klevanski et al., 2020; Michalska et al., 2024). PSA was also previously used to evaluate glycosylation profiles of surfaces of various cells (Rueda et al., 2003; Xie et al., 2020). Diffraction-limited microscopy fails to distinguish WGA membrane binding from intracellular targets, such as endosomes, whereas Glyco-STORM clearly showed WGA binding to synapses at the nanoscale. First, we encountered a signal enrichment of WGA and PSA at synaptic contacts both in calyx of Held synapses (Fig. 4a-c) and cultured hippocampal neurons (Fig. 4d-i). Second, at the calyx of Held synaptic membranes, WGA and PSA showed overall overlapping staining, with both lectins enriched at similar distances from the AZ and PSD (Fig. 4g,h; Fig. S12). Third, apart from the AZ/PSD interface, also other membrane stretches were positive for WGA and PSA. As WGA and PSA signal likely originate from more complex or mature glycans capped by sialic acid or fucose, these lectins allow to study, how mature and immature glycans are distributed along the synaptic membranes. In the following paragraphs, we discuss possible glycan candidates that could be bound by these lectins.

Among the carbohydrate-carrying structures at the neuronal membrane and within the extracellular matrix (ECM), a diverse array of molecules is present. At the neuronal glycocalyx, there are numerous glycoproteins and glycolipids. Glycoproteins comprise neuronal cell adhesion molecules, such as NCAMs, Neurexins, Cadherins, Integrins, and L1, neurotransmitter receptors (e.g., those for AMPA and NMDA), neurotrophic factor receptors, ion channels (e.g., calcium channels, voltage-gated sodium. and potassium channels), and transporters (e.g., glutamate and GABA transporters). Most of these glycoproteins, e.g., neurotransmitter receptors, calcium channels and Neurexins, are specifically enriched at synaptic contacts. Interestingly, in hippocampal cells, Neurexin demonstrated discrete signal-dense nanoclusters, typically one per synaptic contact (Trotter et al., 2019). In contrast, Cadherins have been described as surrounding synaptic contact sites (Uchida et al., 1996). Since at synaptic contacts WGA and PSA binding was mainly confined to the Homer-/Bassoon-occupied area, and showed no distinct clusters neither within nor adjacent to this area, we do not attribute binding of WGA and PSA to Neurexins or Cadherins. However, as long membrane patches that were negative for AZ or PSD markers showed high and contiguous WGA and PSA signals (right head of double-headed arrow in Fig. 4$c_{iii}$), we consider broadly distributed glycoproteins, like NCAM or L1, or ion channels as well as glycolipids, as potential binding sites. For example, at NCAM, WGA could recognize the terminal sialic acid moieties at the poly-sialic acid chains carried by NCAM (Sytnyk et al., 2021). Despite the reported prevalence of high-mannose and bisected glycans in neurons (Williams et al., 2022), synaptic membranes were abundantly bound by PSA, whose binding is inhibited by bisecting GlcNAc, suggesting that core-fucosylated non-bisecting glycans are highly present at these sites.

Furthermore, high-mannose-binding lectins like Con A and POL showed no pronounced staining at the synaptic cleft.

Apart from glycoproteins, also glycolipids could be attractive targets for WGA as sialic acid residues are abundant in the sugar motifs of gangliosides. Beyond membrane-bound carbohydrates, the ECM presents additional potential targets for lectins. While long-chain carbohydrates like chondroitin sulfate proteoglycans and hyaluronic acid are less likely to be recognized by WGA, which favors terminal sugar moieties, and are not expected to bind PSA, which targets core-fucosylated N-glycans, glycoproteins such as laminin and fibronectin within the matrix are more promising candidates for lectin binding.

While at the synaptic membrane between the presynaptic calyx of Held and the postsynaptic principal cell, WGA and PSA signals were confined to a thin layer, the signal at the outer calyx membrane was considerably broader, reinforcing the likelihood of lectin binding to surrounding ECM components or glial cells surrounding the calyx terminal.

*Glycosylation at the active zone*

WGA and PSA were enriched at the synaptic contact sites, showing a comparable staining pattern with similar distribution. In the side view, both lectins peaked at the center of the synaptic cleft, indicating a positional overlap of their targets. Considering an estimated *d*STORM localization precision of approximately 27 nm (pooled for all types of labels) and the maximal label size of 8 nm for PSA and 7 nm for WGA, we cannot distinguish to which extent the signal results from the contribution of pre-, postsynaptic, or ECM glycans. Since PSA is not expected to bind gangliosides or glycosaminoglycans, it is likely that the majority of areas that show overlapping WGA/PSA signal are attributable to glycoproteins. Indeed, most AZ- and PSD-residing receptors, channels and adhesion molecules are glycosylated. For example, NMDA receptors are heavily glycosylated with up to 22 glycosites and some of these carbohydrate modifications are proposed to modulate their integrity and function (Sinitskiy et al., 2017). AMPA receptors and the metabotropic glutamate receptor (mGluR) show multiple N-linked glycosites (Hollmann et al., 1994; Hullebroeck and Hampson, 1992; Nasrallah et al., 2018). Although the precise role of receptor glycosylation is still under investigation, it is increasingly recognized for its critical role in proper receptor trafficking, surface expression, and synaptic plasticity.

Moreover, a recent study reported that in rat synaptosomes brief depolymerization resulted in dynamic changes in sialylation at many glycoproteins residing at transmission sites and propose N-linked sialylation as a modulator of the neurotransmission (Boll et al., 2020). Among the candidates with altered sialylation upon stimulation were NMDA receptors, calcium channels, and numerous cell adhesion molecules. To this day, there are no instruments for targeted, site-specific ablation of carbohydrate motifs at specific proteins. Therefore, precise mapping of these motifs to individual proteins using Glyco-STORM super-resolution imaging offers a unique opportunity to study their function in a spatially related contex. This approach can be further enhanced by comparing stimulated with non-stimulated neurons or different states achieved through other methods, such as drug-treatment.

Interestingly, we observed a spatial anti-colocalization between WGA or PSA and Homer1b/c, and to a lower extent with Bassoon (Fig. 4i). Although Homer1b/c and Bassoon showed a minor correlation to each other, their combined signal was inversely correlated to WGA or PSA (Fig. 4j). This supports the hypothesis for a columnar organizational principle at synaptic contacts (Tang et al., 2016). Although our Glyco-STORM experiments did not include the established nanocolumn constituents, several hypotheses on how the lectin-staining could be related to synaptic nanocolumns can be proposed: (1) Homer1b/c is a central PSD scaffolding protein that via interactions with Shank can bridge metabotropic glutamate receptors and AMPA or NMDA receptors to other PSD components (de Bartolomeis et al., 2022; Samojedny et al., 2022). The bridging nature of Homer could explain its anti-colocalization to the presumable glycan blue-print of neurotransmitter receptors. Alternatively, geometrical arrangement of Homer beneath these receptors might result in antibody binding perpendicularly to the receptor-Homer column. In that case, the positional shift could be due to the label-epitope distance for the primary/secondary antibody complex. (2) As lectins, such as WGA and PSA also bind sugar moieties of proteinaceous ECM components, staining could also reflect ECM nanodomains that may guide neurotransmitter diffusion across the synaptic cleft. This opens up another interesting line of research where co-staining with antibodies against AZ-/PSD-specific receptors, channels and ECM components will allow to precisely map and characterize the glycosylation landscape of the synapse and its functional relevance.

*Lectins targeting synaptic vesicles*

SVs are glycoprotein carriers. At the SV exterior, synapsin I was reported to carry both N-glycans with oligomannose and O-glycans with Lewis$^{X}$ (Galβ1-4(Fucα1-3)GlcNAc) motifs (Sytnyk et al., 2021; Wang et al., 2011). In the lumen of SVs, integral SV proteins, including SV2, Synaptophysin, and Synaptotagmin, feature glycosylation (Abad-Rodríguez and Díez-Revuelta, 2015). SV2 is one of the SV proteins with the highest number of glycosylation sites. A study that investigated botulinum neurotoxins binding to SV2 isoforms revealed core-fucosylated N-glycans as well as sialic acid-modified glycans as co-determinants for the toxin binding (Liu et al., 2023; Yao et al., 2016). Therefore, the binding of WGA and PSA in the SV-occupied area could be conferred by the luminal glycocalyx established by SV2 and other SV proteins. The fact that the lectin signal in the SV cloud was more abundant after multiple rounds of elution using the detergent-containing buffer (Fig. 4c; e.g., Experiment 20 in Table S8) suggests that extensive permeabilization could promote lectin penetration into the SV lumen.

While WGA and PSA, are less expected to target the glyco-modifications of synapsin I, LTL could bind synapsin I via its Lewis$^X$ motif. Mannose-binding lectins, including Con A, LCA, VEA, and POL could recognize oligomannose at synapsin I. Indeed, especially for LCA and VEA (not shown) we observed signal in the presynaptic calyx compartments, however, more quantifications and experiments are necessary to precisely map the localization of their signals.

*Lectins at tubular structures in subsynaptic areas*

Neurons are highly specialized cells equipped with intricate machinery to meet the demands of synaptic maintenance and plasticity, including the precise delivery of neurotransmitter receptors and lipids to synaptic contact sites. Dendrites are known to harbor local protein-synthesis and modification sites. Over the past decades, significant advances have been made in identifying and characterizing these systems within neuronal dendrites. Notably, alongside mRNA, ribosomes, and ER satellites, small structures performing Golgi-like functions have been discovered near synapses (Kemal et al., 2022; Sutton and Schuman, 2006). These include multi-compartment Golgi outposts, typically marked by cis-Golgi protein GM130, and Golgi satellites, which lack GM130 but are rich in glycosylation enzymes. Golgi satellites are often found in spatial proximity to endoplasmic reticulum exit sites (ERES) and endosomes.

Recent studies have shown that neuronal activity can dynamically influence glycosylation patterns, particularly sialylation, at synaptic sites. Sialylation plays a critical regulatory role in neurotransmission, affecting channel gating, receptor assembly, and stability (Scott and Panin, 2014). For instance, depolarization was observed to alter the sialylation of various synaptic transmembrane proteins (Boll et al., 2020). Further research has elucidated that this process involves the formation and dispersal of Golgi satellites, which are responsible for sialylating locally synthesized membrane and secreted proteins (Govind et al., 2021). Specifically, WGA was shown to be endocytosed by stimulated neurons, colocalizing with sialyl-transferase enzymes at presumptive Golgi satellites. This favors the interpretation that the elongated structures observed near the PSD, associated with endosomes (as indicated by clathrin staining), may indeed represent Golgi satellites (Fig. S11a,b). Larger tubular structures with traces of Golgi markers (Fig. S11a-d) could potentially be multi-compartment Golgi outposts. Future Glyco-STORM experiments, utilizing additional markers specific to Golgi satellites and outposts, could further classify these compartments and confirm WGA and PSA as markers to study depolarization-induced glycosylation at the PSD.

Wang, L., Cummings, R.D., Smith, D.F., Huflejt, M., Campbell, C.T., Gildersleeve, J.C., Gerlach,

J.Q., Kilcoyne, M., Joshi, L., Serna, S., Reichardt, N.C., Pera, N.P., Pieters, R.J., Eng, W., Mahal, L.K., 2014. Cross-platform comparison of glycan microarray formats. Glycobiology 24, 507–517. https://doi.org/10.1093/glycob/cwu019

Wang, S., Cesca, F., Loers, G., Schweizer, M., Buck, F., Benfenati, F., Schachner, M., Kleene, R., 2011. Synapsin I is an oligomannose-carrying glycoprotein, acts as an oligomannose-binding lectin, and promotes neurite outgrowth and neuronal survival when released via glia-derived exosomes. J. Neurosci. 31, 7275–7290. https://doi.org/10.1523/JNEUROSCI.6476-10.2011

Westphal, N., Kleene, R., Lutz, D., Theis, T., Schachner, M., 2016. Polysialic acid enters the cell nucleus attached to a fragment of the neural cell adhesion molecule NCAM to regulate the circadian rhythm in mouse brain. Mol. Cell. Neurosci. 74, 114–127. https://doi.org/10.1016/j.mcn.2016.05.003

Westphal, N., Loers, G., Lutz, D., Theis, T., Kleene, R., Schachner, M., 2017. Generation and intracellular trafficking of a polysialic acid-carrying fragment of the neural cell adhesion molecule NCAM to the cell nucleus. Sci. Rep. 7, 1–13. https://doi.org/10.1038/s41598-017-09468-8

Williams, C., Pazos, R., Royo, F., González, E., Roura-Ferrer, M., Martinez, A., Gamiz, J., Reichardt, N.C., Falcón-Pérez, J.M., 2019. Assessing the role of surface glycans of extracellular vesicles on cellular uptake. Sci. Rep. 9, 1–14. https://doi.org/10.1038/s41598-019-48499-1

Williams, S.E., Noel, M., Lehoux, S., Cetinbas, M., Xavier, R.J., Sadreyev, R.I., Scolnick, E.M., Smoller, J.W., Cummings, R.D., Mealer, R.G., 2022. Mammalian brain glycoproteins exhibit diminished glycan complexity compared to other tissues. Nat. Commun. 13. https://doi.org/10.1038/s41467-021-27781-9

Xie, Y., Sheng, Y., Li, Q., Ju, S., Reyes, J., Lebrilla, C.B., 2020. Determination of the glycoprotein specificity of lectins on cell membranes through oxidative proteomics. Chem. Sci. 11, 9501–9512. https://doi.org/10.1039/d0sc04199h

Xu, Z., Ku, X., Tomioka, A., Xie, W., Liang, T., Zou, X., Cui, Y., Sato, T., Kaji, H., Narimatsu, H., Yan, W., Zhang, Y., 2020. O-linked N-acetylgalactosamine modification is present on the tumor suppressor p53. Biochim. Biophys. Acta - Gen. Subj. 1864, 129635. https://doi.org/10.1016/j.bbagen.2020.129635

Yao, G., Zhang, S., Mahrhold, S., Lam, K.H., Stern, D., Bagramyan, K., Perry, K., Kalkum, M., Rummel, A., Dong, M., Jin, R., 2016. N-linked glycosylation of SV2 is required for binding and uptake of botulinum neurotoxin A. Nat. Struct. Mol. Biol. 23, 656–662. https://doi.org/10.1038/nsmb.3245

Zapun, A., Darby, N.J., Tessier, D.C., Michalak, M., Bergeron, J.J.M., Thomas, D.Y., 1998. Enhanced catalysis of ribonuclease B folding by the interaction of calnexin or calreticulin with ERp57. J. Biol. Chem. 273, 6009–6012. https://doi.org/10.1074/jbc.273.11.6009

Zhang, J., Wu, J., Liu, L., Li, J., 2021. The Crucial Role of Demannosylating Asparagine-Linked Glycans in ERADicating Misfolded Glycoproteins in the Endoplasmic Reticulum. Front. Plant Sci. 11, 1–21. https://doi.org/10.3389/fpls.2020.625033
